# EED Maintains the Small Cell Lung Cancer Neuroendocrine Phenotype and Drives Lung Cancer Histological Transformation

**DOI:** 10.1101/2025.07.07.663486

**Authors:** Yixiang Li, Yasmin N. Laimon, Hyeonseo Cho, Marina Vivero, Gabriel Roberti De Oliveira, Andrew Delcea, Varunika Savla, Yuting Chen, Yavuz T. Durmaz, Xintao Qiu, Shweta Kukreja, Rong Li, Talal El Zarif, Wesley Lu, McKayla Van Orden, Jacob E. Berchuck, Roderick T. Bronson, Shuqiang Li, Hongbin Ji, Katerina Politi, Matthew L. Freedman, Henry W. Long, Sabina Signoretti, Matthew G. Oser

## Abstract

Lung cancer histological subtypes include lung adenocarcinoma (LUAD) and small cell lung cancer (SCLC). While typically distinct, combined LUAD/SCLC histology tumors occur, and LUAD can transform into SCLC as a resistance mechanism to targeted therapies, especially in *EGFR*-Mutant LUADs with *RB1*/*TP53*-inactivation. Although PRC2 complex expression increases during this transformation, its functional role has remained unclear. Using CRISPR-based autochthonous immunocompetent GEMMs, we demonstrate that inactivation of EED, the core PRC2 scaffolding subunit, impairs SCLC tumorigenesis and drives histological transformation from ASCL1-positive SCLC to LUAD through a transient NEUROD1-positive intermediate state. Mechanistically, EED loss de-represses bivalent genes co-marked by H3K27me3 and H3K4me3, including LUAD oncogenic RAS, PI3K, and MAPK pathway genes, to promote transformation to LUAD. Consistently, these same signaling genes are bivalently repressed in human SCLC patient-derived xenograft (PDX) tumors, suggesting a conserved PRC2-dependent mechanism to repress LUAD lineage oncogenic signaling to maintain the SCLC neuroendocrine identity. In a complementary *EGFR*-mutant LUAD GEMM with *RB1/TP53* inactivation, EED was required for LUAD-to-SCLC transformation and distant metastasis upon EGFR withdrawal. These findings identify the PRC2 complex as a key epigenetic enforcer of SCLC neuroendocrine identity and nominate EED inhibition as a potential strategy to block SCLC transformation in high-risk LUAD.

## Introduction

Lung cancer is the leading cause of cancer death worldwide and can be broadly divided into the distinct histological subtypes including non-small cell lung cancer (NSCLC) and small cell lung cancer (SCLC). NSCLC is further subdivided into lung adenocarcinoma (LUAD) and squamous cell carcinoma (SCC) which together account for ∼85% of all lung cancers, whereas SCLC accounts for the remaining ∼15%^1^. Although histological subtypes of lung cancer are often mutually exclusive, patients can present with combined LUAD, SCC, and SCLC histology tumors. This may arise from genetically distinct subclones or from lineage plasticity leading to histological transformation of one histology into another^2,3^.

Lineage plasticity has been recognized in lung and other tumor types as a mechanism of therapeutic resistance, most notably in the context of: 1) Histological transformation of *EGFR*-Mutant LUAD to SCLC, as a mechanism of resistance to EGFR inhibitors^4–8^; 2) Histological transformation of prostate adenocarcinoma (PRAD) to neuroendocrine prostate cancer (NEPC), as a mechanism of resistance to androgen deprivation therapy^3,9^. Both LUAD to SCLC and PRAD to NEPC histological transformation portend poor prognoses with relatively few therapeutic treatment options^3^.

Although there have been several correlative studies identifying signatures associated with histological transformation from LUAD to SCLC^5^, causative functional drivers of histological transformation remain largely unknown. Recently, the first genetically-engineered mouse model (GEMM) of *EGFR*-Mutant LUAD to SCLC histological transformation was developed^10^. In this study, highly efficient histological transformation required: 1) Loss of *Rb1* and *Trp53*; 2 tumor suppressors that are nearly universally lost in human SCLC; 2) Overexpression of non-degradable c-Myc; 3) Loss of the *EGFR*-Mutant oncogene^10^. Moreover, inactivation of the tumor suppressor *Pten* leading phosphoinositide 3-kinase (PI3K) pathway activation, when combined with *Rb1* and *Trp53* inactivation and c-Myc overexpression, allowed for SCLC tumorigenesis in alveolar type II cells^10^. This set of genomic alterations required for highly efficient *EGFR*-Mutant LUAD to SCLC histological transformation correlates with alterations that occur in the human counterpart validating the relevance of this GEMM^10^. For example, *RB1* and *TP53* loss near universally occurs in human SCLC^11–13^ and commonly occurs in SCLC transformation^2,4,7,14^. Moreover, co-occurring *RB1* and *TP53* in *EGFR*-Mutant LUAD increase relative risk for SCLC transformation by 43-fold^14^. Alterations that lead to PI3K pathway activation including activation *PIK3CA* and *PTEN* are significantly associated with *EGFR*-Mutant LUAD to SCLC histological transformation in several independent studies^5,7,15^. Together these studies identify genomic alterations that drive LUAD to SCLC histological transformation and allow for the identification of high-risk patients; such as those with concurrent *RB1*, *TP53,* and *PIK3CA* mutations^2,5,7,8,14,15^. However, causative functional drivers that constrain lung cancer histological subtypes and drive LUAD to SCLC histological transformation are not known. There are no current tractable therapeutic strategies to constrain lung cancer histological subtypes and block LUAD to SCLC histological transformation.

SCLC exists broadly in four molecular subtypes: ASCL1, NEUROD1, POU2F3, and Inflammatory^16,17^. However apart from the POU2F3 molecular subtype which often homogenously expresses POU2F3^18^, there is increasing evidence for intra-tumoral heterogeneity of ASCL1 and NEUROD1 subtypes^19^ and for ASCL1 to NEUROD1 subtype plasticity that can be driven by c-MYC^20^ or the epigenetic modifier KDM6A^21^ where c-MYC activation strongly drives further subtype plasticity toward a non-neuroendocrine (non-NE) subtype^20^. Moreover, a recent study identified SCLC subtypes marked by non-negative matrix factorization (NMF) clustering identifying NMF subsets associated with neuroendocrine states and immunotherapy response^12^.

EED is the core scaffolding subunit of the PRC2 complex that canonically deposits tri-methylation marks on histone 3 lysine 27 (H3K27me3) to repress target genes^22,23^. Previous work demonstrated that SCLCs express high levels of EZH2 (the catalytic unit of PRC2) relative to LUADs^24^, and SCLC transformation is associated with signatures of high PRC2 activity^5^. Yet, the role of PRC2 in SCLC tumorigenesis—and whether this complex functionally enforces LUAD-to-SCLC histological transformation, a key mechanism of acquired resistance—remains unknown. Here, we leveraged CRISPR-engineered autochthonous immunocompetent GEMMs to dissect the role of the PRC2 complex in SCLC tumorigenesis and histological transformation, revealing that EED is essential for maintaining the neuroendocrine phenotype and functions as a gatekeeper of lung cancer lineage identity.

## Results

### EED Inactivation Drives Histological Transformation from SCLC to LUAD *In Vivo*

To study the role of EED in SCLC tumorigenesis, we used a CRISPR/Cas9-based, autochthonous, immunocompetent GEMM of SCLC that we developed^21,25–28^ where candidate target genes can be genetically inactivated at tumor initiation together with *Rb1, Trp53,* and *Rbl2* (*RPR2*) and the consequences of target gene inactivation during SCLC tumorigenesis can be studied. We generated adenoviruses that encode: 1) CMV-Cre to activate Cas9 expression; 2) sgRNAs targeting *Rb1, Trp53,* and *Rbl2* (*RPR2*); and 3) an sgRNA targeting *Eed* or a non-targeting sgRNA as a control (**Fig. 1A**). These adenoviruses could simultaneously inactivate all 4 targets and promote Cre expression *in vitro* (**Fig. S1A**). We then intratracheally (IT) injected these adenoviruses into the lungs of lox-stop-lox (LSL) Cas9-P2A-GFP mice for somatic CRISPR-based gene editing to make tumors that were genetically inactivated for *Eed* (*EED*-Mutant *RPR2*) or were *Eed* WT (*EED*-WT *RPR2*) (**Fig. 1A**). In this GEMM, all Cas9-positive tumor cells are GFP-positive. *Eed* was chosen rather than *Ezh2* as EED is the scaffolding protein in all PRC2 complexes and hence eliminates the possibility of paralog compensation from EZH1^29^. Consistent with prior studies nominating EZH2 as a therapeutic target in SCLC^24,30^, *Eed* inactivation significantly delayed tumorigenesis, but tumors did eventually form in most mice injected with the *EED*-Mutant *RPR2* adenovirus (**Fig. 1B-D**). Strikingly, the majority (70%) of (n=26 of 37 tumors) *EED*-Mutant *RPR2* mice had histology consistent with pure LUAD and not SCLC (**Fig. 1C-D, S1B**). Another 27% of mice (n=10 of 37) had both SCLC and LUAD tumors with only 1 mouse showing pure SCLC (**Fig. 1C**). These findings reveal a dramatic phenotypic shift following EED loss.

**Figure 1.**
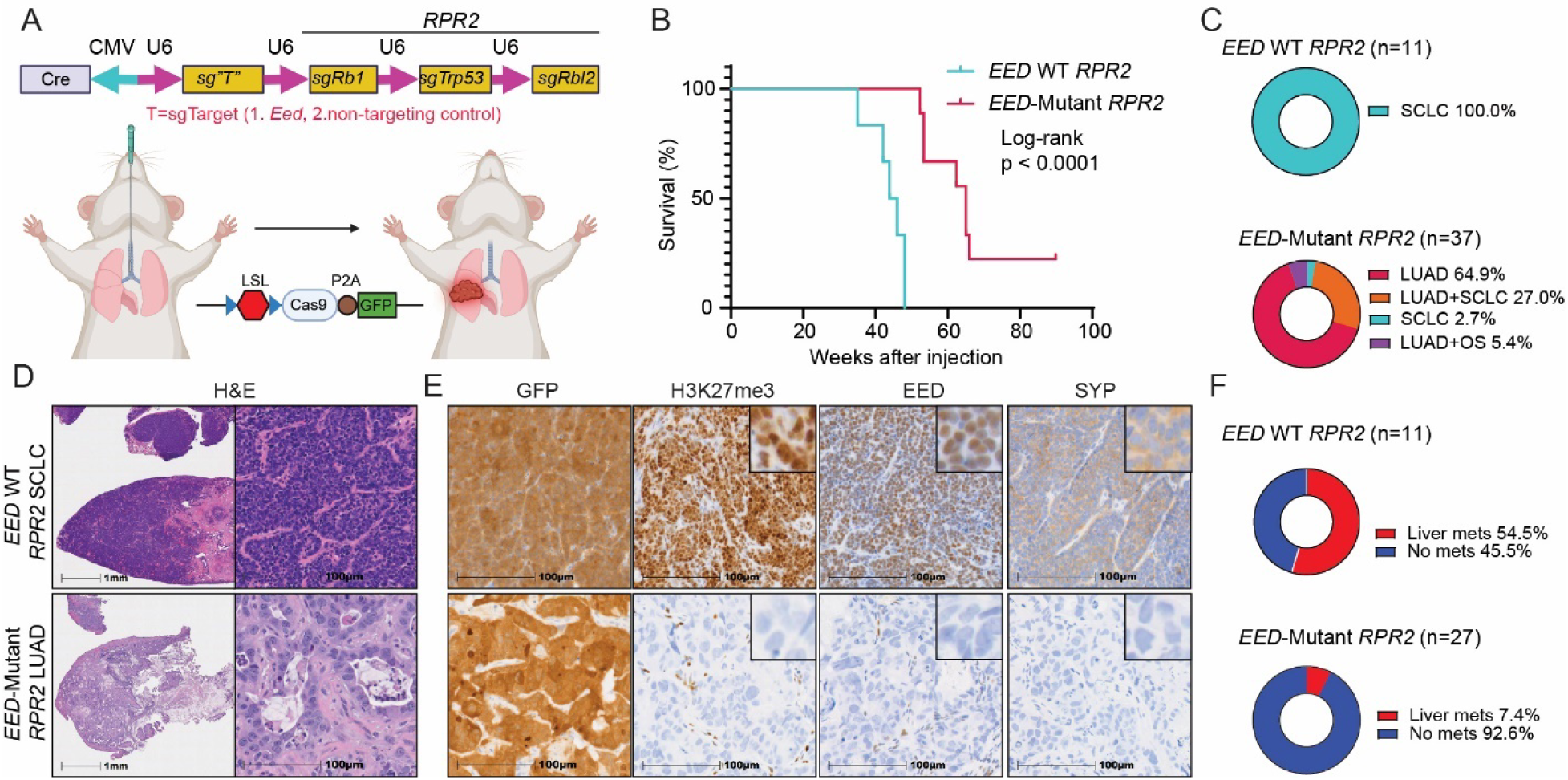
EED Inactivation Drives Histological Transformation from SCLC to LUAD *In Vivo*. **A**. Schematic of the adenovirus used for intratracheal injection (IT) into the lungs of lox-stop-lox (LSL)-Cas9 mice to generate autochthonous SCLC tumors that are *Eed* inactivated (*EED*-Mutant) or *Eed* wild-type (*EED* WT). *RPR2*=sgRb1, sgTrp53, sgRbl2; sg “T”=sg*Eed* or sgControl (non-targeting sgRNA). **B**. Kaplan-Meier survival estimate of LSL-Cas9 mice IT injected with adenoviruses as indicated. Median overall survival: *EED* WT *RPR2*: 44.93 weeks; *EED*-Mutant *RPR2*: 65.00 weeks. N=6 *EED* WT *RPR2* mice, n=9 *EED*-Mutant *RPR2* mice. Log-rank (Mantel-Cox) test was used to calculate p value. **C**. Pie chart of lung cancer histology in *EED*-Mutant *RPR2* tumors (n=37 tumors from 33 independent mice) and *EED* WT *RPR2* tumors (n=11 tumors from 8 independent mice). OS=Osteosarcoma. **D, E**. H&E (**D**), and IHC staining for GFP, H3K27me3, EED, and Synaptophysin (SYP) (**E**) of lung tumors from representative *EED* WT *RPR2* SCLC and *EED*-Mutant *RPR2* LUAD mice. Scale bar is 1 mm (**D**) and 100 microns (**E**). Insets are 9X magnification. (F) Pie chart of liver metastases from *EED*-Mutant *RPR2* (n=27 independent mice) and *EED* WT *RPR2* (n=11 independent mice) mice. See also Figure S1.

Immunohistochemistry (IHC) of tumors confirmed that all tumor cells were GFP-positive, had complete inactivation of EED and its methyltransferase product H3K27me3, and only SCLCs but not LUADs expressed the neuroendocrine marker Synaptophysin (**Fig. 1E, S1C**). CRISPR amplicon sequencing of select tumors similarly confirmed insertion-deletions (indels) in *Eed* and *Rb1* leading to their genetic inactivation (**Fig. S1D-E**). The majority of *EED*-WT *RPR2* SCLCs metastasized to the liver consistent with prior studies using the *RPR2* SCLC model^26,31^ and with SCLC being a highly metastatic tumor^1^ **(Fig. 1F)**. In contrast, *EED*-Mutant *RPR2* LUADs were confined to the lung without any evidence of distant metastasis **(Fig. 1F)**. Together, these data show that EED loss *in vivo* induces a striking histological transformation from SCLC to LUAD, suggesting strong selective pressure for lineage switching as a mechanism of escape from PRC2 loss.

### *EED* Inactivation Epigenetically Restores LUAD Oncogenic Signatures *In Vivo*

To investigate the molecular consequences of EED loss during tumor lineage transformation, we performed RNA-sequencing on primary lung tumors from *EED*-Mutant *RPR2* LUAD and *EED*-WT *RPR2* SCLC mice. As expected, gene set enrichment using MSigDB^32^, ENCODE^33^, and ChEA^34^ revealed that top upregulated genes in *EED*-Mutant tumors were targets normally bound and repressed by PRC2 complex members, including EZH2 and SUZ12 (**Fig. 2A–B**), consistent with functional PRC2 loss. Consistent with differences between human LUAD and human SCLC, *EED*-Mutant *RPR2* LUAD tumors completely lost expression of canonical neuroendocrine (NE) markers including *Chga*, *Insm1*, and *Syp* (**Fig. 2C**) and gained expression of genes involved in receptor tyrosine kinase (RTK) signaling, RAS signaling, and innate immune signaling including genes involved in antigen processing and presentation (**Fig. 2D**). Flow cytometry analysis of lung tumors validated that *EED*-Mutant *RPR2* LUADs restored MHC class I antigen presentation and were significantly more infiltrated with immune and stromal cells (**Fig. S2A-D**). *EED*-Mutant *RPR2* LUAD tumors correlated with both pure and pre-transformed human LUADs, while *EED*-WT *RPR2* SCLCs correlated with both *de novo* and transformed SCLCs (**Fig. 2E**) supporting the translational relevance of this model^5^. Histologic analysis revealed that *EED*-Mutant LUADs displayed mucinous differentiation (**Fig. 2F**), a feature observed in subsets of human LUAD associated with *KRAS* mutations, *NKX2-1* loss, and gastrointestinal lineage gene expression^35–38^. Correspondingly, *EED*-Mutant LUADs upregulated *Cdx2*, *Hnf4A*, and *Pdx1* (**Fig. 2G**), and were enriched for transcriptional signatures of gastrointestinal development (**Fig. 2H**) and mucinous LUADs (**Fig. 2I**).

**Figure 2.**
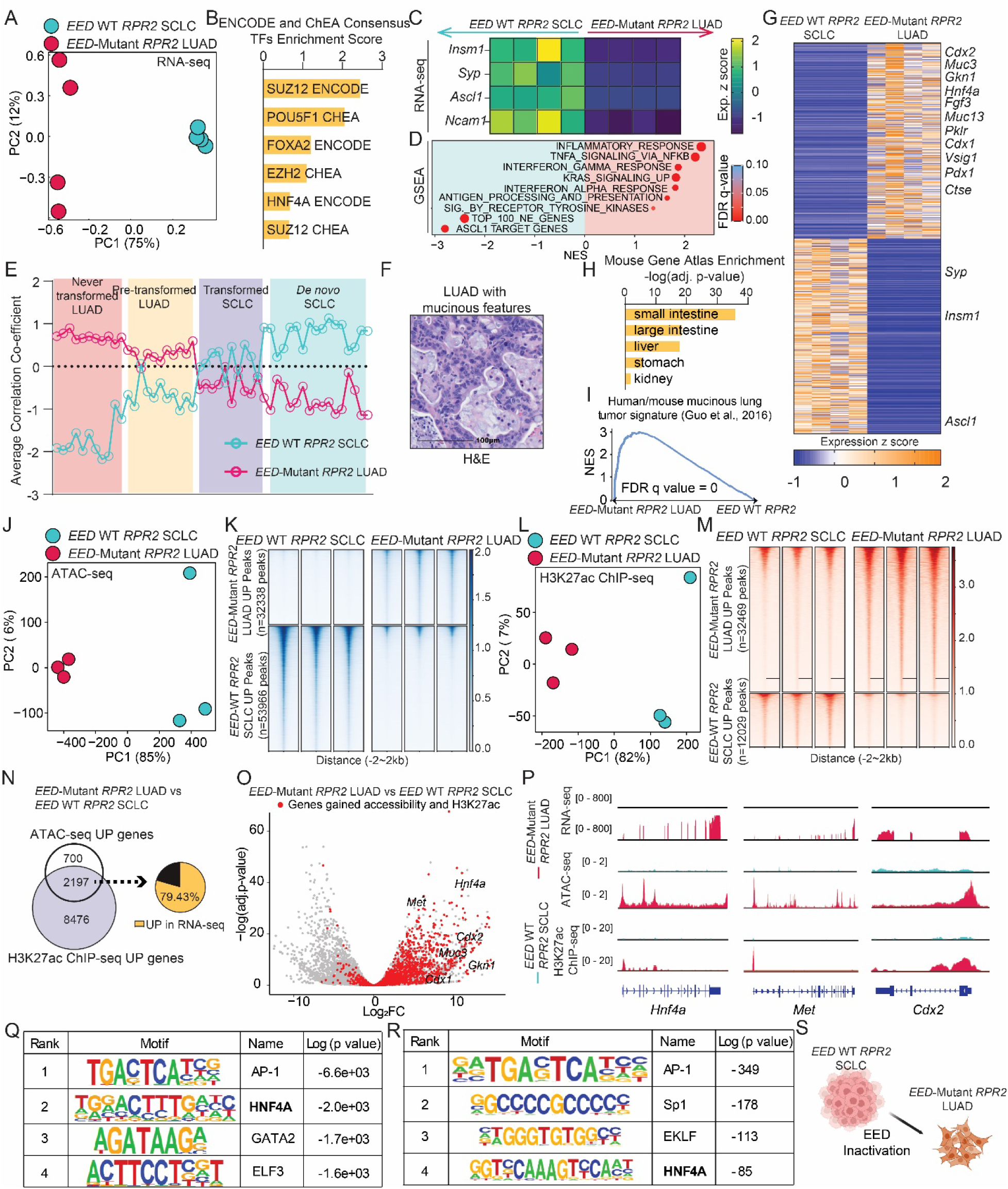
*EED* Inactivation Epigenetically Restores LUAD Oncogenic Signatures *In Vivo*. **A**. Principal component analysis (PCA) plot of RNA-seq data from 4 independent *EED*-WT *RPR2* lung SCLC tumors (blue) and 4 independent *EED*-Mutant *RPR2* LUAD lung tumors (red). **B**. Bar plot of ENCODE and ChEA Consensus Transcription Factors (TFs) enrichment score from RNA-seq data in A showing top enriched TFs whose targets were enriched *EED*-Mutant *RPR2* LUAD vs. *EED*-WT *RPR2* SCLC tumors. TFs with adjusted p value <0.05 were included. **C**. Heatmap depicting z scores of neuroendocrine marker expression from RNA-seq data in **A**. Dark blue is low, yellow is high. **D**. Dot plot of Gene Set Enrichment Analysis (GSEA) from RNA-seq data in A. FDR q-values are visualized by dot color, red=low, blue=high. NES=normalized enrichment score. Size of dot indicates percentage of genes enriched in the leading edge. **E**. Dot plot of the average correlation co-efficient between transcriptomic profiles of *RPR2* tumors and human lung tumors with different histologies from Quintanol-Villalonga et al. 2021^5^. **F**. Representative H&E image of *EED*-Mutant *RPR2* LUAD with mucinous features. Scale bar=100 microns. **G**. Heatmap of top 500 differentially upregulated and downregulated genes in *EED*-Mutant *RPR2* LUAD vs *EED*-WT *RPR2* SCLC ranked by z-score. Yellow is high, blue is low. **H**. Bar plot of negative log transformed adjusted p values of top organs from the mouse gene atlas whose transcriptomic profile enriched in differentially upregulated genes in *EED*-Mutant *RPR2* LUAD vs. *EED*-WT *RPR2* SCLC tumors with an adjusted p-value <0.05. **I**. Enrichment plot of human/mouse mucinous lung tumor signature from Guo et al. 2017^36^ comparing *EED*-Mutant *RPR2* LUAD vs *EED*-WT *RPR2* SCLC. **J, L**. PCA plot of ATAC-seq (**J**) and H3K27ac ChIP-seq (**L**) from 3 independent *EED*-WT *RPR2* SCLC lung tumors (blue) and 3 independent *EED*-Mutant *RPR2* LUAD lung tumors (red). **K, M**. Heatmaps of differential binding peaks from ATAC-seq (**K**) and H3K27ac ChIP-seq (**M**) from lung tumor tissue in the indicated groups. **N, O**. Schematic (**N**) and volcano plot (**O**) showing differentially upregulated genes that are epigenetically activated (gain both accessibility and H3K27Ac) in *EED*-Mutant *RPR2* LUAD vs. *EED* WT *RPR2* SCLC. log2FC=log2 transformed fold change. **P**. Tracks of averaged RNA-seq, ATAC-seq, and H3K27ac ChIP-seq data across all samples above at *Hnf4a* (left), *Met* (middle), and *Cdx2* (right) from the *EED*-WT *RPR2* SCLC (blue) and *EED*-Mutant *RPR2* LUAD (red) mouse tumors. **Q, R**. HOMER motif analysis showing top enriched motifs in *EED*-Mutant *RPR2* LUAD from ATAC-seq (**Q**) and H3K27ac ChIP-seq (**R**). **S.** Schematic showing tumors escape EED inactivation by switching from SCLC to LUAD histology. Figure was made with BioRender. See also Figure S2.

To define the chromatin-level changes underlying this reprogramming, we performed ATAC-seq and H3K27ac ChIP-seq on *EED*-Mutant LUADs and *EED*-WT SCLCs (**Fig. 2J-M, S2E-F**). Integrated analysis revealed that ∼80% of upregulated genes gained both chromatin accessibility and H3K27 acetylation (**Fig. 2N**), indicating widespread epigenetic activation. These included transcription factors associated with gastrointestinal differentiation such as *Hnf4a* and *Cdx2* and LUAD oncogenes including *Met* (**Fig. 2O-P**). Motifs enriched in *EED*-Mutant *RPR2* LUAD for both chromatin accessibility and H3K27ac also included HNF4A (**Fig. 2Q-R**). Together, these findings indicate that EED loss ultimately drives histological transformation from SCLC to LUAD *in vivo* associated with epigenetic activation of LUAD oncogenic programs and gastrointestinal differentiation factors **(Fig. 2S)**.

### EED Inactivation Promotes Rare NEUROD1-positive SCLC with Early Features of LUAD Oncogenic Signaling

While *EED*-Mutant *RPR2* mice predominantly developed LUAD tumors, a small subset formed tumors with SCLC histology (**Fig. 1C**). These rare *EED*-Mutant *RPR2* SCLCs exhibited distinctive morphology, including giant cells and enlarged nuclei compared to *EED*-WT *RPR2* SCLCs (**Fig. 3A**), and were transcriptionally distinct, with enrichment for EMT, inflammatory response, and pancreatic beta cell signatures (**Fig. 3B–C**). The latter included *Neurod1*, a transcription factor that marks the NEUROD1-high (SCLC-N) molecular subtype of human SCLC^12,16,17^. These tumors showed high expression of *Neurod1* and *Neurod1*-correlated genes, alongside reduced expression of *Ascl1* and its transcriptional program (**Fig. 3D–E**), suggesting that EED inactivation promotes ASCL1 to NEUROD1 subtype switching. Consistently, *EED*-Mutant *RPR2* SCLCs correlated with human SCLC-N gene expression profiles from IMpower133^12^ (**Fig. 3E**).

**Figure 3.**
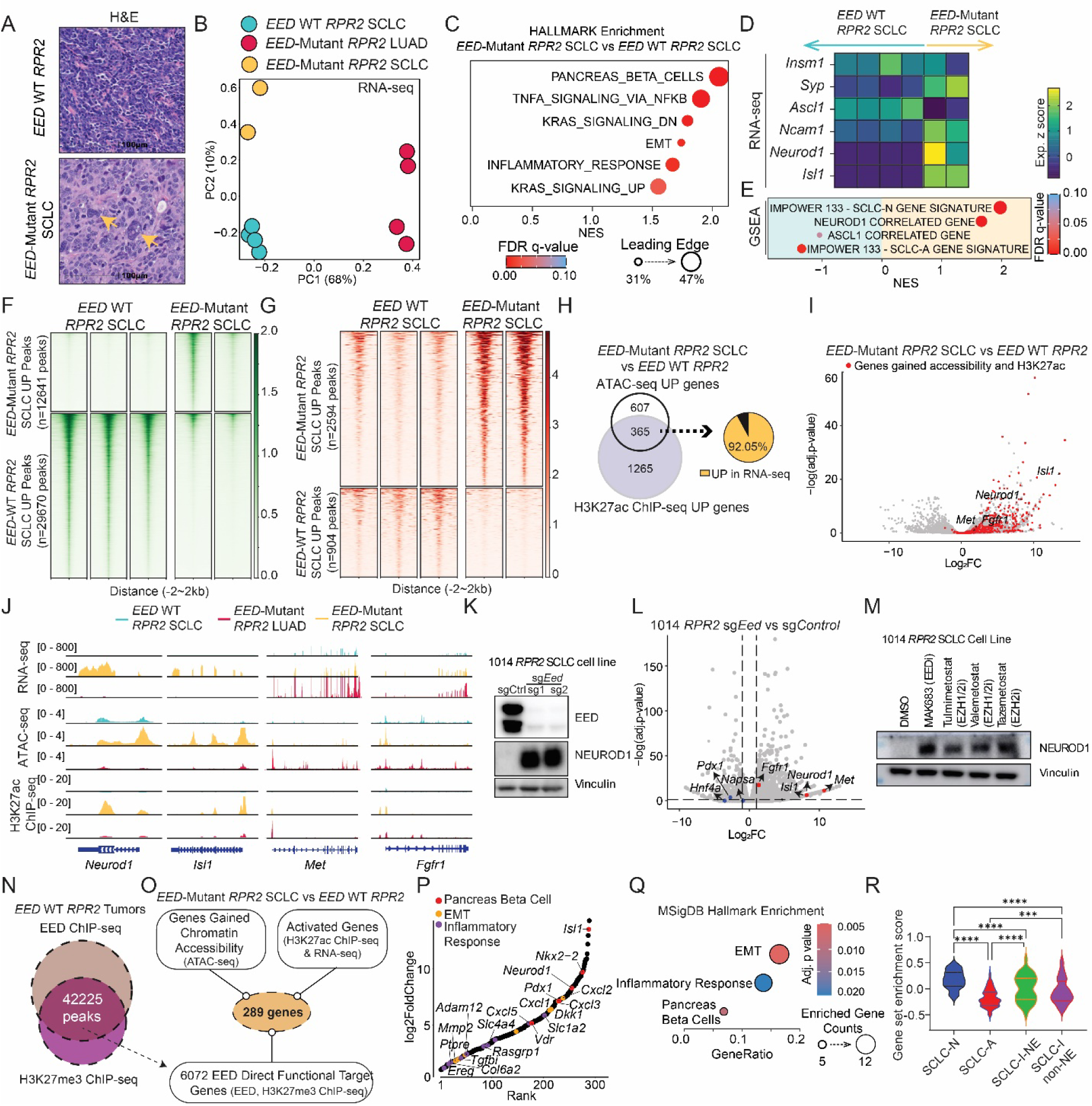
EED Inactivation Promotes Rare NEUROD1-Positive SCLC with Early Features of LUAD Oncogenic Signaling. **A**. Representative H&Es of an *EED*-WT *RPR2* SCLC and an *EED*-Mutant *RPR2* SCLC. Yellow arrows point to giant-cell feature in the *EED*-Mutant *RPR2* SCLC. Scale bar=100 microns. **B**. PCA plot of RNA-seq data from *EED*-WT and *EED*-Mutant *RPR2* SCLC and LUAD tumors. **C**. Dot plot of hallmark pathway GSEA output comparing *EED*-Mutant *RPR2* SCLCs vs. *EED*-WT *RPR2* SCLCs from B. Dot color indicates FDR q-values. Red=low, blue=high. Size of dot indicates percentage of genes enriched in the leading edge. NES=normalized enrichment score. Leading edge was visualized by the size of the dots. EMT=epithelial to mesenchymal transition. **D**. Heatmap showing neuroendocrine marker expression from z-scored RNA-seq data in B. Dark blue is low, yellow is high. **E**. Dot plot of the indicated gene sets from RNA-seq data in B comparing *EED*-Mutant *RPR2* SCLCs vs. *EED*-WT *RPR2* SCLCs. FDR q-values are visualized by dot color, red=low, blue=high. NES=normalized enrichment score. Size of dot indicates percentage of genes enriched in the leading edge. **F,G.** Heatmaps of differential binding peaks from ATAC-seq (**F**) and H3K27ac ChIP-seq (**G**) comparing 2 independent *EED*-Mutant *RPR2* SCLC vs. 3 independent *EED* WT *RPR2* SCLC tumors. **H, I**. Schematic (**H**) and volcano plot (**I**) showing differentially upregulated genes that are epigenetically activated (gain both accessibility and H3K27Ac) in *EED*-Mutant *RPR2* SCLC vs. *EED* WT *RPR2* SCLC. log2FC=log2 transformed fold change. **J**. Tracks of averaged RNA-seq, ATAC-seq, and H3K27ac ChIP-seq data across all samples above from the *EED*-WT SCLC (blue), *EED*-Mutant SCLC (yellow) and *EED*-Mutant LUAD (red) mouse tumors at *Neurod1, Isl1, Met*, and *Fgfr1*. **K, L**. Immunoblot analysis (**K**) and volcano plot of RNA-seq data (**L**) of 1014 *RPR2* EED-WT SCLC cells transduced with sgRNAs targeting *EED* or a non-targeting sgRNA as a control (sgControl). For N, vertical lines indicated fold change of 2 (absolute value). Horizontal line, adjusted p value=0.05. **M**. Immunoblot analysis of 1014 *RPR2* EED-WT SCLC cells treated with the EZH1/2 or EED inhibitors indicated. MAK683 (EED inhibitor)=3 uM, Tulmimetostat (EZH1/2 inhibitor)=100 nM, Valemetostat (EZH1/2 inhibitor)=100 nM, Tazemetostat (EZH2 inhibitor)=5 micromolar. **N**. Schematic of overlap between EED ChIP-seq and H3K27me3 ChIP-seq from *EED* WT *RPR2* SCLC tumors to identify direct EED target genes. **O**. Schematic showing strategy to integrate direct EED target genes with genes reactivated upon EED loss using RNA-seq, ATAC-seq and H3K27ac ChIP-seq to identify direct functional EED targets in *EED*-Mutant SCLC. **P**. Rank plots of RNA-seq log2FoldChange of functional EED targets in *EED*-Mutant SCLC. **Q.** Dot plots of hallmark pathway enrichment analysis of direct functional EED targets in *EED*-Mutant SCLC. EMT: Epithelial to mesenchymal transition. BH adjusted p values were visualized by the dot color, red=low, blue=high. Enriched gene counts were visualized by the size of the dots. **R**. Gene set enrichment score of direct functional EED targets in *EED*-Mutant SCLC (**O**) on different subtypes of human SCLC patient samples RNA-seq from IMPOWER133 (Nabet et al., 2024^12^). SCLC-N=NEUROD1+ SCLC, SCLC-A=ASCL1+ SCLC, SCLC-I-NE=Inflammatory Neuroendocrine SCLC, SCLC-I-non-NE=Inflammatory non-Neuroendocrine SCLC. One way ANOVA test with multiple comparisons was used. *=p<0.05, **=p<0.01, ***=p<0.001, ****=p<0.0001. See also Figure S3.

Similarly, ATAC-seq and H3K27ac ChIP-seq also revealed that top enriched motifs with increased chromatin accessibility in *EED*-Mutant *RPR2* SCLCs were NEUROD1/NEUROG3 and ISL1/2 motifs (**Fig. 3F-G, S3A-G**) with *Neurod1* and *Isl1* gaining both chromatin accessibility and H3K27ac (**Fig. 3H-J**). Interestingly, genes associated with LUAD oncogenic RTK signaling including *Met* and *Fgfr1* were also significantly epigenetically activated in *EED*-Mutant *RPR2* SCLCs relative to *EED*-WT *RPR2* SCLCs (**Fig. 3H-J**). EED genetic inactivation in an ASCL1-positive murine SCLC cell line (1014) similarly upregulated *Neurod1*, *Met*, and *Fgfr1* mRNA (**Fig. 3K–L**). Moreover, EED inactivation or treatment with EZH1/2 or EED inhibitors increased NEUROD1 protein expression (**Fig. 3K,3M**). RNA-seq also revealed an *EED*-Mutant *RPR2* tumor that was positive for POU2F3 which marks a non-NE SCLC subtype^17^ (**Fig. S4A-C**). However, EED inactivation in 1014 cells did not induce POU2F3 expression (**Fig. S4D**) suggesting plasticity toward POU2F3-positive SCLCs after EED inactivation is likely indirect.

Finally, ChIP-seq for EED and H3K27me3 identified 6,072 direct EED target genes, 289 of which were epigenetically activated and expressed in *EED*-Mutant *RPR2* SCLCs (**Fig. 3N–O, Fig. S3H**), including *Neurod1*, *Isl1*, and *Pdx1* (**Fig. 3P**). Gene signatures enriched in these tumors were associated with cell state changes and NEUROD1 subtype switching including epithelial mesenchymal transition (EMT), pancreas beta cells, and the inflammatory response (**Fig. 3Q**). These genes were also highly enriched in the NEUROD1 and inflammatory human SCLC subtypes (**Fig. 3R**) validating relevance of *EED*-Mutant *RPR2* SCLC tumors to the human counterpart. Together, these results show that EED inactivation de-represses NEUROD1 and promotes rare NEUROD1-positive SCLC tumors with early transcriptional activation of LUAD oncogenic pathways, suggesting a transition state on the path to LUAD histological transformation.

### EED Inactivation Promotes SCLC-to-LUAD Transformation Through a NEUROD1-Positive Intermediate Cell State and Requires Cues from the Tumor Immune Microenvironment *In Vivo*

Although EED loss promoted NEUROD1 expression *in vitro* and generated occasional NEUROD1+ SCLC tumors *in vivo*, the vast majority of tumors in *EED*-Mutant *RPR2* mice were LUADs (**Fig. 1C–D**), suggesting selective pressure favoring LUAD fate. Consistently, careful histological examination revealed rare early NE-positive lesions within LUAD-bearing lungs that expressed various SCLC NE subtype transcription factors including ASCL1, NEUROD1, and POU2F3 (**Fig. S4E**). To further explore the relationship of NEUROD1 to lung cancer histological subtypes formed after EED inactivation *in vivo*, we first performed IHC for NEUROD1 and ASCL1 (**Fig. 4A**). Consistent with our results above, *EED*-Mutant *RPR2* SCLC tumors expressed NEUROD1, while *EED*-WT *RPR2* tumors expressed ASCL1 (**Fig. 4A**). Surprisingly, *EED*-Mutant *RPR2* LUADs retained sporadic NEUROD1-positive cells intermixed within tumors (**Fig. 4A-B**). This was a unique feature of *EED*-Mutant *RPR2* LUADs as no NEUROD1-positive cells were found in a *de novo* LUAD *EGFR*-Mutant *RPR2* GEMM (see Methods) nor in *EED*-WT *RPR2* SCLCs (**Fig. 4A-B**). *Neurod1* mRNA expression was also significantly elevated in *EED*-Mutant *RPR2* LUADs relative to *EED*-WT *RPR2* SCLCs (**Fig. 4C**). These data suggest that *EED*-Mutant *RPR2* LUADs may have evolved from NEUROD1-positive *EED*-Mutant *RPR2* SCLCs.

**Figure 4.**
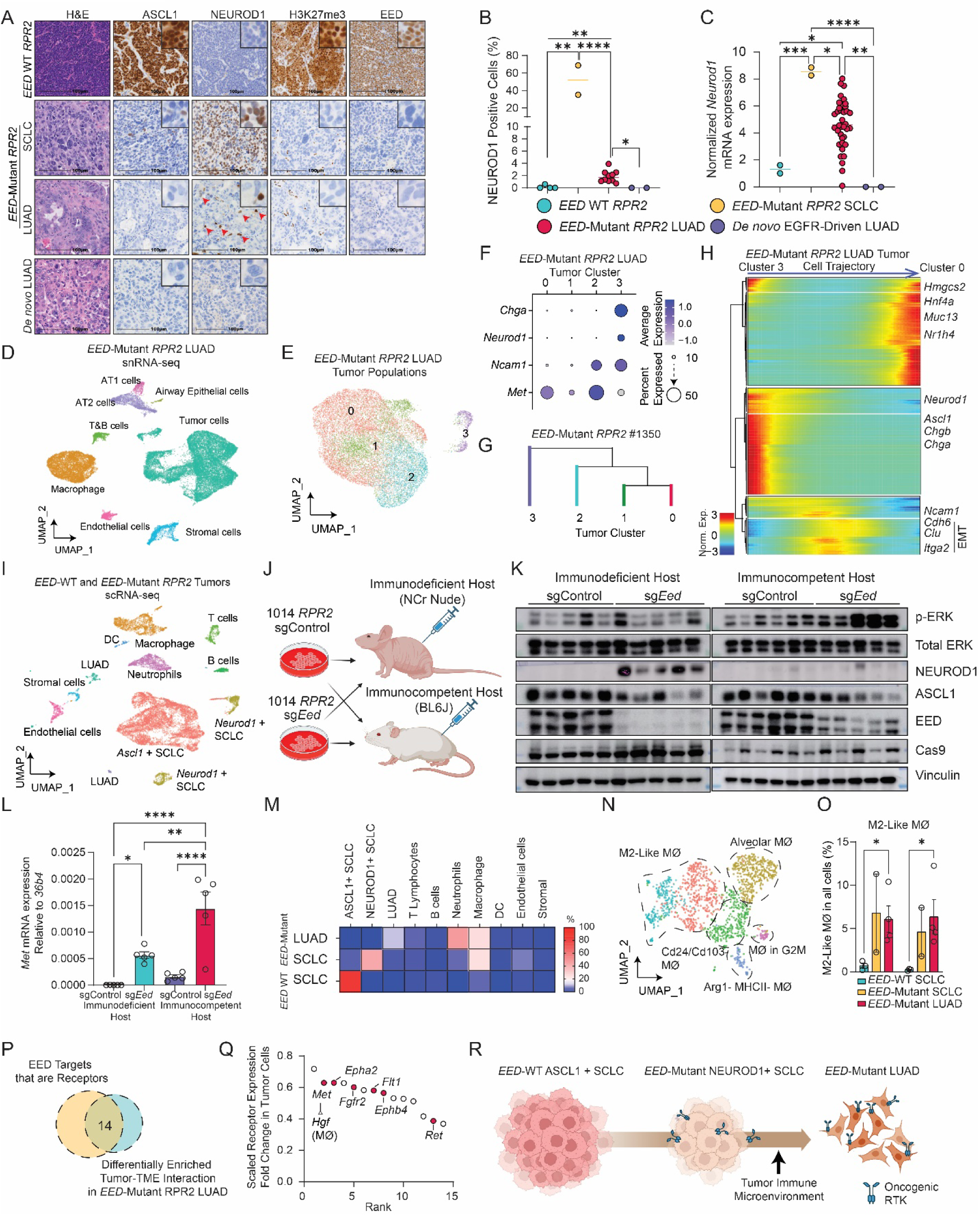
EED Inactivation Promotes SCLC-to-LUAD Transformation Through a NEUROD1-Positive Intermediate Cell State and Requires Cues from the Tumor Immune Microenvironment *In Vivo*. **A**. Representative H&E and IHC staining for ASCL1, NEUROD1, H3K27me3 and EED of lung tumors from GEMMs of *EED*-WT *RPR2* SCLC, *EED*-Mutant *RPR2* SCLC, *EED*-Mutant *RPR2* LUAD, and *de novo* LUAD. Red arrows indicate rare scattered NEUROD1 positive staining. Scale bar=100 microns. Insets are 9X magnification. **B**. Quantitation of NEUROD1-positive cells from NEUROD1 IHC in A. N=4 *EED* WT SCLC *RPR2* tumors, n=2 *EED*-Mutant SCLC *RPR2* tumors, n=10 *EED*-Mutant LUAD *RPR2* tumors, n=2 *de novo* LUAD tumors. **C**. Dot plot of normalized *Neurod1* mRNA expression from RT-qPCR in GEMM tumors indicated. N= 2 *EED* WT SCLC *RPR2* tumors, n=2 *EED*-Mutant SCLC *RPR2* tumors, n=39 *EED*-Mutant LUAD *RPR2* tumors, n=2 *de novo* LUAD tumors. **D, E**. Uniform Manifold Approximation and Projection (UMAP) of all cells (**D**) and tumor cells (**E**) from single-nucleus RNA-seq (snRNA-seq) from 3 *EED*-Mutant *RPR2* LUAD tumors from 3 independent mice. N=27,117 total cells, n=16,590 tumor cells. **F**. Dot plot showing average expression of indicated genes in each tumor cell cluster. Dot size represents percentage of cells expressing the indicated genes. Dot color represents expression level, blue=high, white=low. **G**. Phylogenetic tree of tumor cell clusters from *EED*-Mutant *RPR2* LUAD tumor 1350 from **E** using Maximum Parsimony method. **H**. Heatmap of normalized expression level of top 300 trajectory-defining genes along the *EED*-Mutant *RPR2* LUAD tumor cell trajectory. Blue is low, red is high. **I.** UMAP of all cells from single-cell RNA-seq (scRNA-seq) on 3 independent *EED*-WT SCLC (n=7889 cells), 2 *EED*-Mutant SCLC (n=2064 cells), and 4 *EED*-Mutant LUAD lung tumors (n=2013 cells). Each tumor is from an independent mouse. Mϕ: macrophages. DC: dendritic cell. **J-L**. Schematic (**J**), immunoblot analysis (**K**), and RT-qPCR (**L**) of 1014 *RPR2 EED*-WT SCLC cells transduced with an sgRNA targeting *EED* (sg*Eed*) or a non-targeting sgRNA as a control (sgControl) and then subcutaneously implanted in immunodeficient (NCr nude) or immunocompetent (BL6J) mice. **M**. Heatmap of both tumor and immune cell types indicated comparing *EED*-Mutant LUAD, *EED*-Mutant SCLC, and *EED*-WT SCLC from the scRNA-seq data in **I**. **N,O**. UMAP (**N**) and quantification (**O**) of percentages of two M2-like Mϕ clusters relative to total cells in each sample. **P,Q**. Schematic (**P**) and rank plot (**Q**) of scaled receptor expression fold changes in *EED*-Mutant LUAD tumor cells whose cell-cell interactions were enriched in *EED*-Mutant LUAD tumors when compared to *EED* WT SCLC tumors and receptors upregulated after *EED* genetic inactivation in 1014 *EED*-WT SCLC cell line after *EED* CRISPR-mediated inactivation *in vitro*. For *Met*, the corresponding ligand (*Hgf*) and the tumor microenvironment component with significantly higher expression (Mϕ) is indicated in parenthesis. Mϕ: macrophages. **R**. Schematic showing SCLC to LUAD histological transformation occurs through a NEUROD1-positive transient intermediate state that requires cues from the tumor immune microenvironment. Figure was made with BioRender. *=p<0.05, **=p<0.01, ***=p<0.001, ****=p<0.0001. Student’s t test was used to calculate two-sided p-value. See also Figure S4, S5.

To investigate this possibility, we performed single-nucleus RNA-seq (snRNA-seq) on three independent *EED*-Mutant *RPR2* LUADs. Unsupervised clustering identified four tumor cell clusters (**Fig. 4D–E**). One cluster (cluster 3) expressed NE markers including *Neurod1*, *Chga*, and *Ncam1* and accounted for ∼4% of tumor cells, while other clusters expressed LUAD genes such as *Met* (**Fig. 4F**). To ask whether there was a clonal relationship between cluster 3 *Neurod1*-positive cells with the other tumor cell clusters, we used InferCNV^39^ to estimate copy number variations (CNVs) of each cluster within samples. InferCNV revealed diverse CNV landscapes between samples (**Fig. S5A-C**). However, we observed substantial shared copy number alterations between *Neurod1*-positive cluster 3 tumor cells and tumor cells within other clusters (**Fig. S5A-C**) suggesting tumor cells from the same sample are likely derived from a common clone. *Neurod1*-positive cluster 3 tumor cells showed significantly less CNV events compared to other clusters within the same tumor where tumor cells of other clusters gained or lost CNVs relative to *Neurod1*-positive cluster 3 cells (**Fig. S5D**). Indeed, phylogeny tree construction also supported *Neurod1*-positive cluster 3 tumor cells as the likely ancestor (**Fig. 4G, S5E-F**). Similarly, pseudo-time cell trajectory analysis revealed that first NE genes (*Ascl1*, *Neurod1*, *Ncam1*, *Chgb*) were highly expressed followed by persistence of *Neurod1* and *Ncam1* expression with eventual loss of NE genes and upregulation of LUAD and GI differentiation genes (*Met*, *Hnf4a*, *Nr1h4*) (**Fig. 4H, S5G-H**). Interestingly, EMT genes including *Cdh6, Clu*, and *Itga2,* were also highly expressed during the persistent *Neurod1+* and *Ncam1+* intermediate cell state.

To further validate this trajectory, we performed single-cell RNA-seq (scRNA-seq) on *EED*-Mutant *RPR2* LUADs (n=4) and *EED*-Mutant *RPR2* SCLCs (n=2), comparing them to previously published scRNA-seq data from *EED*-WT *RPR2* SCLCs (n=3)^21^ (**Fig. 4I, S5I**). As expected, *EED*-Mutant SCLCs expressed *Neurod1*, *Isl1*, and other NE markers, while *EED*-Mutant LUADs lost NE features and expressed *Hnf4a* and *Muc13* (**Fig. S5J**). Unsupervised ordering of *EED*-WT and *EED*-Mutant tumor cells predicted a single lineage trajectory of *EED*-WT *RPR2* SCLC (ASCL1-positive) to *EED*-Mutant *RPR2* SCLC (NEUROD1-positive) to *EED*-Mutant *RPR2* LUAD (HNF4A-positive), further supporting NEUROD1-positive SCLC is an intermediate state during histological transformation from SCLC to LUAD (**Fig. S5K-M**).

Despite upregulating *Neurod1* and *Met*, EED inactivation in 1014 cells *in vitro* did not induce LUAD lineage markers (e.g. *Hnf4a*) or complete transformation, suggesting that extrinsic *in vivo* cues from the tumor immune microenvironment are required to drive SCLC-to-LUAD transition (**Fig. 3L**). To test this, we transplanted *EED*-isogenic 1014 cells into immunocompetent (BL6J) and immunodeficient (NCr nude) mice (**Fig. 4J**). In immunodeficient hosts, *EED*-inactivated tumors retained high NEUROD1 expression and failed to activate MAPK signaling. In contrast, *EED*-inactivated tumors in immunocompetent mice lost NEUROD1 and robustly activated MAPK signaling and *Met*, suggesting that the tumor immune microenvironment promotes LUAD fate selection following EED loss (**Fig. 4K-L**).

To explore which immune-derived cues might promote LUAD histological transformation, we analyzed the tumor immune microenvironment (TIME) **(Fig. 4I)**. Consistent with differences between human SCLC and human LUAD, *EED*-WT *RPR2* SCLCs were mostly comprised of tumor cells with relatively few immune cells, while *EED*-Mutant *RPR2* LUADs were significantly more immune cell infiltrated **(Fig. 4M).** Interestingly, *EED*-Mutant *RPR2* SCLCs also had significantly more immune cell infiltration relative to *EED*-WT *RPR2* SCLCs (**Fig. 4M**). Immunosuppressive M2-like macrophages^40,41^ expressing *Msr1*, *Arg1*, and *Tgfb1* were enriched in both *EED*-Mutant LUADs and SCLCs (**Fig. 4N-O, S5N-O**).

To identify potential microenvironmental drivers of transformation, we performed a cell–cell interaction analysis^42^ comparing TIME ligands and tumor cell receptors in *EED*-Mutant LUADs versus *EED*-WT SCLCs (**Fig. 4P-Q**), revealing 58 enriched receptor–ligand pairs. Cross-referencing this list with RNA-seq data from EED-inactivated 1014 cells (**Fig. 3L**) identified 14 upregulated tumor-expressed receptors with corresponding TIME ligands, including LUAD RTK oncogenes such as *Met*, *Ret*, and *Fgfr2* (**Fig. 4P-Q**), all of which converge on RAS signaling. Notably, *Met* was the second-highest expressed receptor, with its canonical ligand *Hgf* highly expressed by macrophages in *EED*-Mutant *RPR2* LUADs. These findings identify NEUROD1-positive tumor cells as a transitional state in the lineage switch from SCLC to LUAD following EED loss, and demonstrate that the tumor immune microenvironment promotes LUAD RTK oncogenic signaling to complete this transformation *in vivo* (**Fig. 4R**).

### EED Directly Binds and Represses RAS, PI3K, and MAPK Pathway LUAD Oncogenic Signaling Genes

To elucidate how EED maintains the ASCL1-positive SCLC neuroendocrine cell state and why its loss drives SCLC-to-LUAD transformation, we performed ChIP-seq for EED and H3K27me3, identifying 6,072 direct EED target genes (**Fig. 3N**). Of these, 831 were epigenetically activated and expressed in *EED*-Mutant *RPR2* LUADs (**Fig. 5A**), including LUAD oncogenes (*Met*) and gastrointestinal lineage genes (*Hnf4a*, *Cdx2*) (**Fig. 5B**). Gene signatures enriched among these activated targets included RAS, PI3K, and MAPK signaling, EMT, and gastric cancer features (**Fig. 5C–D**), with strong overlap in inflammatory subtypes of human SCLC (**Fig. 5E**).

**Figure 5.**
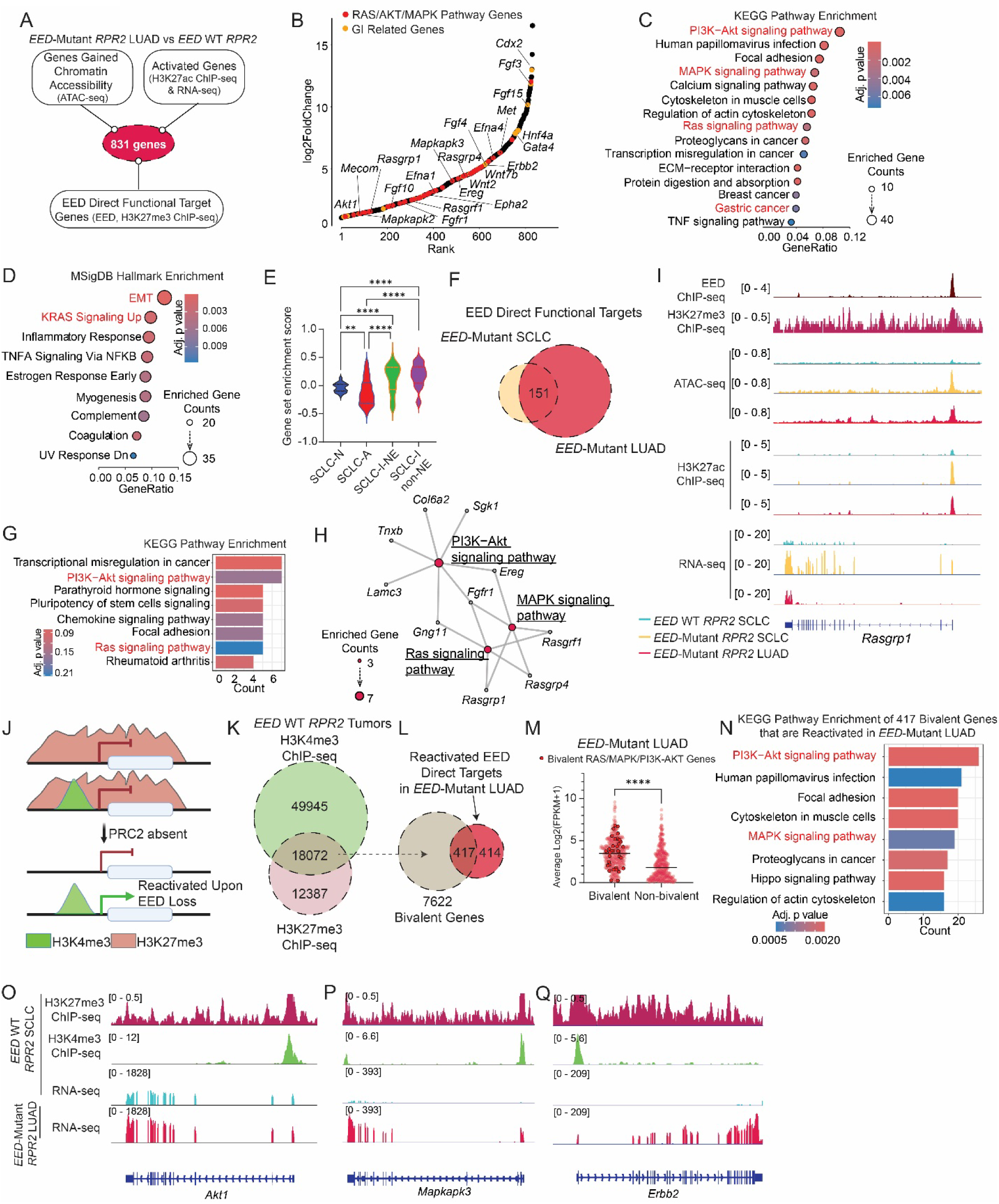
EED Directly Represses Bivalently Marked RAS, PI3K, and MAPK Pathway Genes to Suppress LUAD Oncogenic Signaling and Enforce SCLC Neuroendocrine Identity. **A**. Schematic showing strategy to integrate direct EED target genes with genes reactivated upon EED loss using RNA-seq, ATAC-seq and H3K27ac ChIP-seq to identify direct functional EED targets in *EED*-Mutant LUAD lung tumors. **B**. Rank plots of RNA-seq log2FoldChange of functional EED targets in *EED*-Mutant LUAD lung tumors. **C,D.** Dot plots of KEGG pathway enrichment analysis (**C**) and hallmark pathway enrichment analysis (**D**) of direct functional EED targets in *EED*-Mutant LUAD. EMT: Epithelial to mesenchymal transition. BH adjusted p values were visualized by the dot color, red=low, blue=high. Enriched gene counts were visualized by the size of the dots. **E**. Gene set enrichment score of direct functional EED targets in *EED*-Mutant LUAD on different subtypes of human SCLC patient samples RNA-seq from IMPOWER133 (Nabet et al., 2024^12^). SCLC-N=NEUROD1+ SCLC, SCLC-A=ASCL1+ SCLC, SCLC-I-NE=Inflammatory Neuroendocrine SCLC, SCLC-I-non-NE=Inflammatory non-Neuroendocrine SCLC. One way ANOVA test with multiple comparisons was used. *=p<0.05, **=p<0.01, ***=p<0.001, ****=p<0.0001. **F-H**. Schematic (**F**), KEGG pathway enrichment analysis (**G**), and gene network analysis of enriched Ras, AKT, and MAPK pathways (**H**) of overlapping EED direct functional targets in *EED*-Mutant SCLC and *EED*-Mutant LUAD. Enriched gene counts were visualized by the size of the dots. BH adjusted p values were visualized by the bar color, red=low, blue=high. **I**. Tracks of averaged RNA-seq, ATAC-seq, and H3K27ac ChIP-seq data across all samples from the *EED*-WT SCLC (blue), *EED*-Mutant SCLC (yellow) and *EED*-Mutant LUAD (red) mouse lung tumors and EED (brown) and H3K27me3 (purple) ChIP-seq data from *EED*-WT SCLC at *Rasgrp1.* **J.** Schematic showing genes with bivalent H3K4me3 and H3K27me3 histone marks and primed for reactivation upon PRC2 loss. Figure was made with BioRender. **K,L**. Schematic showing strategy to identify bivalent genes reactivated upon EED loss by overlapping peaks of H3K4me3 and H3K27me3 ChIP-seq data from *EED*-WT *RPR2* SCLC tumors (**K**) and then overlap these bivalent peaks with reactivated EED direct target genes in *EED*-Mutant *RPR2* LUADs (**L**). **M**. Expression of reactivated EED direct targets that are bivalent or not in *EED*-Mutant LUAD. **N**. Bar plot of enriched KEGG pathways of reactivated bivalent EED direct targets in *EED*-Mutant LUAD. BH adjusted p-values are visualized by the bar color, red=low, blue=high. Count: Enriched gene counts. **O-Q**. Tracks of averaged H3K27me3 (red) and H3K4me3 (green) ChIP-seq and RNA-seq at *Akt1* (left), *Mapkapk3* (middle), and *Erbb2* (right) showing bivalent peaks and RNA-seq activation upon EED loss. See also Figure S6.

Only 151 EED target genes were commonly activated in both *EED*-Mutant *RPR2* LUADs and SCLCs (**Fig. 5F**), yet this shared set was significantly enriched for RAS and PI3K-AKT pathway genes (**Fig. 5G-H**). For instance, *RASGRP1*, a RAS guanine nucleotide exchange factor, was a direct EED-repressed gene epigenetically activated in both contexts (**Fig. 5I**). These data suggest that EED directly represses RAS pathway components, and its loss upregulates RAS signaling even in tumors retaining SCLC histology, potentially priming for LUAD transformation.

RAS pathway activation is a conserved hallmark downstream of nearly all LUAD oncogenic RTKs and has been shown to antagonize the NE phenotype in murine SCLC^43^—a finding we validated in human ASCL1-positive SCLC cell lines overexpressing wild-type or mutant (G12C) KRAS (**Fig. S6A–B**). To investigate whether EED loss induces RAS pathway dependence, we derived five cell lines from *EED*-Mutant *RPR2* LUAD tumors (1350, 1339, 1343, 1344, 1345), which, unlike SCLCs, grew as adherent monolayers (**Fig. S6C–D**). These lines exhibited elevated phospho-ERK levels and showed strong sensitivity to the pan-RAS inhibitor RMC-6236^44^ and the MEK inhibitor Trametinib, compared to *EED*-WT *RPR2* SCLC cells (**Fig. S6E–H**), consistent with high RAS signaling and dependency.

To test whether this phenotype was reversible, we re-expressed wild-type EED or a catalytically inactive mutant (EED-inactive) in two *EED-*Mutant LUAD lines (1339 and 1344) and performed RNA-seq (**Fig. S7A–B, S7E**). Re-expression of *EED*-WT, but not *EED*-inactive, downregulated PRC2 (EZH2/SUZ12) target genes, validating on-target activity (**Fig. S7C, S7F**). KEGG enrichment analysis further confirmed suppression of MAPK and RAP1 signaling pathways upon *EED*-WT re-expression (**Fig. S7D, S7G**). Together, these data demonstrate that EED directly represses RAS signaling to maintain the SCLC neuroendocrine phenotype, and that its loss induces RAS activation and LUAD oncogenic signaling, facilitating histological transformation.

### Loss of the PRC2 Complex Initiates SCLC-to-LUAD Histological Transformation Through Epigenetic Activation of Bivalent Genes Enriched in PI3K and MAPK Signaling

While 6,072 genes were marked by both EED and H3K27me3, relatively few were activated upon EED loss—289 in SCLC and 831 in LUAD (**Fig. 3O, 5A**). Bivalent genes, which carry both the activating H3K4me3 and the repressive H3K27me3 histone marks, are known to be selectively de-repressed upon PRC2 inhibition during differentiation^45^. We therefore hypothesized that bivalent genes would be preferentially activated upon EED loss and include key regulators of SCLC-to-LUAD histological transformation (**Fig. 5J**). To identify bivalent genes, we performed ChIP-seq for H3K4me3 in *EED*-WT *RPR2* SCLC tumors and overlaid these peaks with H3K27me3 peaks in the same samples, identifying 7,622 gene promoters with bivalent chromatin marks (**Fig. 5K, S8A–B**). As in previous studies^46,47^, GREAT and hallmark enrichment analyses showed that bivalent genes were associated with MHC class I antigen presentation and EMT (**Fig. S8C–E**). In line with our hypothesis, a large percent of genes activated upon EED loss had bivalent marks at their promoters including 118 of 289 (41%) genes activated upon EED loss in SCLC and 417 of 831 (∼50%) genes activated upon EED loss in LUAD (**Fig. 5L, S8F**). Consistent with genes marked by bivalency being poised for activation upon H3K27me3 loss^45^, reactivated EED direct targets in both *EED*-Mutant SCLC and LUAD that are bivalent have significantly higher expression than those that are not bivalent (**Fig. 5M, S8G**). Strikingly, bivalent genes activated in LUADs following EED loss were significantly enriched in PI3K-Akt and MAPK signaling pathways including *Akt1*, *Mapkapk3*, as well as upstream growth factor receptors (e.g. *Erbb2*) (**Fig. 5M-Q**). These findings indicate that EED directly represses genes that promote RAS pathway activation and that these genes are bivalent and poised for activation—thereby rapidly initiating SCLC-to-LUAD histological transformation upon PRC2 inactivation.

On the other hand, *Neurod1* is a bivalent target reactivated in *EED*-Mutant *RPR*2 SCLC and among the top highly expressed genes (**Fig. S8F-H**). Some reactivated EED direct targets were not bivalent including the gastrointestinal lineage genes *Hnf4A* and *Cdx1* (**Fig. S8F-H**) suggesting epigenetic activation of these genes requires further selection *in vivo.* This is consistent with our RNA-seq data in 1014 cells as *Hnf4 and Cdx1* were not upregulated upon EED inactivation *in vitro* (**Fig. 3L**). Taken together, these findings suggest that bivalency marked by H3K27me3 and H3K4me3 poises RAS/PI3K/MAPK pathway genes and *Neurod1* for activation, all of which are robustly induced upon EED loss.

### EED Inactivation Blocks LUAD to SCLC Transformation in an *EGFR*-Mutant GEMM as a Mechanism of Resistance to EGFR Inactivation

Our earlier data showed that EED is required to maintain ASCL1-positive SCLC histology in the *RPR2* GEMM, with EED inactivation promoting lineage infidelity and histological transformation from SCLC to LUAD (**Fig. 1**). Based on this, we hypothesized that EED/PRC2 activity may also be required for the acquisition of SCLC histology during LUAD-to-SCLC transformation in the setting of resistance to EGFR inhibition, a process observed in *EGFR*-Mutant LUADs harboring *RB1* and *TP53* mutations^14^. To test this, we developed an *EGFR*-Mutant LUAD GEMM capable of histological transformation. We used a doxycycline (Dox)-inducible EGFR L858R transgenic model^48,49^ crossed with: 1) *Trp53* (P) flox/flox mice; 2) lox-stop-lox (LSL) rtTA mice to activate the Dox inducible promoter; and 3) LSL-Cas9-P2A-GFP mice (**Fig. 6A**). We then generated adenoviruses encoding: 1) Cre to activate Cas9 and rtTA expression; 2) sgRNAs targeting *Rb1* (R). We also included an sgRNA targeting either *EED* (E) or a non-targeting sgRNA as a control (C) to ultimately make *EGFR*-Mutant LUAD GEMMs with concurrent *Rb1* and *Trp53* inactivation with EED inactivated (referred to hereafter as *EGFR*-Mutant *ERP*) or where *EED* was WT (referred to hereafter as *EGFR*-Mutant *CRP*). Immunoblot analysis validated that these adenoviruses induced Cre expression and efficiently inactivated RB1 and EED (**Fig. S9A**).

**Figure 6.**
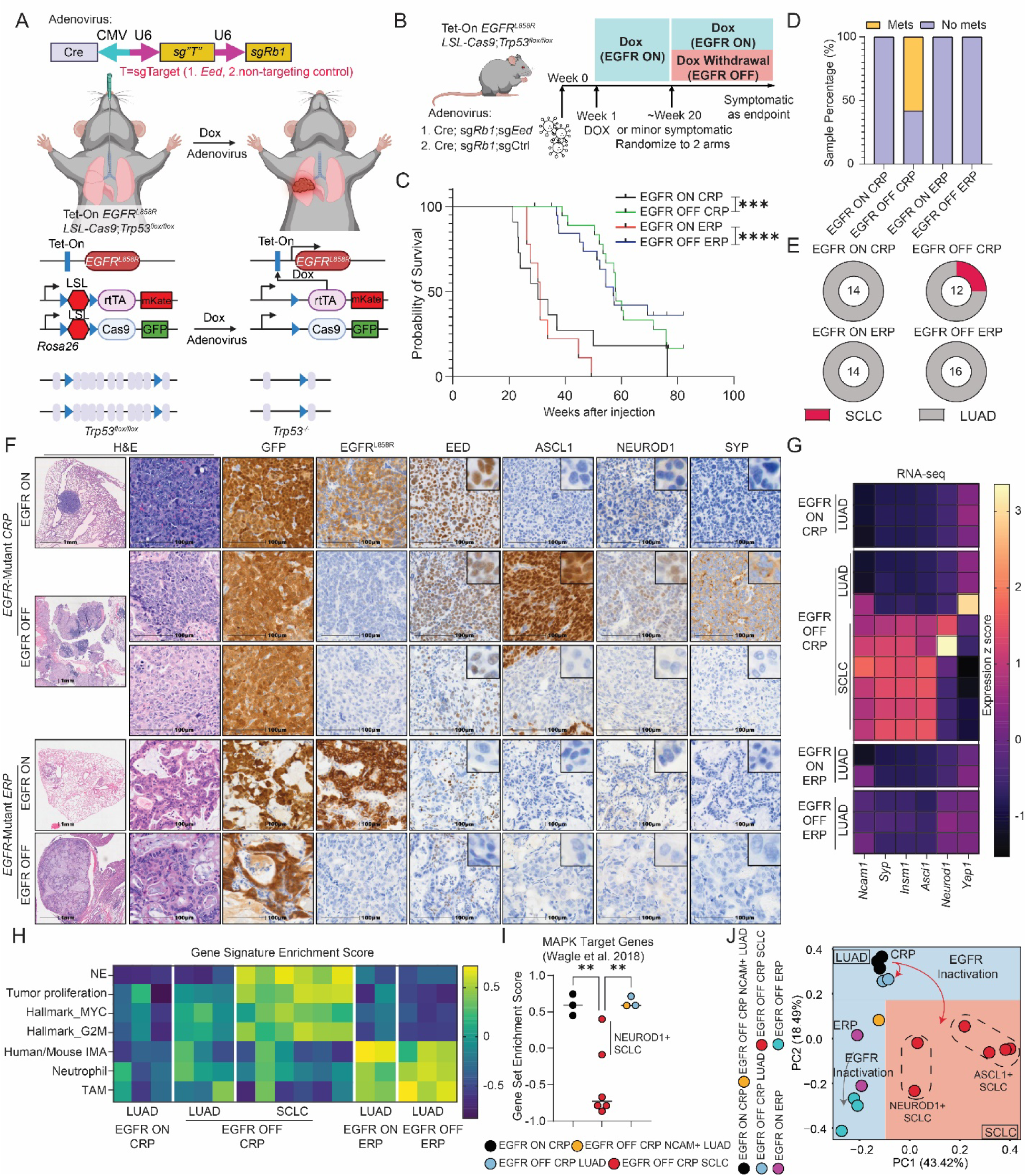
EED Inactivation Blocks LUAD to SCLC Transformation as a Mechanism of Resistance to EGFR Inactivation in an *EGFR*-Mutant GEMM. **A.** Schematic of the CRISPR-based *EGFR*-Mutant LUAD GEMM with concurrent *Rb1* and *Trp53* inactivation made isogenic to *Eed* by including an sgT to target *Eed* to make ERP tumors or by including a non-targeting sgRNA as a control to make CRP tumors. Dox=doxycycline. LSL=lox-stop-lox. **B**. Schematic of adenovirus injection and Dox induction and withdrawal timeline. **C**. Kaplan-Meier survival estimate of mice IT injected with adenoviruses indicated and then treatment as indicated in (B). Log-rank p-value was used to calculate p-values. **D**. Bar plot of percent of mice with metastases present in each group. n=14 EGFR ON CRP independent mice, 12 EGFR OFF CRP independent mice, 14 EGFR ON ERP independent mice, and 16 EGFR OFF ERP independent mice. **E-F**. Pie charts of histology (**E**) with representative H&E and IHC staining for GFP, EGFPL858R, EED, ASCL1, NEUROD1 and SYP (**F**) of lung tumors from the groups indicated. Scale bar (1^st^ column) =1 mm. Scale bar (2^nd^-8^th^ columns) =100 microns. Insets are 9X magnification. **G**. Heatmaps of z-scored gene expression of SCLC NE and non-NE markers from RNA-seq data of the *EGFR*-Mutant lung tumor genotypes indicated. Each row is an individual tumor. Black is low, yellow is high. n=3 for EGFR ON CRP lung tumors, n=9 for EGFR OFF CRP lung tumors, n=2 for EGFR ON ERP lung tumors, n=3 for EGFR OFF ERP lung tumors. **H.** Heatmaps of gene set enrichment scores of the indicated gene lists in *EGFR*-Mutant lung tumor genotypes indicated. Blue is low, yellow is high. IMA=invasive mucinous adenocarcinoma. TAM=tumor associated macrophages. **I**. Gene set enrichment score of MAPK target genes (Wagle, et al., 2018^67^) in the *EGFR*-Mutant lung tumor genotypes indicated. Student’s t test was used to calculate two-sided p-values. **J**. PCA plot of RNA-seq data from the *EGFR*-Mutant lung tumor genotypes indicated. CRP: sgControl; sg*Rb1*; *Trp53*-/-, ERP: sg*Eed*; sg*Rb1*; *Trp53*-/-. *=p<0.05, **=p<0.01, ***=p<0.001, ****=p<0.0001. EGFR ON=Dox ON, EGFR OFF=Dox withdrawal. See also Figure S9.

These adenoviruses were intratracheally (IT) injected into the mice above and 1 week following IT injection, all mice were maintained on Dox-containing chow to induce expression of the EGFR L858R oncogene. Pilot experiments showed that after 20 weeks on Dox-containing chow, mice developed symptoms of dyspnea related to lung tumors with lung pathology showing *EGFR*-Mutant LUAD (**Fig. S9B-C**), which is consistent with the latency observed in this EGFR L858R transgenic mouse model^48,49^. Therefore after ∼20 weeks, both *EGFR*-Mutant *CRP* and *EGFR*-Mutant *ERP* mice were randomized to two groups with one group being maintained on Dox (Dox ON) and the other group switched back to regular chow (Dox OFF) to turn off expression of the EGFR L858R oncogene. Animals were then followed for symptoms until they reached their endpoint at which point necropsies were performed (**Fig. 6B**). Consistent with EGFR L858R acting as a potent oncogene, both *EGFR*-Mutant *CRP* and *EGFR*-Mutant *ERP* Dox ON mice more rapidly succumbed to their disease with a median overall survival of 30.3 weeks for *EGFR*-Mutant *CRP* and 31.0 weeks for *EGFR*-Mutant *ERP* mice (**Fig. 6C**). Survival was significantly prolonged in both *EGFR*-Mutant *CRP* and *EGFR*-Mutant *ERP* Dox OFF mice with a median overall survival of 57.9 and 57.1 weeks, respectively (**Fig. 6C**).

Interestingly, the phenotypes of *EGFR*-Mutant *CRP* Dox OFF and *EGFR*-Mutant *ERP* Dox OFF mice were quite distinct. Tumors eventually recurred in *EGFR*-Mutant *CRP* Dox OFF mice with a significant number of mice (7 of 12) developing spontaneous distant metastases to the liver, kidney, and brain, while none of the *EGFR*-Mutant *ERP* Dox OFF mice developed spontaneously metastases (**Fig. 6D, S9D**). This is consistent with our observation that EED inactivation in the SCLC *RPR2* GEMM blocked distant metastasis (**Fig. 1F**). IHC validated expression of the EGFR L858R transgene according to Dox treatment, GFP expression in tumor cells, and loss of EED protein only in *EGFR*-Mutant *ERP* mice but not in *EGFR*-Mutant *CRP* mice (**Fig. S9E**). DNA sequencing and immunoblot analysis confirmed expect patterns of inactivation in *EGFR*-Mutant *CRP* and *EGFR*-Mutant *ERP* tumors (**Fig. S9C,F,G**). Consistent with concurrent *RB1* and *TP53* mutations significantly increasing risk for SCLC transformation in human *EGFR*-Mutant LUAD^14^, some tumors in the *EGFR*-Mutant *CRP* Dox OFF mice (3 of 12) recurred as SCLC consistent with SCLC transformation where no tumors in the *EGFR*-Mutant *ERP* Dox OFF mice recurred as SCLC (**Fig. 6E**) suggesting that EED inactivation blocks LUAD-to-SCLC histological transformation. IHC demonstrated that transformed SCLC tumors in the *EGFR*-Mutant *CRP* Dox OFF mice either expressed only ASCL1 or heterogeneously expressed ASCL1 and NEUROD1, while the *EGFR*-Mutant *ERP* counterparts were completely negative (**Fig. 6F**), which is consistent with *EGFR*-Mutant human transformed SCLCs showing heterogenous expression of transcription factors with enrichment of variant subtypes including NEUROD1^50^. This also provided evidence to support that these tumors transformed into SCLC as a mechanism of escape from loss of EGFR L858R rather than formed as *de novo* SCLC as NEUROD1 expression is not observed in the *RPR2* SCLC GEMM^20,21,28,51^. Consistent with the *EED*-Mutant *RPR2* GEMM model, both *EGFR*-Mutant *ERP* Dox ON and Dox OFF tumors also showed evidence of LUAD with mucinous features (**Fig. 6F**). In line with this, there were only a small shared subset of 36 genes marked by bivalency in both human SCLC and human LUAD and this shared subset were highly enriched in drivers of gastric differentiation including *FGFR2* and *FGF18* (**Fig. S9H**).

Consistent with our IHC data, bulk RNA-seq data showed increased expression of NE markers including *Syp, Insm1*, and *Ncam1* as well as *Ascl1* with heterogenous *Neurod1* and enrichment for SCLC-related signatures (e.g. MYC, G2M pathways, pancreas beta cell signatures) in transformed SCLCs relative to *EGFR*-Mutant *CRP* Dox ON tumors, *EGFR*-Mutant *CRP* Dox OFF tumors that did not transform to SCLC, or all *EGFR*-Mutant *ERP* tumors (**Fig. 6G-H, S9I-K**). Moreover, transformed SCLCs had lower MAPK target gene expression (**Fig. 6I**). *EGFR*-Mutant *CRP* Dox OFF LUADs were higher grade tumors relative to *EGFR*-Mutant *CRP* Dox ON LUADs and enriched expression of genes involved in EMT consistent with its poorly differentiated histology and higher propensity to metastasize (**Fig. S9L, see also** **Fig. 6D**). In line with findings in *EED*-Mutant *RPR2* tumors, genes expressed in mucinous LUAD, tumor associated macrophages, and neutrophils were significantly enriched in both *EGFR*-Mutant *ERP* Dox ON and OFF tumors relative to their *EGFR*-Mutant *CRP* counterparts (**Fig. 6H, S9M-N**). Together, these data demonstrate that *EGFR*-mutant LUADs with *RB1* and *TP53* loss can evade EGFR dependency through either SCLC transformation or poorly differentiated metastatic LUAD. EED inactivation blocks both escape routes—preventing SCLC transformation and aggressive LUAD recurrence—and instead promotes a mucinous LUAD phenotype as a distinct mechanism of resistance to EGFR oncogene withdrawal (**Fig. 6J**).

### PRC2 Represses LUAD Oncogenic Programs in Human SCLC Through Bivalent Silencing of RAS, PI3K, and MAPK Pathway Genes

To evaluate the clinical relevance of our findings and determine whether PRC2-mediated repression of LUAD oncogenic programs is conserved in human tumors, we analyzed H3K27me3 and H3K4me3 ChIP-seq data from 5 patient-derived xenograft (PDX) models of *de novo* neuroendocrine (NE) SCLC and 4 PDXs of *de novo EGFR*-Mutant LUAD^52^ (**Fig. 7A-B, 7F)**. *EGFR*-Mutant LUAD was selected given its established susceptibility to SCLC transformation^3^, making it a pertinent comparator for lineage plasticity. This allowed us to test whether PRC2 activity differentially regulates lineage-specific epigenetic programs in human lung cancer.

**Figure 7.**
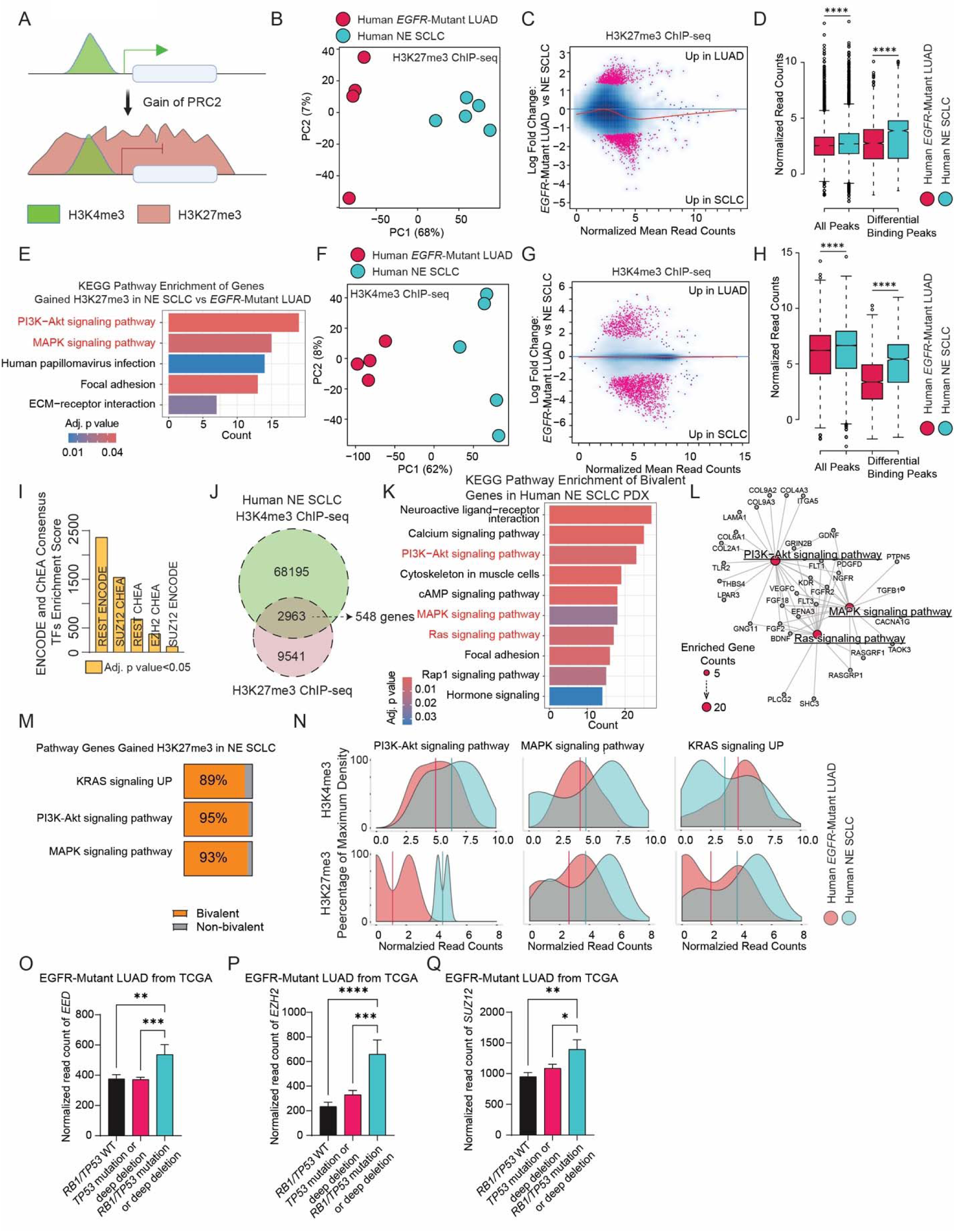
PRC2 Represses LUAD Oncogenic Programs in Human SCLC Through Bivalent Silencing of RAS, PI3K, and MAPK Pathway Genes. **A**. Schematic showing gain of PRC2-mediated repression of genes with bivalent H3K4me3 and H3K27me3 histone marks. Figure was made with BioRender. **B, F**. PCA plot of H3K27me3 ChIP-seq (**B**) and H3K4me3 ChIP-seq (**F**) from 5 independent human *de novo* SCLC neuroendocrine patient-derived xenograft (PDX) tumors and 4 independent human *EGFR*-Mutant LUAD PDXs tumors. **C,G**. Log ratio/average (MA) plot comparing *EGFR*-Mutant LUAD PDX tumors vs. *de novo* SCLC neuroendocrine (NE) PDX tumors above to visualize the relationship between the log ratio (M) and the average (A) of H3K27me3 ChIP-seq data (**C**) and H3K4me3 ChIP-seq data (**G**). **D,H**. Box plots of normalized read counts of H3K27me3 ChIP-seq (**D**) and H3K4me3 ChIP-seq (**H**) in the indicated groups. **E**. Bar plot of enriched KEGG pathways in genes mapped to the differential H3K27me3 peaks gained in NE SCLC PDX tumors vs. *EGFR*-Mutant LUAD PDX tumors. BH adjusted p-values are visualized by the bar color, red=low, blue=high. Count: Enriched gene counts. **I**. Bar plot of ENCODE and ChEA Consensus TFs enrichment score from top enriched transcription factors whose targets were enriched in genes mapped to the differential H3K4me3 peaks that gained signal in NE SCLC PDX tumors. Yellow: significantly enriched with BH adjusted p-value<0.05. **J**. Schematic to identify bivalent genes marked by both H3K4me3 and H3K27me3 in human NE SCLC PDX tumors. **K**. Bar plot of enriched KEGG pathways in genes marked with bivalent H3K4me3 and H3K27me3 in human NE SCLC PDX tumors. BH adjusted p-values were visualized by the bar color, red=low, blue=high. Count: Enriched gene counts. **L**. Gene network analysis of enriched RAS, PI3K-AKT, and MAPK pathways from **P**. Enriched gene counts are visualized by dot size. **M**. Bar plots indicating the percentage of genes marked by H3K4me3 of the indicated pathways that gained H3K27me3 in NE SCLC PDX tumors and and hence are bivalent. **N**. Density plots of normalized read count of H3K4me3 (top) and H3K27me3 (bottom) ChIP-seq peaks mapped to genes that belong to KRAS signaling UP, PI3K-Akt and MAPK KEGG pathways in human NE SCLC and *EGFR*-mutant LUAD PDX tumors. **O-Q**. Bar plots of RNA-seq batch normalized expression of *EED* (**O**), *EZH2* (**P**), *SUZ12* (**Q**) from human *EGFR*-Mutant LUAD tumors with the *RB1* and *TP53* genetic alterations indicated from the Cancer Genome Atlas (TCGA)^68^. *RB1/TP53* WT: n=22 tumors, *TP53* mutation or deep deletion: n=34 tumors, *RB1/TP53* mutation or deep deletion: n=10 tumors. One way ANOVA with Tukey test was used to calculate two-sided p values adjusted for multiple comparisons. *=p<0.05, **=p<0.01, ***=p<0.001, ****=p<0.0001. See also Figure S10.

ChEA and ENCODE transcription factor analyses of H3K27me3 peaks identified SUZ12 and EZH2 as dominant regulators of both histologies (**Fig. S10A-B**), validating the fidelity of the ChIP-seq dataset. Consistent with our data from mouse tumors and previous studies^46,47^, GO analysis showed MHC I antigen presentation genes were enriched for H3K27me3 in SCLC, while axon guidance pathways were enriched for H3K27me3 in LUAD (**Fig. S10C-D**) together showing that H3K27me3 gene repression patterns helps define key differences between SCLC and LUAD. *De novo* NE SCLC had overall more and stronger H3K27me3 peaks relative to LUAD and the top gained H3K27me3 peaks in SCLC vs. LUAD included PI3K and MAPK pathway genes (**Fig. 7C-E**)—mirroring findings from our GEMM studies. SCLC also showed elevated H3K4me3 at neurodevelopmental genes (**Fig. 7G–H, S10E-F**), whereas LUAD-specific H3K4me3 peaks enriched for EMT, inflammatory response, and KRAS signaling (**Fig. S10G-I**). Unexpectedly, the strongest H3K4me3 peaks gained in SCLC enriched for PRC2 targets, suggesting that increased PRC2 activity may coincide with H3K4me3 deposition at poised, bivalent loci (**Fig. 7I**). To test this, we identified bivalent promoters (co-marked by H3K27me3 and H3K4me3) in human SCLC and LUAD. We found 548 bivalent genes in SCLC and only 202 in LUAD (**Fig. 7J, S10J**). Bivalent genes in SCLC were significantly enriched for RAS, PI3K-AKT, and MAPK signaling, including *RASGRP1* (**Fig. 7K–L**), and were largely distinct from those in LUAD (**Fig. S10K-L**). Notably, ∼90% of all RAS/ PI3K-AKT/MAPK pathway genes that gained H3K27me3 in SCLC were bivalent (**Fig. 7M**), and genome-wide H3K27me3 signal was substantially higher across these pathways in SCLC than LUAD (**Fig. 7N**). Finally, to investigate why PRC2 activity is elevated in SCLC, we analyzed expression of PRC2 components in human LUAD samples from TCGA. Given that *RB1* and *TP53* loss is nearly universal in SCLC^11^ and common in LUADs that undergo transformation to SCLC^14^, and that RB1 loss promotes E2F transcription factor activity, which directly drives expression of PRC2 components such as EZH2 and EED^53^, we hypothesized that RB1 inactivation is associated with elevated PRC2 expression. Indeed, *RB1/TP53*-mutant LUADs showed significantly higher expression of all three core PRC2 components (EZH2, EED, and SUZ12) compared to *RB1/TP53*-WT LUAD tumors (**Figs. 7O-Q, S10M-O**). Together, these data provide human tumor evidence that PRC2 blocks LUAD oncogenic potential in SCLC by repressing key signaling effectors through bivalent chromatin, and suggest a mechanistic axis in which RB1 and *TP53* loss drives PRC2 overexpression to silence bivalent RAS, PI3K, and MAPK genes—thereby enforcing the neuroendocrine phenotype.

## Discussion

Here, we demonstrate that EED, a core component of the PRC2 complex, is essential for maintaining the ASCL1-positive NE identity and histological state of SCLC during tumorigenesis. When tumorigenesis is initiated in the *RPR2* genomic context that normally drives formation of SCLC tumors, tumors escape from EED inactivation by losing ASCL1 and adopting LUAD histology. Alternatively, in an *EGFR*-Mutant LUAD GEMM with concurrent *Rb1* and *Trp53* loss—genetic lesions strongly associated with LUAD-to-SCLC transformation^14^—EED was required for SCLC histology to emerge following EGFR oncogene withdrawal. Together, these findings identify EED as a critical determinant of the SCLC NE histological fate in lung cancer.

Mechanistically, EED directly binds and promotes H3K27me3-mediated gene silencing of RAS, PI3K, and MAPK signaling genes to repress LUAD oncogenic signaling and maintain SCLC NE identity. These loci are bivalent—marked by both H3K27me3 and H3K4me3—in SCLC, rendering them epigenetically poised for rapid activation. Loss of EED triggers SCLC-to-LUAD histological transformation by derepressing these bivalent targets, unleashing LUAD oncogenic signaling through a NEUROD1-positive intermediate state that requires cues from the tumor microenvironment (**Fig. 8**). Supporting this model, ChIP-seq analysis of human NE SCLC PDXs revealed bivalent chromatin marks at the same RAS, PI3K, and MAPK pathway genes, indicating dynamic PRC2-mediated repression in human disease. This supports a model in which PRC2 activity silences LUAD oncogenic signaling to enforce and maintain SCLC identity.

**Figure 8.**
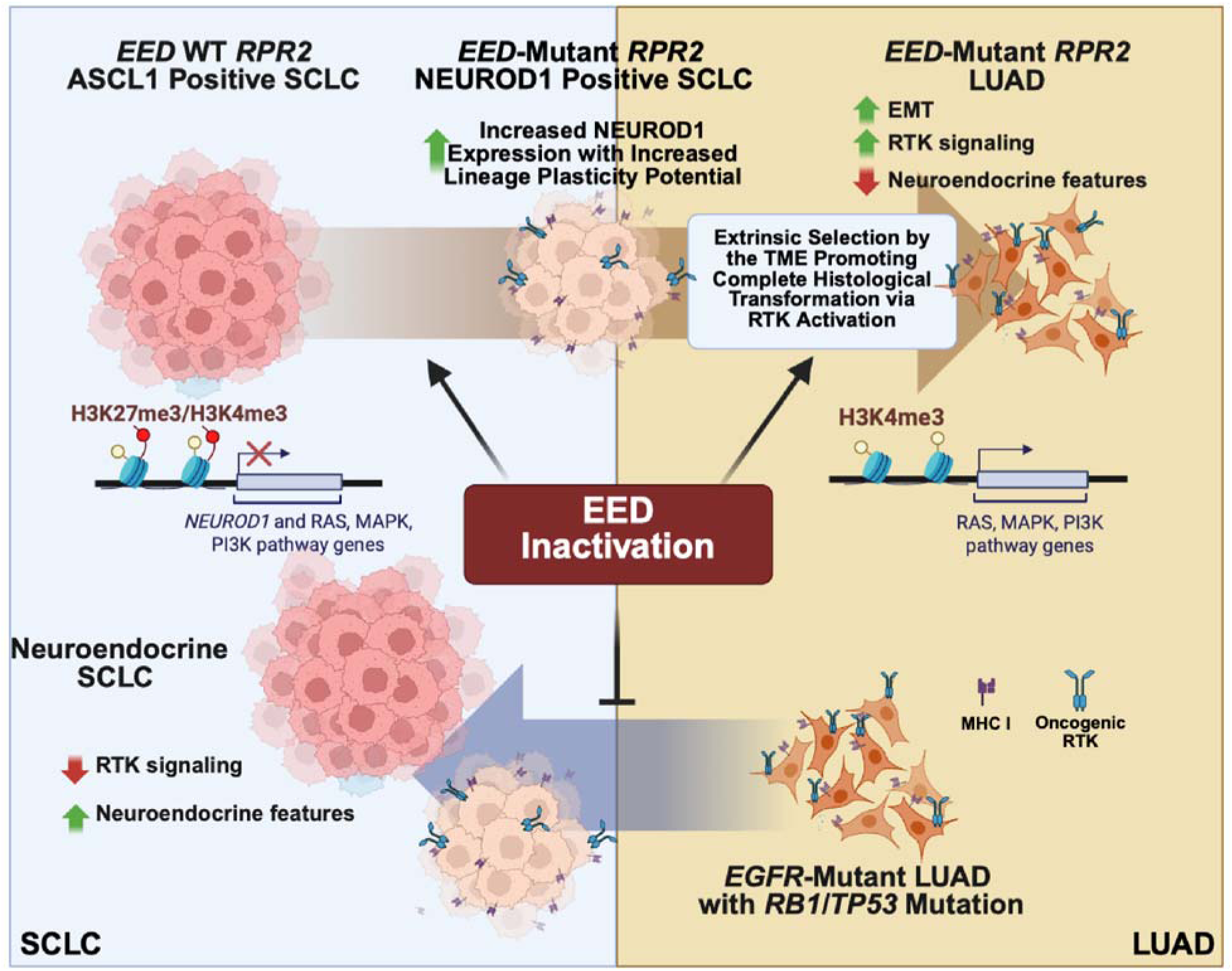
Schematic Showing How EED Constrains the SCLC Neuroendocrine Phenotype and Drives Lung Cancer Histological Transformation. When SCLC tumorigenesis is initiated in the *RPR2* GEMM model, inactivation of *EED* delays tumor formation and tumors eventually form by a histological transformation to lung adenocarcinoma (LUAD). SCLC-to-LUAD histological transformation occurs through a transient NEUROD1-positive EMT-like intermediate cell state that requires cues from the tumor microenvironment. When LUAD tumors are initiated by the EGFR L858R oncogene and recur after EGFR oncogene withdrawal, only tumors with EED-intact undergo histological transformation to SCLC as a mechanism of resistance to EGFR withdrawal. Together both models show that intact EED is strictly required to promote the SCLC neuroendocrine phenotype during lung cancer pathogenesis. Mechanistically, EED maintains the SCLC neuroendocrine identity by directly repressing bivalent chromatin-marked RAS, PI3K, and MAPK pathway genes to oppose LUAD oncogenic signaling. RTK: receptor tyrosine kinase. EMT: epithelial to mesenchymal transition. Figure created with BioRender.

Human lung cancers can exist as combined LUAD and SCLC histology tumors and can also histologically transform from LUAD to SCLC as a mechanism of resistance to targeted therapy^2,3^. Our data suggests that mechanisms to promote PRC2 complex activity would select for SCLC histology in each of these settings. Although SCLC-to-LUAD transformation has not been observed in human tumors, this may reflect the limited efficacy of current SCLC therapies, which exert less selective pressure to escape the SCLC neuroendocrine state. Recently the highly effective neuroendocrine targeted therapy Tarlatamab, a DLL3 bispecific T cell engager, was approved^54,55^. It will be important to determine whether resistance to Tarlatamab could involve SCLC-to-LUAD histological transformation as a strategy to escape the neuroendocrine state and promote tumor survival.

EED inactivation in both SCLC *RPR2* GEMMs and *EGFR*-Mutant *RP* LUAD GEMMs led to LUAD with aberrant expression of gastrointestinal differentiation genes. Similarly, prior work showed that EED loss promoted mucinous LUAD histology in *KRAS*-mutant LUAD GEMMs^56,57^, suggesting that PRC2 normally represses GI differentiation programs in lung tumors. We speculate that this function of PRC2 mirrors the foregut endoderm anterior to posterior patterning where it likely represses genes involved in GI differentiation such as *HNF4A*, *WNT* and *FGF* signaling to give rise to the respiratory system and inhibit gut development^58^. We found that EED inactivation also de-repressed *NEUROD1* *in vitro* and occasionally drove ASCL1-to-NEUROD1 subtype switching in SCLC tumors *in vivo*. *NEUROD1* is a direct EED target marked by bivalent H3K4me3 and H3K27me3, enabling rapid activation upon PRC2 loss. While ASCL1 is expressed in pulmonary neuroendocrine cells^59^, NEUROD1 is absent in the lung but highly expressed in endocrine pancreatic β cells^60–62^, which are also derived from foregut endoderm^63^. There are several studies that now show ASCL1 to NEUROD1 subtype switching in SCLC and NEPC after selective pressure against ASCL1^20,21,64^. Our findings reveal another mechanism that drives ASCL1 to NEUROD1 subtype switching and suggests that this can occur when tumor cells adopt development programs expressed in pancreatic β cells. While our previous work showed that the epigenetic modifier KDM6A maintains an epigenetic state permissive for the ASCL1 subtype by preferentially through epigenetic activation of genes associated with the ASCL1 subtype^21^, this works shows that the PRC2 complex achieves an ASCL1-positive subtype by repressing genes associated with the NEUROD1 subtype.

Following EED inactivation, NEUROD1-positive SCLC tumors were rarely sustained, and nearly all mice ultimately developed LUADs, suggesting that NEUROD1-positive SCLCs are either transient and selected against, or that LUAD histology is actively selected for. We observed NEUROD1-positive cells within LUADs that emerged after EED loss, and our snRNA-seq data identified a NEUROD1-expressing, EMT-like subpopulation, supporting a model in which LUADs arise from SCLC through a NEUROD1-positive intermediate. Notably, EED inactivation alone was insufficient to drive this transition *in vitro*, implying that the tumor immune microenvironment (TIME) is required to complete SCLC-to-LUAD histological transformation, potentially by enhancing LUAD oncogenic signaling. Alternatively, prior work, including our own, has shown that PRC2 loss in SCLC increases MHC class I expression^46,47^, which correlates with enhanced immune visibility and response to checkpoint blockade^46,65^. It is therefore possible that EED-inactivation renders SCLC cells immunogenic and subject to immune-mediated clearance, thereby favoring outgrowth of LUAD. Consistent with this, EED inactivated tumors exhibited a marked increase in immunosuppressive M2-like macrophages, which may emerge in response to restored MHC class I expression and facilitate immune evasion. Future studies will aim to dissect how the TIME contributes to lineage plasticity in lung cancer and whether transformation between lung cancer histological subtypes reflects tumor-intrinsic reprogramming, immune selection, or both.

LUAD-to-SCLC transformation is a well-established mechanism of resistance to targeted therapies, most frequently observed following EGFR inhibition in *EGFR*-Mutant LUADs with co-occurring *RB1* and *TP53* mutations^2,3,5^. While these alterations increase the relative risk of transformation by 43-fold^14^, only ∼18% of such cases undergo SCLC transformation clinically^8^. In our study, EED inactivation fully blocked the ability of *EGFR*-Mutant LUADs with *Rb1* and *Trp53* loss to transform into SCLC following EGFR oncogene withdrawal. Although only ∼25% of tumors in our model exhibited SCLC histology—closely aligning with the rate in *EGFR*-Mutant *RB1*/*TP53* mutated human tumors—most remaining tumors recurred as poorly differentiated, metastatic LUADs, highlighting divergent lineage escape routes. This incomplete penetrance likely reflects the absence of additional co-occurring events, such as PI3K pathway activation^5,10,15^ or MYC overexpression^10^, which are frequently found in human LUADs that undergo lineage transformation. Indeed, a recent *EGFR*-Mutant GEMM incorporating non-degradable c-MYC showed a higher frequency of LUAD-to-SCLC transformation^10^.

Our study identifies PRC2 as a central epigenetic gatekeeper of lineage plasticity in lung cancer, maintaining the SCLC neuroendocrine phenotype by repressing LUAD oncogenic signaling through bivalent chromatin silencing of RAS, PI3K, and MAPK pathway genes. Integrating murine models with human PDX ChIP-seq data, we demonstrate that these signaling genes are marked by both H3K27me3 and H3K4me3 in SCLC, positioning them for rapid activation upon PRC2 loss. Importantly, we show that *RB1* and *TP53* co-mutation—common in both *de novo* SCLC and LUADs at risk of SCLC transformation—is associated with elevated expression of PRC2 components (EZH2, EED, SUZ12). These findings suggest a mechanistic link between genetic drivers and chromatin-based lineage restriction, and implicate PRC2 activity as a driver of LUAD-to-SCLC transformation under therapeutic pressure. As such, targeting PRC2 may represent a rational strategy to prevent histological transformation in high-risk *EGFR*-Mutant LUADs with *RB1* and *TP53* loss, and more broadly, to modulate lineage plasticity in lung cancer.

## Limitations of the study

Our data suggests that inhibition of the PRC2 complex with small molecule EZH1/2 or EED inhibitors as a therapeutic approach to block SCLC histological transformation in patients at high-risk for SCLC transformation; particularly patients with *EGFR*-Mutant LUADs with co-occurring *RB1* and *TP53* inactivation^8,14^. However, in our experiments, EED was genetically inactivated at tumor initiation in mice, not in established EGFR oncogene-driven tumors. Our analysis of ChIP-seq data from human LUAD and SCLC PDXs uncovering a gain of H3K27me3 in SCLC to silence genes involved in RAS, PI3K, and MAPK signaling, and drive SCLC histology supports this therapeutic rationale in human tumors, which is consistent with a previously published independent data in human tumors showing a gain of PRC2 complex transcriptional signatures during LUAD-to-SCLC transformation^5^. Future studies will focus on determining whether clinical grade inhibitors that disrupt the PRC2 complex can block histological transformation in established tumors driven by EGFR or other LUAD oncogenic drivers such as ALK rearrangements or KRAS mutations, where SCLC histological transformation has also been observed; albeit less commonly^66^.

## Acknowledgements

This work was supported by a William Raveis Charitable Fund Damon Runyon Clinical Investigator Award (CI-101-19, M.G.O.), a DF/HCC SPORE CEP Award (P50CA265826, M.G.O.), an NCI P01 award (P01CA295524, M.G.O.), an American Cancer Society Postdoctoral Fellowship (PF-24-1252572-01-CCB, Y.L.), the Kaplan Family Fund (M.G.O), and an NCI R01 award (R01CA263715, K.P.). S.L. is supported by the NCI Research Specialist Award (R50CA251956). We thank Dr. David Barbie for thoughtful discussions and members of the Oser, Barbie, and Janne labs for their critical feedback. We thank the DFCI Molecular Biology Core Facility including Zach Herbert and Maura Berkeley who used an Illumina NovaSeq X Plus that was purchased with funding from a National Institutes of Health SIG grant 1S10OD036228-01 for this work. We also thank the Dana-Farber/Harvard Cancer Center for the use of the Specialized Histopathology Core, which provided histology and immunohistochemistry service. Dana-Farber/Harvard Cancer Center is supported in part by an NCI Cancer Center Support Grant # NIH 5 P30 CA06516. The results in Figs. 7O-Q, S10M-O are based upon data generated by the TCGA Research Network: https://www.cancer.gov/tcga.

## Author Contributions

Y.L.: Conceptualization, methodology, validation, formal analysis, investigation, data curation, supervision, writing. Y.N.L, H.C., G.R.D.O., T.E.Z.: Methodology, validation, formal analysis, investigation, data curation. M.V., Y.D., A.D., X.Q., S.K., R.L., R.B., S.L., W.L., M.V.O.: Investigation. K.P., J.B., M.F., H.L., S.S.: Resources, visualization, supervision, project administration. Y.C., H.J.: Methodology. M.G.O: Conceptualization, methodology, investigation, resources, data curation, writing, visualization, supervision, project administration, funding acquisition.

## Methods

All experiments herein comply with all ethical regulations. Specifically, all mouse experiments complied with National Institutes of Health guidelines and were approved by Dana-Farber Cancer Institute Animal Care and Use Committee (DFCI, protocol 19-009). All adenoviral and lentiviral transduction experiments complied with the Biohazard Control Committee (DFCI, protocol 19-1133).

### Adenoviral sgRNA Expression Vector Cloning to Create *RPR2* Genetically Engineered Mouse Model

Effective sgRNAs targeting mouse *Rb1*, *Trp53*, and *Rbl2* were first validated using lentiviral vectors as described previously^26^. Effective sgRNAs targeting mouse *Eed* were validated in mouse embryonic fibroblasts expressing Cas9. The cloning method for generation of adenoviral sgRNA expression vectors encoding CMV-Cre recombinase and sgRNAs targeting *Rb1, Trp53, and Rbl2* and “T” sgRNA (in this case sgEed#1, sgEed#2, or a non-targeting sgRNA control, *C0111*) was also described previously^25,27^. Briefly, a pENTR223-CMV-Cre-U6-sgX-U6-sgRb1-U6-sgTrp53-U6-sgRbl2 where X is sgEed#1, sgEed#2, or sgC0111, was used in an LR recombination reaction to clone the 4 pENTR223-CMV-Cre-U6-sgX-U6-sgRb1-U6-sgTrp53-U6-sgRbl2 vectors described above into pAd-PL DEST (Invitrogen) according to the manufacturer’s instructions. The recombinants were transformed into HB101 cells and ampicillin-resistant colonies were screened by restriction digestion of miniprep DNA and subsequently validated by DNA sequencing. The following sgRNA oligos were used (including BsmBI sites): *Rb1* mouse #11 sense (5’- CACCGCAACTAGAAAATGATACG-3’), *Rb1* mouse #11 anti-sense (5’- AAACCGTATCATTTTCTAGTTGC-3’), *Trp53* mouse #8 sense (5’- CACCGGTGTAATAGCTCCTGCATGG-3’), *Trp53* mouse #8 anti-sense (5’- AAACCCATGCAGGAGCTATTACACC-3’), *Rbl2* mouse #6 sense (5’- CACCGAGGAGGATGGCGACGCCG-3’), *Rbl2* mouse #6 anti-sense (5’- AAACCGGCGTCGCCATCCTCCTC-3’), *Eed* mouse #1 sense (5’- CACCGACAAATACGCCAAATGCACC-3’), *Eed* mouse #1 anti-sense (5’- AAACGGTGCATTTGGCGTATTTGTC-3’), *Eed* mouse #2 sense (5’-CACCGTGCACCAGGAAGGAAAAGCT-3’), *Eed* mouse #2 anti-sense (5’-AAACAGCTTTTCCTTCCTGGTGCAC-3’), *C0111 (*Non-targeting sgRNA, sgControl) sense (5’- CACCGGGAGGCTAAGCGTCGCAA-3’), *C0111* (Non-targeting sgRNA, sgControl) anti-sense (5’- AAACTTGCGACGCTTAGCCTCCC-3’).

### Adenoviral sgRNA Expression Vector Cloning to Create *EGFR*-Mutant Genetically Engineered Mouse Model with Concurrent *Rb1* and *Trp53* Mutation

The cloning method for generation of adenoviral sgRNA expression vectors encoding CMV-Cre recombinase and sgRNAs targeting *Rb1* and “T” sgRNA (in this case sgEed#1, sgEed#2, or a non-targeting sgRNA control) is described below. The DNA fragment containing CMV-Cre-U6-sgX-U6-sgRb1 was generated by inverse polymerase chain reaction (PCR) using KOD Xtreme™ Hot Start DNA Polymerase (Sigma-Aldrich, Cat# 71975) with primers (sense: 5’-CACCCAGCTTTCTTGTACAAAGTTGGC-3’, and anti-sense: 5’-TTGTACAAGAAAGCTGGGTGTTGTATA-3’) from the pENTR223-CMV-Cre-U6-sgX-U6-sgRb1-U6-sgTrp53-U6-sgRbl2 template. Linear DNA fragment was annealed into a circular plasmid by Gibson Assembly (NEB, cat# E5510S) according to the manufacturer’s instructions. The plasmid was then transformed in HB101 cells, and isolated using the QIAprep Spin Miniprep Kit – Plasmid Purification Kit (Qiagen, 27106). Whole plasmid sequencing was performed to confirm the correct sequences (pENTR223-CMV-Cre-U6-sgX-U6-sgRb1 where sgX is sgEed#1, sgEed#2, or a non-targeting sgRNA control). Subsequently LR reactions were performed with pAd-PL DEST as described above and previously^25,27^ to make pAd-PL CMV-Cre-U6-sgX-U6-sgRb1 where sgX is sgEed#1, sgEed#2, or a non-targeting sgRNA control.

### Adenovirus Production and Purification

Adenoviral production and purification were performed as described previously^25^. 5 μg of the adenovirus vector (pAd/PL Invitrogen #V494-20) containing the desired sgRNA sequences and Cre recombinase expression cassette (see above) was digested with PacI (New England Biolabs) for 2 hours at 37°C according to the manufacturer’s instructions and column purified using Qiagen’s gel extraction kit. 1 μg of PacI-digested pAd/PL was transfected into 1.5 X 10^6^ 293AD cells plated on a 6 cm tissue-culture dish using Lipofectamine 2000. The following day, the media was exchanged, and subsequently exchanged every 48 hours thereafter. Once 293AD cells showed evidence of adenovirus production (determined by comet formation with lysis), the cells and supernatant were harvested, which were then subjected to 4 freeze-thaw cycles by alternating between an ethanol dry ice bath and 37°C. Cell debris was removed by centrifugation and the supernatant was collected, passed through a 0.45 μm filter, aliquoted, and frozen at -80°C until use.

To generate high titer adenovirus for *in vivo* experiments, adenovirus was generated as described above. 50 μl of the adenovirus stock was added to each 10 cm tissue-culture dish of 293FT cells plated at 3 X 10^6^ cells per dish (4 10 cm dishes in total for each purification). When 293FT cells showed evidence of adenovirus production, as determined by cell rounding and partial detachment (∼48-72 hours after addition of adenoviral stock), the cells were collected, and adenovirus was purified using Virabind Adenovirus Purification Kit (Cell Biolabs #VPK-5112). The purified adenovirus was titered using QuickTiter Adenovirus Quantitation Kit (Cell Biolabs #VPK-106) according to the manufacturer’s instructions.

### Intratracheal Injections

Intratracheal injections were performed as described previously^69^. Briefly, mice were anesthetized with ketamine and xylazine and pedal reflexes were monitored to ensure adequate anesthesia. Mice were maintained on a heated stage at 37° C while anesthetized. Mice were hung on stage with their top incisors and intubated with a 22-gauge 1 inch catheter (ThermoFisher Scientific #1484120). Once intubated, adenovirus (4 X 10^8^ VP/mouse) in a total volume of 75 μl (diluted in PBS) was added to the catheter and subsequently inhaled by the mice.

### Establishing Genetically-Engineered Mouse Models using CRISPR/Cas9

For the *RPR2* GEMM, pure congenic *Lox-stop-lox (LSL) Cas9* BL6J mice were purchased from Jackson Labs (Jackson No. 026175) and maintained as homozygous BL6J mice. Genotyping of Cas9 and GFP at the ROSA26 were confirmed for all mice on the study (Transnetyx). Above-mentioned *RPR2* adenovirus was injected into these mice to generate *EED*-isogenic *RPR2* tumors.

For the *EGFR*-Mutant GEMM with concurrent *Rb1* and *Trp53* Mutation, *TetO-EGFR^L858R^; Trp53^flox/flox^; Rosa26^CAGs-LSL-rtTA3-IRES-mKate^* and *TetO-EGFR^L858R^; Trp53^flox/flox^; Rosa26^CAGs-LSL-Cas9-GFP^* were a kind gift from the laboratory of Dr. Katerina Politi^49^. These mice are mixed BL6/129/FVB background and maintained as homozygous strains. These two strains were bred together to generate *TetO-EGFR^L858R^; Trp53^flox/flox^; Rosa26^CAGs-LSL-rtTA3-IRES-mKate^; Rosa26^CAGs-LSL-Cas9-GFP^* mice used in this study (referred to hereafter as *EGFR*-Mutant GEMM experimental mice). Genotyping of *EGFR^L858R^, Trp53^flox/flox^*, and *Cas9, GFP*, and *rtTA3* at the *ROSA26* locus were confirmed for all mice (Transnetyx). For experiments in Fig. 6 and S9, pAd-PL CMV-Cre-U6-sgEed#2-U6-sgRb1 or pAd-PL CMV-Cre-U6-sgControl-U6-sgRb1 adenoviruses were intratracheally injected as above into the *EGFR*-Mutant GEMM experimental mice and then all mice were fed doxycycline-containing chow (625 ppm, cat# 5AW9, ScottPharma Solutions Inc.). For the control *de novo* LUAD tumors in Fig. 4A, *RPR2* adenovirus were used to match the adenoviruses used to make the SCLC *RPR2* model and intratracheally injected into the *EGFR*-Mutant GEMM experimental mice followed by administration of doxycycline-containing chow (625 ppm) after 1 week. For all studies with the *RPR2* GEMM and the *EGFR*-Mutant GEMM, both male and female mice were used at roughly equal numbers.

Housing conditions for mice at the DFCI Vivarium include a 12 hour/12 hour day-night cycle where temperature is maintained at 72 F. Roughly equal numbers of male and female 3-4 month-old mice were intratracheally injected with 4 X 10^8^ VP/mouse adenovirus. All mice were monitored at least twice a week and euthanized when they became symptomatic (primarily respiratory distress), moribund, or lost 15% of their total body weight. The maximal tumor size allowed by the Dana-Farber Cancer Institute Animal Care and Use Committee is 2 cm and the maximal tumor size was not exceeded in any of our studies. Upon euthanization, half of the lung tumor specimen was immediately flash frozen on dry ice for subsequent DNA, RNA, and protein analysis, while the other half and the rest of lung was fixed in 10% formalin for 24 hours and then stored in 70% ethanol prior to paraffin embedding. For some tumors, cell lines were generated (see method below). Livers, kidneys, and brains were harvested and fixed and embedded as above. Slides were made for Hematoxylin and Eosin (H&E) and immunohistochemistry (IHC). H&E slides were analyzed by a specialized rodent pathologist Dr. Roderick Bronson, or pulmonary pathologist Dr. Marina Vivero for diagnosis.

### Generation of Cell Lines from Mouse Tumors and Cell Culture

sgControl (EED-WT) *RPR2* (631, 1014) cell lines were generated from CRISPR-based SCLC GEMMs (see above) as described previously^25^. Once tumors developed, mice were euthanized with CO2 and their tumors were quickly extracted, washed in ice cold PBS, and minced several times using an ethanol sterilized razor blade. 3mL of collagenase/hyaluronidase (Stem cell biology #07912) diluted 1:10 in complete RPMI media containing [10% FBS, P/S, and HITES (10 nM hydrocortisone (Sigma Aldrich # H0135), Insulin-Transferrin-Selenium (Gemini #400-145), and 10 nM beta-estradiol (Sigma Aldrich# E2257), 100 U/mL of penicillin (P), and 100 µg/mL of streptomycin (S)], and 1mL dispase (Corning # 354235) was added to the tumor, and incubated at 37°C for 20-40 minutes with periodic pipetting ∼10 every minutes (until most of the tumor cells were in suspension). The cells were then collected, centrifuged at 200 x g for 5 minutes, resuspended in complete RPMI media (see above), filtered through a 70 μm cell strainer (BD #352350), centrifuged again at 1000 rpm for 5 minutes, resuspended in fresh RPMI HITES media and placed in ultra-low adherence tissue culture dishes (Corning #3471). Media was subsequently replaced every 3 days. Histopathology on the tumors confirmed SCLC for all cell lines generated. All SCLC cell line were grown in Ultra-Low Attachment flasks (Corning™ 3814CONV) or plates (Corning™ 3471) at 37°C in the presence of 5% CO2. Once established, all cell lines were validated using immunoblot analysis for Cas9 and the SCLC neuroendocrine markers ASCL1.

sg*Eed RPR2* (1339, 1343, 1344, 1345, 1350) cell lines were generated from *EED*-Mutant *RPR2* LUAD tumors using Mouse Tumor Dissociation Kit (Miltenyi, cat# 130-096-730) following manufacturer’s instructions. All LUAD *RPR2* cell line were grown in RPMI media containing [10% FBS, 100 U/mL of penicillin (P), and 100 µg/mL of streptomycin (S) (P/S)] on tissue culture dishes (Falcon™, cat# 353003) at 37°C in the presence of 5% CO2. Once established, all cell lines were validated by immunoblot analysis for expression of Cas9, loss of TP53 and RB1 expression, and loss of ASCL1 expression, and by FACS for GFP expression. Early passage cell lines were tested for Mycoplasma (Lonza #LT07-218) and were negative, and then were frozen using Bambanker’s freezing media (Bulldog Bio). Mouse embryonic fibroblasts expressing Cas9 used to validate the adenoviruses were described previously^26^ and cultured in DMEM media with 10% FBS and P/S.

### Human Cell Lines

NCI-H1876, NCI-H2081 and 293FT cells were originally obtained from American Type Culture Collection (ATCC). 293AD cells (AD-100) were obtained from Cell Biolabs. NCI-H1876 and NCI-H2081 cells were maintained in DMEM/F12 media 5% FBS, P/S, and HITES. 293T and 293AD were maintained in DMEM media with 10% FBS and P/S. Early passage cell lines were tested negative for Mycoplasma (Lonza #LT07-218), and then were frozen using Bambanker’s freezing media (Bulldog Bio). All experiments were performed with cell lines that were maintained in culture for <3 months at which time an early passage cell lines were thawed. No commonly misidentified cell lines were used in this study.

### Pharmacological Inhibitors

The following chemicals (stored at -20°C or -80°C) were added to cell culture where indicated: MAK683 (Selleckchem #S8983, stock 10 mM in DMSO), Tazemetostat (Selleckchem #S7128, stock 10 mM in DMSO), Tulmimetostat (Selleckchem #E1497, stock 10 mM in DMSO), Valemetostat (Selleckchem #S8926, stock 10 mM in DMSO), Trametinib (Selleckchem #S2673, stock in 10mM in DMSO) or RMC-6236 (Selleckchem #E1597, stock 10mM in DMSO).

### sgRNA Cloning to Make Lentiviruses

sgRNA sequences were designed using the Broad Institute sgRNA designer tool (http://portals.broadinstitute.org/gpp/public/analysis-tools/sgrna-design) and synthesized by IDT technologies. The sense and antisense oligonucleotides were mixed at equimolar ratios (0.25 nanomoles of each sense and antisense oligonucleotide) and annealed by heating to 100°C in annealing buffer (1X annealing buffer 100 mM NaCl, 10 mM Tris-HCl, pH 7.4) followed by slow cooling to 30°C for 3 hours. The annealed oligonucleotides were then diluted at 1:400 in 0.5X annealing buffer.

For CRISPR/Cas9 knockout experiments in cells, the annealed oligos were ligated into LentiGuide Puro (Addgene #52963) for experiments in mouse SCLC cell lines. Ligations were done with T4 DNA ligase for 2 hours at 25°C. The ligation mixture was transformed into HB101 competent cells. Ampicillin-resistant colonies were screened by restriction digestion of miniprep DNAs and subsequently validated by DNA sequencing.

The following sgRNA oligos were used for LentiGuide Puro vector for CRISPR knockout experiments: *Eed* mouse #1 sense (5’- CACCGACAAATACGCCAAATGCACC-3’), *Eed* mouse #1 anti-sense (5’- AAACGGTGCATTTGGCGTATTTGTC-3’), *Eed* mouse #2 sense (5’-CACCGTGCACCAGGAAGGAAAAGCT-3’), *Eed* mouse #2 anti-sense (5’-AAACAGCTTTTCCTTCCTGGTGCAC-3’), *C0111* (Non-targeting sgRNA, sgControl) sense (5’- CACCGGGAGGCTAAGCGTCGCAA-3’), *C0111* (Non-targeting sgRNA, sgControl) anti-sense (5’- AAACTTGCGACGCTTAGCCTCCC-3’).

### Lentivirus Production

Lentiviruses were made by Lipofectamine 2000-based co-transfection of 293FT cells with the respective lentiviral expression vectors and the packaging plasmids psPAX2 (Addgene #12260) and pMD2.G (Addgene #12259) at a ratio of 4:3:1. Virus-containing supernatant was collected at 48 and 72h after transfection, pooled together (15 mL total per 10-cm tissue culture dish), passed through a 0.45-µm filter, aliquoted, and frozen at -80°C until use.

### Lentiviral Infection

The cells were counted using a Vi-Cell XR Cell Counter (Beckman Coulter) and 2 X10^6^ cells were resuspended in 1mL lentivirus with 8 μg/mL polybrene in individual wells of a 12 well plate. The plates were then centrifuged at 448 x g for 2h at 30° C. 16 hours later the virus was removed, and cells were grown for 72 hours before being placed under drug selection. Cells were selected in puromycin (0.5 μg/mL).

### Generation of SCLC Xenograft Model in BL6J and NCr Nude Mice

The syngeneic *RPR2* 1014 BL6J model was described previously^27^. 1014 cells were first transduced with plentiguide-puro lentiviruses containing an sgRNA targeting EED (*Eed* mouse #2 sense (5’-CACCGTGCACCAGGAAGGAAAAGCT-3’), *Eed* mouse #2 anti-sense (5’-AAACAGCTTTTCCTTCCTGGTGCAC-3’)) (EED-inactivated) or a non-targeting sgRNA as a control (EED-WT). Immunoblot analysis validated loss of EED protein with increased NEUROD1 expression (see Fig. 3K). Then 8x10^6^ *RPR2* 1014 with *EED*-WT or *EED*-Inactivated cells were washed 3 times in DPBS and then resuspended in DPBS with 33% Matrigel (Corning cat# 354234) and implanted subcutaneously into 7-9 week-old female albino B6 mice (C57BL/6J-Tyrc-2J, stock no. 000058; RRID:IMSR_JAX:000058) from The Jackson Laboratory (Bar Harbor, ME) or immunodeficient female NCr nude mice (Taconic #NCRNU). Once established, mice were euthanized with CO2, and half of each tumor was immediately flash frozen on dry ice for subsequent immunoblot analysis.

### Immunoblotting

Cell pellets were lysed in a modified EBC lysis buffer (50mM Tris-Cl pH 8.0, 250 mM NaCl, 0.5% NP-40, 5 mM EDTA) supplemented with a protease inhibitor cocktail (Complete, Roche Applied Science, #11836153001) and phosphatase inhibitors (PhosSTOP Sigma #04906837001). Soluble cell extracts were quantified using the Bradford Protein Assay. 20 µg of protein per sample was boiled after adding 3X sample buffer (6.7% SDS, 33% Glycerol, 300 mM DTT, and Bromophenol Blue) to a final concentration of 1X, resolved by SDS-PAGE using either 10% or 8% SDS-PAGE, semi-dry transferred onto nitrocellulose membranes, blocked in 5% milk in Tris-Buffered Saline with 0.1%Tween 20 (TBS-T) for 1h, and probed with the indicated primary antibodies overnight at 4°C. Membranes were then washed three times in TBS-T, probed with the indicated horseradish peroxidase conjugated (HRP) secondary antibodies for 1h at room temperature, and washed three times in TBS-T. Bound antibodies were detected with enhanced chemiluminescence (ECL) western blotting detection reagents (Immobilon, Thermo Fisher Scientific, #WBKLS0500) or Supersignal West Pico (Thermo Fisher Scientific, #PI34078). The primary antibodies and dilutions used were: Rabbit anti-EED (Cell Signaling #85322S, 1:1000), rabbit anti-ASCL1 (Abcam #Ab211327, 1:1000), rabbit anti-NEUROD1 [EPR4008] (Abcam #Ab109224, 1:1000), rabbit anti-RAS (E8N8L) (Cell Signaling #67648, 1:1000), rabbit anti-Rb1 (Abcam #181616, 1:2000), rabbit rodent specific anti-p53 (D2H9O) (Cell Signaling Technology #32532, 1:1000), rabbit anti-p130 (Abcam #Ab76234, 1:1000), rabbit anti-Cre Recombinase (D7L7L) (Cell Signaling Technology #15036, 1:1000), Mouse Anti-Cas9 (Cell Signaling Technology #14697, 1:1000), mouse anti-β-actin (clone AC-15) (Sigma, #A3854, 1:25,000), mouse α-Vinculin (hVIN-1) (Sigma, # V9131, 1:10000). Histone extractions were performed as described previously^26^ with the following primary antibodies: Rabbit anti-Histone H3 (D1H2) (Cell Signaling Technology #4499S, 1:1000) and rabbit anti-Tri-Methyl-Histone H3 (Lys27) (Cell Signaling Technology #9733S, 1:1000). The secondary antibodies and dilutions used were: Goat Anti-Mouse (Jackson ImmunoResearch #115-035-003) and Goat anti-Rabbit (Jackson ImmunoResearch #111-035-003) and used at 1:5000.

### Bulk RNA-Sequencing and Analysis

Tumors were harvested at necropsy and were flash-frozen. RNA was extracted using Quick-RNA™ Miniprep kit (Cat # R1055, Zymo Research, CA, USA) including a DNase digestion step according to the manufacturer’s instructions and RNA sequencing was performed as described below. Total mRNA samples in each experiment were submitted to Novogene Inc. The libraries for RNA-seq are prepared using NEBNext Ultra II non-stranded kit. Paired end 150bp sequencing was performed on Novaseq6000 sequencer using S4 flow cell. Sequencing reads were mapped to the mm10 genome by STAR. Statistics for differentially expressed genes were calculated by DESeq2 (1.36.0). Bulk RNA-sequencing heatmaps were generated by calculating z score from log transformed FPKM values. To calculate z score, a data point was subtracted the mean and then divide the result by the standard deviation.

### Transcriptomic Profile Correlation Analysis

Transcriptomic profiles of never-transformed and pre-transform LUAD and transformed and *de novo* SCLC [Transcript Per Million (TPM) values from RNA-seq data] were obtained from published work by Quintanal-Villalonga et al. 2021^5^ and top 100 signature genes from each group vs. all others were determined using log-transformed TPM values with the eBays function in the limma package (3.56.2) in R (4.3.1). Log transformed TPM of signature genes were merged with *EED*-isogenic *RPR2* bulk tumor RNA-seq data, Pearson correlations were calculated with the cor function in base stats package in R (4.3.1). Correlation coefficients were further normalized by calculating z score as described above.

### Transcriptomic Data Analysis of the Cancer Genome Atlas (TCGA)

Batch normalized mRNA expression, mutation, and copy number alteration data from Lung adenocarcinoma (LUAD) patient tumors in the TCGA dataset were downloaded from cBioPortal (cbioportal.org)^68^. Samples with either mutation or deep deletion (copy number alteration value=-2) in *RB1* and/or *TP53* were categorized as a group and data was analyzed for all LUADs or only LUADs harboring EGFR mutations. Statistical testing between groups were performed using Tukey test to adjust for multiple comparisons.

### Gene Set Enrichment Analysis

GSEA software was downloaded from the Gene Set Enrichment Analysis website [http://www.broad.mit.edu/gsea/downloads.jsp]^70^. Pre-ranked GSEA was performed using hallmark gene sets from the human MSigDB Collections^32^ in Figs. 2D,3C,4M,S9L, top 100 neuroendocrine genes, ASCL1 target genes^59^ in Fig. 2D, human and mouse mucinous lung tumor signature^36^ in Fig. 2I, and ASCL1 correlated genes^59^, NEUROD1 correlated genes^59^, IMPOWER133-SCLC-N (NMF1) and SCLC-A (NMF2) gene signatures^12^ in Fig. 3E.

The Gene Set Variation Analysis (GSVA) method from the GSVA Bioconductor package (version 1.50.5), was used to calculate enrichment scores from tuft cell marker gene lists (Fig. S4B) and top 100 NE gene list (Fig. S4C) using normalized read counts from bulk *EED*-isogenic *RPR2* tumor RNA-seq data and from gene signatures of MAPK targets^67^, neuroendocrine markers^12^, tumor proliferation rate^12^, hallmark_MYC, hallmark_G2M, and neutrophil markers^12^ and human and mouse mucinous lung tumor signature (Fig. 6H-I) using normalized read counts from bulk RNA-seq data from *EGFR*-Mutant CRP and ERP GEMM lung tumors. GSVA was also performed using functional EED target genes on normalized read counts from IMPOWER133 NE and non-NE SCLC bulk RNA-seq data as previously reported^12^ (Fig. 3R,5E).

For gene set over-representation analysis, candidate gene sets were tested if over-represented against the human hallmark pathways from the MSigDB in Figs. 3Q, 5D, S8E, S9K,L, and S10I,L, mouse KEGG pathway collection^71^ in Figs. 5C,G,N, 7E,K, S7D,G, S9I, and S10K using R package Clusterprofiler (4.8.3). Mouse gene atlas^72^ (Fig. 2H), Gene Ontology Biological Processes^73^ (Fig. S10C,D), and ENCODE and ChEA consensus TFs^74^ (Figs. 2B, 7I, S7C,F, S10A,B) using R package enrichR (3.4).

### Reverse-Transcriptase Quantitative PCR (RT-qPCR)

RNA was extracted using Quick-RNA™ Miniprep kit (Cat#R1055, Zymo Research, CA, USA) according to the manufacturer’s instructions. RNA concentration was determined using the Nanodrop 8000 (Thermofisher Scientific). A cDNA library was synthesized using iScript Reverse Transcription Supermix for RT-qPCR (Biorad #1708841) according to the manufacturer’s instructions. qPCR were performed using the SsoAdvanced Universal SYBR Green Supermix (Biorad #1725271) according to the manufacturer’s instructions. The ΔΔC_T_ Method was used to analyze data. The C_T_ values for each primer set were then normalized to the C_T_ value of *36b4*. The data from Fig. 4C was then normalized to the control to determine the relative fold change in mRNA expression. The following probes were used for qPCR with SYBR Green: Mouse *Neurod1* Forward (5’-AGGCTCCAGGGTTATGAGATCG-3’), Mouse *Neurod1* Reverse (5’-TGAGAACTGAGACACTCATCTG-3’), Mouse *Met* Forward (5’-GACCTTAAGCGAGAGCACGA-3’), Mouse *Met* Reverse (5’-ATGCACTGTATTGCGTCGTC-3’), Mouse *36b4* Forward (5’- CTGTTGGCCAATAAGGTGCC-3’), Mouse *36b4* Reverse (5’- GTTCTGAGCTGGCACAGTGA-3’).

### Immunohistochemistry

Immunohistochemistry was performed on the Leica Bond III automated staining platform using the Leica Biosystems Refine Detection Kit (Leica; DS9800). FFPE tissue sections were baked for 30 minutes at 60°C and deparaffinized (Leica AR9222) prior to staining. Primary antibodies were incubated for 30 minutes, visualized via DAB, and counterstained with hematoxylin (Leica DS9800). The slides were rehydrated in graded alcohol and coverslipped using the HistoreCore Spectra CV mounting medium (Leica 3801733). 4 µm-thick sections were cut from formalin-fixed paraffin-embedded mouse tumor samples to perform single IHC studies using antibodies recognizing the following antigens: ASCL1 (rabbit monoclonal antibody clone EPR19840, Abcam; 1:100 concentration, antigen retrieval with citrate for 30 min.); NEUROD1 (rabbit monoclonal antibody clone EPR20766, Abcam; 1:100 concentration, antigen retrieval with citrate for 30 min.); POU2F3 (rabbit polyclonal antibody, Atlas Antibodies; 1:400 concentration, antigen retrieval with citrate for 30 min.); EED (rabbit monoclonal antibody clone E4L6E, Cell Signaling Technology; 1:200 concentration, antigen retrieval with EDTA for 30 min.); EGFR mutant-specific L858R antibody (rabbit monoclonal antibody clone 43B2, Cell Signaling Technology; 1:100 concentration, antigen retrieval with citrate for 30 min.); tri-methyl-histone H3 (Lys27) (rabbit monoclonal antibody clone C36B11, Cell Signaling Technology; 1:100 concentration, antigen retrieval with citrate for 30 min.); and Synaptophysin (rabbit monoclonal antibody clone SP11, Invitrogen; 1:50 concentration, antigen retrieval with citrate for 30 min.).

The immunostained slides were scanned at 40x magnification using the PhenoImager HT slide imager (Akoya Biosciences). For all markers, visual assessment of positive and negative cells was done by two independent pathologists (YNL, GRO). To quantify NEUROD1-positive tumor cells, tumor areas on each slide were identified and manually annotated by two pathologists (YNL, GRO) using the HALO image analysis platform (Indica Labs). The HALO multiplex-IHC v3.2.5 algorithm (Indica Labs) was then used to quantify the number of positive and negative tumor cells, and the percentage of NEUROD1-positive tumor cells was subsequently calculated.

### ATAC-sequencing

Nuclei isolation for ATAC-sequencing from mouse tumor tissue was performed using ATAC-Seq Kit (ActiveMotif, cat# 53150) following manufacture’s instruction. Briefly, 30mg frozen mouse *RPR2* lung tumor tissues were minced with a razor blade and homogenized in ATAC-seq Lysis Buffer in a dounce homogenizer. Lysates were then filtered through a 40 micron cell strainer and Trypan blue at 0.4% was used to count 100,000 nuclei which were subsequently rinsed with ice-cold PBS and resuspended in ice-cold ATAC-seq Lysis Buffer. After mixing with Tagmentation Master Mix, tagmentation reactions were incubated at 37 °C and shaken at 800 RPM for 30 minutes on a thermomixer. Transposed DNA was purified using columns and libraries were amplified following manufacture’s manual. 50 million paired-end reads, where each read is 150 base pairs long, were sequenced on a NextSeq instrument (Illumina).

### ATAC-seq Data Analysis

All samples were processed through the computational pipeline developed at the Dana-Farber Cancer Institute Center for Functional Cancer Epigenetics (CFCE) using primarily open-source program CHIPS^75,76^ . Sequence tags were aligned with Burrows-Wheeler Aligner (BWA)^77^ to build mm10 and uniquely mapped, non-redundant reads were retained. These reads were used to generate binding sites with Model-Based Analysis of ChIP-Seq 2 (MACS v2.1.1.20160309), with narrow peak option and a q-value (FDR) threshold of 0.01^78^. Average peak tracks were calculated with bigwigAverage function in deeptools (3.5.1) and visualized by IGV (v2.14.1)^79^. Peaks from all samples were processed using the DiffBind pipeline (v3.18)^80^ for differential binding analysis. Briefly, peaks were merged to create a union set of sites. Sequencing depth normalization was performed on each sample and DEseq2 was used to determine differential peaks. Log-transformed fold changes were subsequently shrunk with lfcshrunk for more accurate foldchange estimation. Differential binding peaks from each group were used for motif analysis by the motif search findMotifsGenome.pl in HOMER (v3.0.0)^81^, with cutoff q-value≤ 1e-10. The signals of each sample on differential binding sites were visualized by deeptools^82^. For principal component analysis, all peaks were used in the analysis and graphic visualization.

### Chromatin Immuno-Precipitation Sequencing (ChIP-seq) Analysis of *De Novo* SCLC and *EGFR*-Mutant LUAD Patient Derived Xenograft (PDX) Tumors

ChIP-seq for H3K27me3 and H3K4me3 from 4 NE *de novo* SCLC and 5 *EGFR*-Mutant LUAD PDX samples were previously published^52^ and the fastq files were obtained from the Gene Expression Ominibus (GEO accession#: GSE269746). High quality reads passing quality control using fastp were aligned to HumanG1Kv37 reference with BWA^83^. Uniquely mapped, non-redundant reads were generated by picard tools (2.19.0) and samtools (1.9) and subsequently used to generate binding sites with MACS2 (MACS v2.1.1.20160309) with q-value (FDR) threshold of 0.01 and narrow peak option for H3K4me3 and broad peak option for H3K27me3^78^. Peaks from all samples were processed using the DiffBind pipeline (v3.18)^80^ for differential binding analysis as described above.

### Mouse *EED*-Isogenic *RPR2* GEMM Tumors ChIP-seq Sample Preparation and Sequencing

Frozen mouse tumor tissue was dry pulverized in liquid nitrogen until it was finely powdered. Pulverized tissue was resuspended for crosslinking in 2 mM of disuccinimidyl glutarate (DSG, Pierce) for 45 minutes at room temperature. The crosslinked material was pelleted by centrifugation at 2500 RCF for 5 min and resuspended in 1 ml of 1% formaldehyde for 10 minutes. Formaldehyde was quenched with 0.125 M glycine for 5 minutes at room temperature and then the crosslinked material washed with PBS. Washed cells were now pelleted and resuspended in 500 μl of 1% SDS (50 mM Tris-HCl pH 8, 10 mM EDTA) and sonicated for 10 minutes using Covaris E220 sonicator at the following settings: 140 peak incident power, 5% duty factor and 200 cycles per burst) in 1 ml AFA fiber millitubes. Chromatin was immunoprecipitated with 5 μg of antibody (EED Cell Signaling E4L6E; H3K27ac Diagenode C15410196; H3K27me3 Active Motif 61018; H3K4me3 Abcam ab8580). 5 μg of chromatin was used for histone mark ChIPs, and 40 μg of chromatin was used for EED ChIPs. Imunoprecipitated Chromatin was pulled down using Protein A/G beads (Invitrogen Dynabeads 1000-2D and 1000-4D). Chromatin antibody complex immobilized on beads was washed 6X with RIPA buffer and eluted in 1% SDS and 0.1M Sodium bicarbonate. ChIP-ed DNA was crosslinked with Proteinase K and column purified. ChIP-seq libraries were made using Swift DNA Library Prep Kit (Swift Biosciences 10024). 75bp paired end reads were sequenced on a NextSeq instrument (Illumina).

### ChIP-seq Analysis and Peak Calling of Mouse *EED*-Isogenic *RPR2* GEMM Tumors

#### Peak calling and data analysis

All samples were processed through the computational pipeline developed at the Dana-Farber Cancer Institute Center for Functional Cancer Epigenetics (CFCE) using primarily open-source programs^75,76^. Sequence tags were aligned with Burrows-Wheeler Aligner (BWA)^83^ to build mm9 and uniquely mapped, non-redundant reads were retained. These reads were used to generate binding sites with Model-Based Analysis of ChIP-Seq 2 (MACS v2.1.1.20160309), with a q-value (FDR) threshold of 0.01^78^. We evaluated multiple quality control criteria based on alignment information and peak quality: (i) sequence quality score; (ii) uniquely mappable reads (reads that can only map to one location in the genome); (iii) uniquely mappable locations (locations that can only be mapped by at least one read); (iv) peak overlap with Velcro regions, a comprehensive set of locations – also called consensus signal artifact regions – in the genome that have anomalous, unstructured high signal or read counts in next-generation sequencing experiments independent of cell line and of type of experiment; (v) number of total peaks (the minimum required was 1,000); (vi) high-confidence peaks (the number of peaks that are tenfold enriched over background); (vii) percentage overlap with known DHS sites derived from the ENCODE Project (the minimum required to meet the threshold was 80%); and (viii) peak conservation (a measure of sequence similarity across species based on the hypothesis that conserved sequences are more likely to be functional).

#### Differential binding analyses

Peaks from all samples were merged to create a union set of sites for each transcription factor and histone mark using bedops^84^. Sample-sample correlation and differential peaks analysis were performed by the CoBRA pipeline^76^. Read densities were calculated for each peak for each sample and used for the comparison of cistromes across samples. Sample similarity was determined by hierarchical clustering using the Spearman correlation between samples. differential peaks were identified by DEseq2 with adjusted P ≤ 0.05. A total number of reads in each sample was applied to the size factor in DEseq2, which can normalize the sequencing depth between samples. Peaks from each group were used for motif analysis by the motif search findMotifsGenome.pl in HOMER (v3.0.0)^81^, with cutoff q-value ≤ 1e-10.

### Single Cell RNA-Sequencing Sample Preparation

The single-cell RNA sequencing (scRNA-seq) experiments in autochthonous CRISPR-based *EED*-WT or *EED*-Mutant *RPR2* GEMM models were performed as previously described^85^. Briefly, 3-4 month old male and female homozygous BL6J LSL-Cas9 mice (Jackson No. 026175) were intratracheally injected with *EED*-Mutant or *EED*-WT adenoviruses (see adenovirus method above). Once mice became symptomatic from their tumors (see method above), 6 independent *EED-*Mutant mice (4 *EED*-Mutant *RPR2* LUADs and 2 *EED*-Mutant *RPR2* SCLCs) and 3 independent *EED* WT mice were euthanized and lung tumors dissected and finely minced mechanically using a razor blade and then enzymatically digested with Mouse Tumor Dissociation Kit (Miltenyi Biotec, #130-096-730) following the manufacturer’s instruction. Briefly, minced tumor tissue was transferred to a gentleMACS C Tube containing enzyme mix prepared with 20% of Enzyme R option to preserve cell surface epitopes. Dissociation using the gentleMACS Octo Dissociator with Heaters (Miltenyi Biotec, #130-096-427) was performed using the 37C_m_TDK_2 gentleMACS Program. The single cell suspensions were resuspended in RPMI containing 10% FBS and subsequently passed through a 70 μm Cell Strainer (Greiner, #542070) and centrifuged at 300 x g for 3 min followed by 2 washes with 0.04% UltraPure Bovine Serum Albumin (Invitrogen, AM2616) in DPBS. Finally, dissociated cells were resuspended in DPBS with 0.04% UltraPure BSA and cell counts were measured with a Vi-CELL XR Cell Viability Analyzer (Beckman Coulter). Cells were then diluted in 0.04% BSA/DPBS at a cell concentration of 1000 cells/µL. About 16,000 cells were loaded onto a 10x Genomics Chromium^TM^ instrument (10x Genomics) according to the manufacturer’s instructions. The scRNA-seq libraries were processed using Chromium Next GEM Single Cell 5’ Kit v2 kit (10x Genomics). Quality controls for amplified cDNA libraries and final sequencing libraries were performed using Bioanalyzer High Sensitivity DNA Kit (Agilent). The sequencing libraries for scRNA-seq were normalized to 4 nM concentration and pooled. The pooled sequencing libraries were sequenced on Illumina NovaSeq S4 300 cycle platform. The sequencing parameters used were: Read 1 of 26bp, Read 2 of 90bp, Index 1 of 10bp and Index 2 of 10bp. *EED* WT RPR2 scRNA-seq data from 3 independent mice were previously published^85^.

### Single-Cell RNA Sequencing Data Analysis

Custom reference genome was established by adding *EGFP* and *Cas9* sequence^86^ to the mouse genome (mm10-2020-A). Cell ranger version 6.0.2 pipeline (10x Genomics) was used to align sequencing data to the custom mouse reference genome and generate the gene-level counts matrix. Unfiltered raw counts data was imported into Seurat v4 R package (version 4.1.0)^87^ for downstream processing and analysis. Low quality cells were filtered out using following thresholds: total UMI counts < 500, number of transcripts < 350, log_10_TranscriptsPerUMI <= 0.8, and cells with more than 15% transcripts mapping to mitochondrial genes. In addition, genes expressed in less than ten cells were removed. The UMI counts matrices were then natural-log normalized and scaled with Seurat’s ‘NormalizeData’ and ‘ScaleData’ functions.

### Single Nuclei RNA-Sequencing Sample Preparation

The single-nuclei RNA sequencing (snRNA-seq) sample preparation from *EED*-Mutant *RPR2* LUAD tumors were performed using Chromium Nuclei Isolation Kit with RNase Inhibitor (10x Genomics, PN-1000494) following manufacture’s instruction (10x Genomics, CG000124). Briefly, 30 mg snap-frozen tumor tissue from 3 independent *EED*-Mutant *RPR2* mice was dissociated in Lysis Buffer with pestle and centrifuged for 20s, in 16,000 rcf under 4 °C to a column to collect lysed nuclei in the flowthrough. Debris in the flowthrough containing nuclei was removed by Debris Removal Buffer before nuclei were washed 2 times with 5 min, 500 rcf, 4 °C spin. Nuclei was then counted using ViaStain AOPI Staining Solution (Revvity, CS2-0106-5ML) on a Nexcelom Cellometer K2 Fluorescent Cell Counter (Revvity, CMT-K2-MX-150) and resuspended in 1,000 nuclei/ul DPBS with 0.04% ultra-pure Bovine Serum Albumin (Invitrogen, AM2616). About 20,000 nuclei were loaded onto a ChromiumTM X instrument (10x Genomics) according to the manufacturer’s instructions. The snRNAseq libraries were processed using Chromium GEM-X Single Cell 5’ Kit v3 (10x Genomics). Quality controls for amplified cDNA libraries and final sequencing libraries were performed using Bioanalyzer High Sensitivity DNA Kit (Agilent). Libraries were subsequently quantified by Qubit fluorometer and Agilent 4200 TapeStation system. Library pooling and indexing was evaluated with shallow sequencing on an Illumina MiSeq. Subsequently, libraries were sequenced by the Molecular Biology Core Facilities at Dana-Farber Cancer Institute on an Illumina NovaSeq X Plus with the following sequencing parameters: 28bp, 10bp, 10bp, 90bp.

### Single-Nuclei RNA Sequencing Data Analysis

Custom reference genome was established by adding *EGFP* and *Cas9* sequence^86^ to the mouse genome (mm10-2020-A). Cell ranger version 9.0.0 pipeline (10x Genomics) was used to align sequencing data to the custom mouse reference genome and generate the gene-level counts matrix. Unfiltered raw counts data was imported into Seurat v4 R package (version 4.1.0)^87^ for downstream processing and analysis. Low quality cells were filtered out using following thresholds: total UMI counts < 500, number of transcripts < 375, log_10_TranscriptsPerUMI <= 0.8, mitochondrial gene ratio >= 0.2 for scRNA-seq and >= 0.1 for snRNA-seq. In addition, genes expressed in less than five cells were removed. The UMI counts matrices were then natural-log normalized and scaled with Seurat’s ‘NormalizeData’ and ‘ScaleData’ functions.

### Dimension Reduction, Cluster Analysis and Visualization of scRNA-Seq and snRNA-Seq Data

The Seurat v4 R package was used for dimension reduction and clustering. Top 2,000 genes with the highest variance were selected using the ‘FindVariableFeatures’ function with ‘vst’ method to perform linear dimensional reduction (principal component analysis) using the ‘RunPCA’ function, and top 40 principal components for scRNA-seq and top 25 principal components for snRNA-seq were used to perform graph-based unsupervised clustering with the ‘FindNeighbors’ and ‘FindClusters’ functions and Uniform Manifold Approximation and Projection for Dimension Reduction (UMAP) (<u>arXiv:1802.03426</u>) for data visualization in two-dimensional space. Automatic cluster annotation using the R package SingleR (version 1.8.1)^88^ with ImmGenData^89,90^ and manual tumor cluster annotation with markers of *EGFP* and *Cas9* were performed to select the tumor population. Subsequently, *Ptprc* positive tumor cells, *Cas9* and *EGFP* positive tumor microenvironment were filtered out. Then, the tumor subpopulations were reanalyzed. Seurat’s ‘CellCycleScoring’ function and S phase and G2/M phase specific gene lists were used to calculate the G2M and S phase gene expression scores and to be regressed out using the ‘var.to.regress’ option in the ‘SCTransform’ function with default parameters. Top 3,000 genes excluding mitochondrial and ribosomal genes with the highest variance were selected using the ‘FindVariableFeatures’ function with ‘vst’ method to perform linear dimensional reduction (principal component analysis) using the ‘RunPCA’ function, and top 40 principal components were used to plot UMAP for data visualization in two-dimensional space in scRNA-seq and top 9 principal components were used in snRNA-seq. Differential expression profile were obtained with Seurat’s ‘FindMarkers’ function using test.use = “wilcox” to compare differentially expressed genes.

Cell trajectory analysis was performed on tumor populations with the Monocle2 package (V2.24.0). For snRNA-seq, top 100 of each of differentially upregulated and downregulated genes between the *Neurod1*+ cluster and *Muc13*+ cluster were used as the ordering genes. For scRNA-seq, top 300 differentially expressed genes between *EED*-Mutant *RPR2* LUAD tumor cells and *EED*-WT *RPR2* SCLC tumor cells were used as ordering genes. To predict cell biological trajectories, Monocle2 estimated size factor and dispersion, followed by reduceDimension function with the “DDRTree” method to reduce the high-dimensional space as a way to infer cells progression through a given biological process. For snRNA-seq, top 300 non-mitochondrial genes that significantly correlates to the trajectory (based on descending order of q value from “differentialGeneTest” function) were plotted in the heatmap in Fig. 4H.

### Tumor and its Microenvironment Communication Analysis using Differential NicheNet

To study communication between the tumor and cells of the microenvironment in *EED*-WT SCLC, *EED*-Mutant SCLC, and *EED*-Mutant LUAD tumors, we used the nichenetr package V 1.1.0.39. NicheNet predicts which differentially expressed ligands from one tumor microenvironment population are most likely to affect target gene expression of tumor population and which specific target genes are affected by which of these predicted ligands. It uses curated ligand-receptor and ligand-target databases^42^ as references to evaluate the cellular communication networks between two cell types. Human gene names were converted to their one-to-one orthologs using ‘convert_human_to_mouse_symbols’ function. Ligands, receptors, and targets that expressed in at least 10% (default cutoff) of the single cells in a specific cluster were included. Differential genesets of interest were calculated using a log transformed fold change threshold of 0.15 (recommended cutoff for 10x datasets), and 2889 genes were included in *EED*-WT SCLC and 1668 genes of interest were included in *EED*-Mutant LUAD. Next, ligand-receptor-target links were prioritized using the weights of scaled properties of interest as follows: “scaled_ligand_score” = 5, “scaled_ligand_expression_scaled” = 1, “ligand_fraction” = 1, “scaled_ligand_score_spatial” = 0, “scaled_receptor_score” = 0.5, “scaled_receptor_expression_scaled” = 0.5, “receptor_fraction” = 1, “ligand_scaled_receptor_expression_fraction” = 1, “scaled_receptor_score_spatial” = 0, “scaled_activity” = 0, “scaled_activity_normalized” = 1, “bona_fide” = 1. Confident Tumor-TME interactions that are significantly enriched in *EED*-Mutant LUAD were further identified by significantly different expression of ligand in TME and receptor in tumors with Bonafide interaction. Tumor-expressing receptors in significantly enriched interaction pairs were further overlapped with significantly upregulated genes (p value<0.05) in mouse SCLC *RPR2* cell line 1014 after *EED* genetic inactivation. Scaled receptor score (Log2FoldChange comparing *EED*-Mutant LUAD and *EED*-WT SCLC) of unique receptors that are 1. Directly activated by loss of EED, 2, predicted activated by ligand from TME only in *EED*-Mutant LUAD, were plotted in Fig. 4Q.

### CNV Inference and Phylogenetic Tree Construction

R package inferCNV (1.16.0) was used to detect copy number variations from snRNA-seq data. Normal pulmonary epithelium and stroma cells were used as reference for the tumor population from the corresponding sample. The following parameters in the “run” function of infercnv were used: cutoff = 0.1, denoise = TRUE, cluster_by_groups = T, HMM = T. All genes from HMM copy number inferences (from file 17_HMM_predHMMi6.hmm_mode-samples.pred_cnv_genes.dat) were used to construct the phylogenetic tree; copy number was set to diploid if a cluster did not have an inferred CNV event predicted. Phylogenetic trees were inferred using the Maximum Parsimony (MP) method with the Subtree-Pruning-Regrafting (SPR) algorithm^91^ with search level 1 in MEGA11^92^.

### Flow Cytometry

Fresh single cell suspensions from *EED* WT or *EED*-Mutant *RPR2* mice tumors were prepared, washed twice in PBS, resuspended in FACS buffer (D-PBS containing 2% FBS) and transferred in 1.5 mL Eppendorf tubes. After washing, cells were incubated with fluorophore conjugated anti-H-2Ld/H-2Db (Biolegend #114507, clone 28-14-8) or the isotype control (Biolegend #400212, IgG2a, κ isotype Ctrl) at 1:100 dilution in the dark for 30 minutes at room temperature. After washing, cells were resuspended in FACS buffer, transferred to flow cytometry tubes containing a 40 μm filter and analyzed on a LSR Fortessa flow cytometer (Becton Dickinson, Franklin Lakes, NJ). Data analyses were performed with FlowJo software (V10).

### DNA Sequencing of GEMM Tumors

Genomic DNA from the tumors indicated was isolated using the QIAamp DNA Blood Mini Kit (Qiagen #51106) according to the manufacturer’s instructions. PCR was done by using KOD Xtreme™ Hot Start DNA Polymerase (Sigma-Aldrich, Cat# 71975) and following set of primers to amplify the genomic region of *Eed* targeted by sg*Eed* #2: forward *Eed #2* (5’- TACTGCTCACAGGACGAT -3’), reverse *Eed #2* (5’- GCTTCCAGCACATACCTT -3’). Two consecutive PCRs were used for *Rb1* where a 731 bp product was generated from the first PCR and subsequently used as template for the second PCR with the following sets of primers: 1^st^: forward *Rb1* (5’- CTATCATCTTCATGCTACAA -3’), reverse *Rb1* (5’- GCAGAATAAAATTCTACCAGG -3’), 2^nd^: forward *Rb1* (5’- GCATATATATCTACTTCAGCTG -3’), reverse *Rb1* (5’- GGTCAATGTGGAATACACAATTG -3’).The final PCR product was then column purified using Qiagen’s gel extraction kit and sequenced using next-generation Amplicon sequencing by the MGH DNA Core Facility or Sanger Sequencing (Azenta Life Sciences).

### EED cDNA for Exogenous EED Re-Expression

pTwist Lenti SFFV Puro WPRE lentiviral vectors encoding: 1) EED WT-HA or 2) EED Triple Mutant-HA (F97A/Y148A/Y365A) which encodes a functionally inactive EED Mutant-HA were codon optimized and synthesized by Twist Biosciences. All plasmids were sequence verified by sanger sequencing. The functionally inactive EED Mutant was confirmed as was unable to restore H3K27me3 upon re-expression relative to EED-WT which restored H3K27me3 upon re-expression.

### Statistics and Reproducibility

For all GSEA, enrichment analysis from RNA-sequencing, ATAC-sequencing and ChIP-sequencing data, statistical significance was calculated corrected for multiple hypothesis testing. For survival analysis in Fig. 1B, 6C, log-rank test was used. For correlation analysis in Fig. 2E, pearson correlation was calculated. For the GREAT analysis of ATAC-sequencing data in Figs. S8C,D, S10F,H, a binomial p-value was calculated as described previously^93^. For all other experiments, statistical significance was calculated using unpaired, two-sided Students t-test. *p*-values were considered statistically significant if the *p*-value was <0.05. Error bars represent S.E.M. unless otherwise indicated.

For all experiments with statistical data, the number of independent biological experiments are described in each figure legend. For immunoblot analyses in Figs. 3K,M, S6A,D-F, S7A, at least 3 biological independent experiments were performed and representative immunoblots are shown. For immunoblot analyses in Figs. 4K, S9C, immunoblots contains multiple independent tumors. For immunohistochemistry experiments and H&E staining, representative micrographs were shown from independent tumors from independent mice.

For the mouse experiments in Fig. 1, 48 mice were included and mice were completely randomized to receive sgEED *RPR2* or sgControl *RPR2* adenoviruses. For the mouse experiments in Fig. 6, 59 mice were included and mice were completely randomized to receive sgEED *RP* or sgControl *RP* adenoviruses. For all sequencing experiments (RNA-seq, ATAC-seq, scRNA-seq, snRNA-seq and ChIP-seq) the investigators that processed and sequenced the samples [DFCI’s CFCE core (ChIP-seq), DFCI MBCF core (snRNA-seq and ATAC-seq), DFCI TIGL core (scRNA-seq), or Novogene (RNA-seq)] were blinded to the identity of the samples. For all cell culture experiments, each experiment was repeated in at least 3 biological independent experiments as specified in the figure legend. No statistical methods were used to pre-determine sample sizes, but our sample sizes are similar to those reported in previous publications^25,26^. Data distribution was assumed to be a normal distribution, but this was not formally tested. For cell culture experiments, blinding was not possible. For all experiments, no data were excluded from any analyses.

