## Supplementary Data for "EED Maintains the Small Cell Lung Cancer Neuroendocrine Phenotype and Drives Lung Cancer Histological Transformation"

**Li et al. Supplementary Figures**

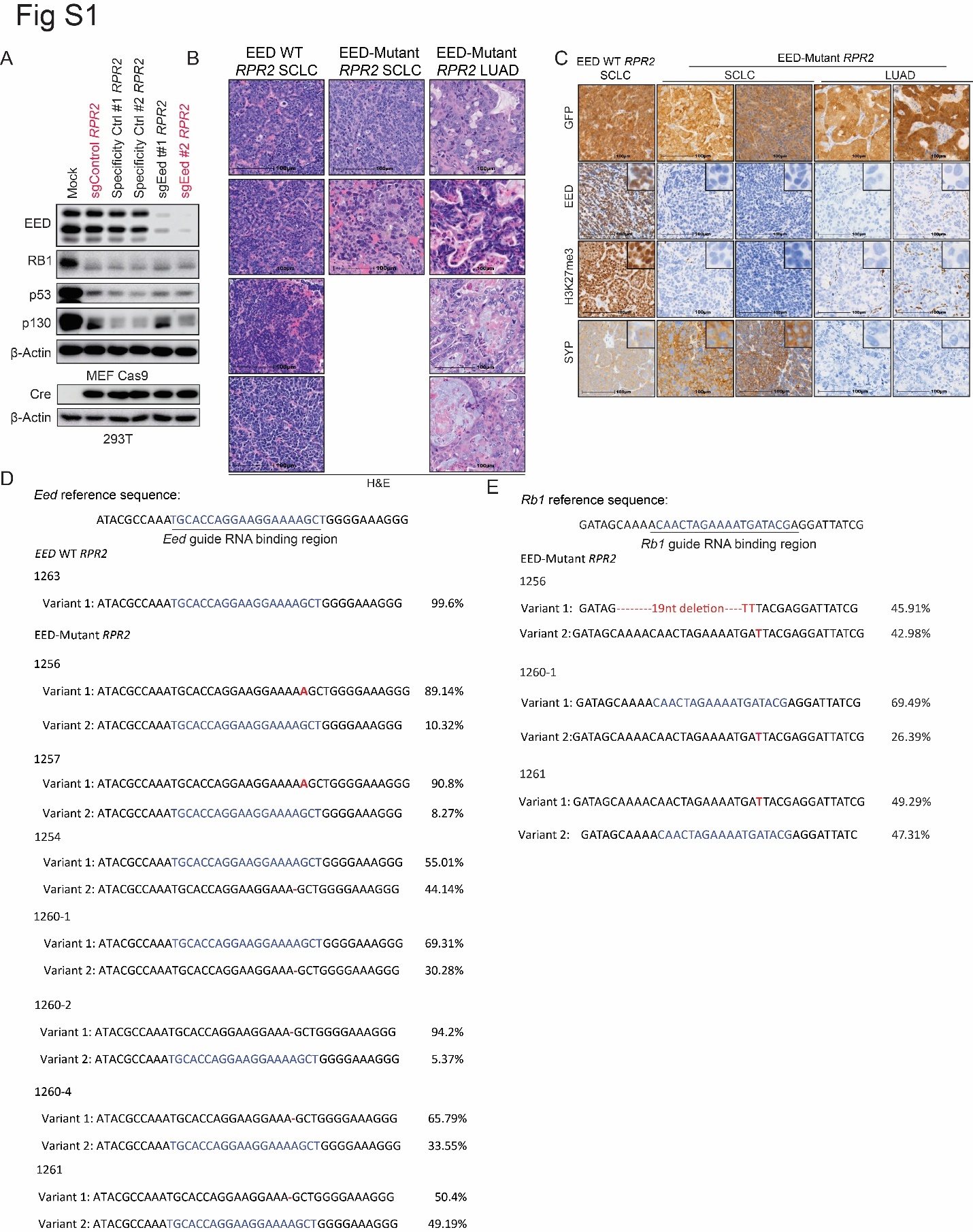

**Figure S1. EED Isogenic *RPR2* Adenovirus and SCLC GEMM Tumor Validation. Related to Figure 1.**

**A**. Immunoblot analysis of Cas9 expressing mouse embryonic fibroblast (MEF) cells and 293T cells infected with indicated *RPR2* adenoviruses. Adenoviruses in red were used for intratracheal injections in LSL-Cas9 mice. **B-C**. Representative H&E (**B**) and IHC for GFP, EED, H3K27me3, and Synaptophysin (SYP) (**C**) from additional lung tumors each from individual mice in the indicated groups (see also Fig. 1D-E). Scale bar=100 microns. Inserts are 9X magnification of the main graph. **D-E**. Sequence variants with percentages at the sgRNA binding region of endogenous *Eed* (**D**) and *Rb1* (**E**) from CRISPR amplicon sequencing of lung tumors from indicated genotypes. Variant nucleotides relative to the reference sequence are indicated in red text.

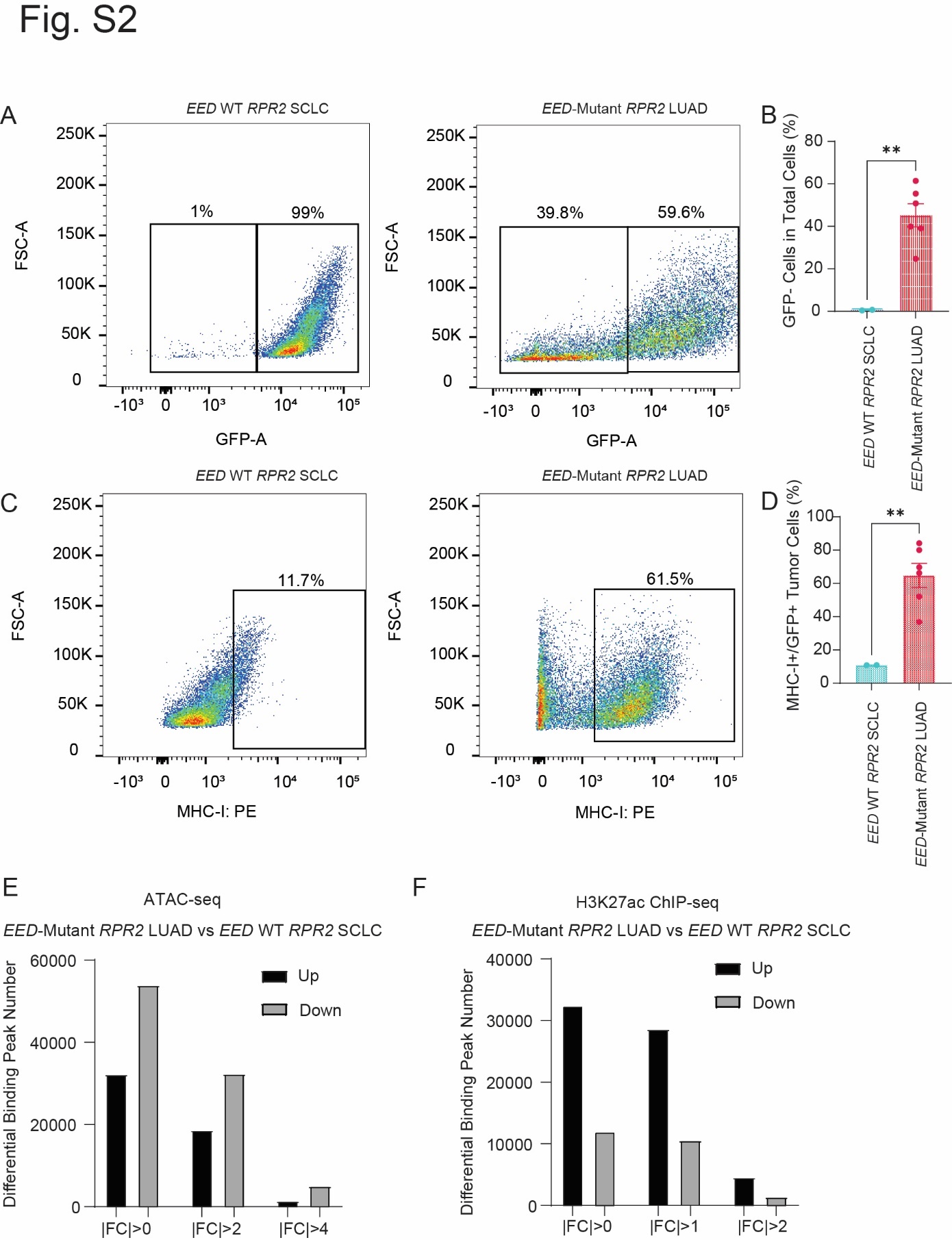

**Figure S2. *EED* Inactivation Epigenetically Promotes Lung Adenocarcinoma with Increased Restored MHC Class I Surface Expression and Increased Infiltration of Stromal and Immune Cells. Related to Figure 2.**

**A-B**. Representative scatter plots (**A**) and quantification (**B**) of flow cytometry analysis for GFP of *EED*-WT *RPR2* SCLC and *EED*-Mutant *RPR2* LUAD mouse tumors. GFP marks tumors cells and hence increased GFP negative cells in *EED*-Mutant *RPR2* LUAD mouse tumors indicates an increase in stromal/immune cells within tumors. **C-D**. Representative scatter plots (**C**) and quantification (**D**) of flow cytometry analysis for cell surface MHC class I on GFP-positive tumor cells. For B, D, n= 2 independent tumors for *EED*-WT *RPR2* SCLC and n=6 independent tumors for *EED*-Mutant *RPR2* LUAD. Two-sided p-values were calculated using a student’s t test. *=p<0.05, **=p<0.01, ***=p<0.001, ****=p<0.0001. **E-F**. Bar plots of differential binding peak numbers from ATAC-seq (**E**) and H3K27ac ChIP-seq (**F**) comparing *EED*-WT *RPR2* SCLC and *EED*-Mutant *RPR2* LUAD mouse tumors at the fold change (FC) thresholds indicated.

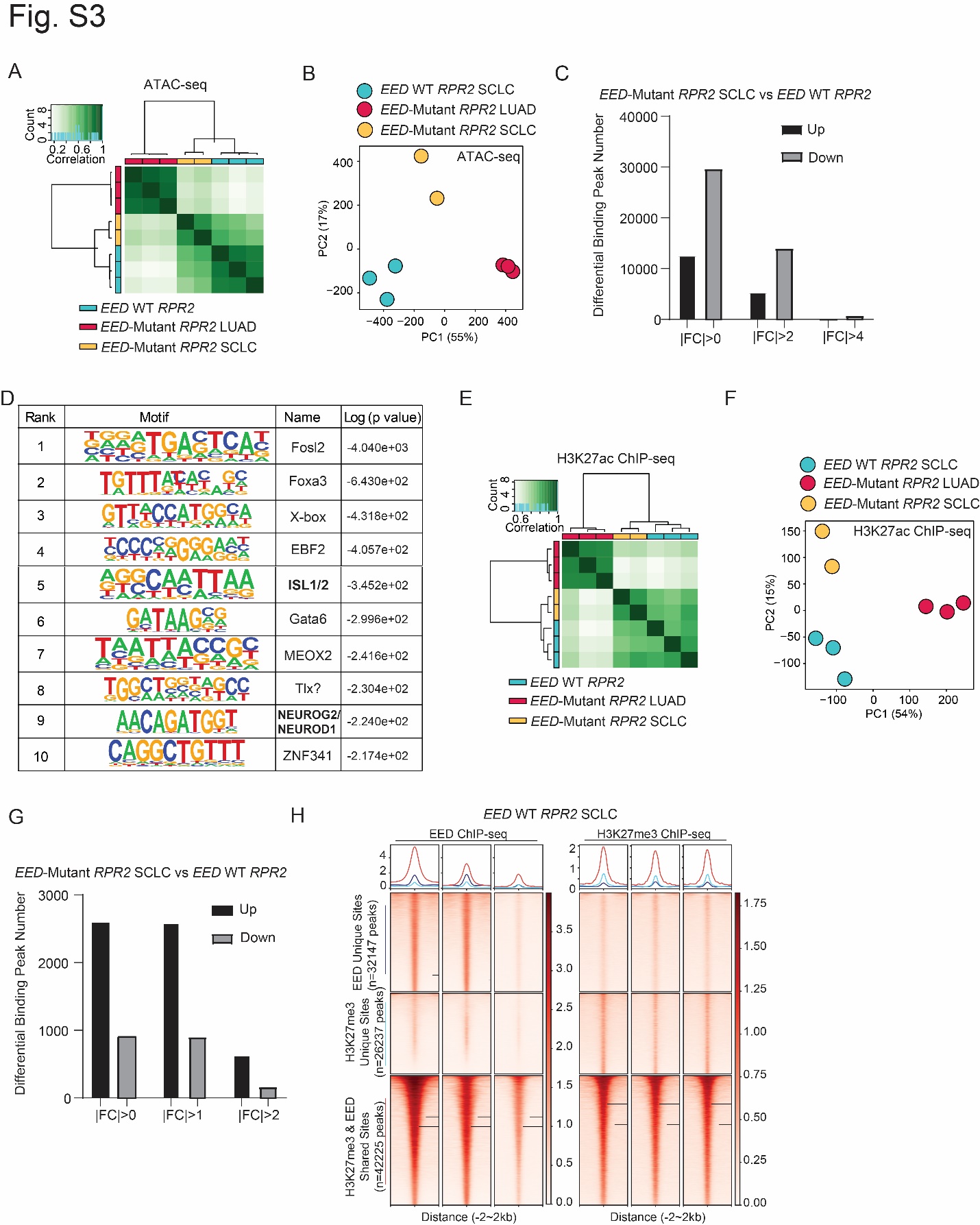

**Figure S3. Epigenetic Changes after *EED*-Mutant RPR2 SCLCs. Related to Figure 3.**

**A, B**. Correlation plot (**A**) and PCA plot (**B**) of all ATAC-seq data from *EED*-Mutant LUAD, *EED*-Mutant SCLC, and *EED*-WT SCLC lung tumors. **C**. Differential peak numbers of ATAC-seq comparing *EED*-Mutant SCLC vs *EED*-WT SCLC. **D**. HOMER motif enrichment analysis of top enriched motifs in *EED*-Mutant SCLC vs *EED*-WT SCLC. **E,F**. Correlation plot (**E**) and PCA plot (**F**) all H3K27Ac ChIP-seq data from *EED*-Mutant LUAD, *EED*-Mutant SCLC, and *EED*-WT SCLC lung tumors. **G**. Differential peak numbers of H3K27Ac ChIP-seq comparing *EED*-Mutant SCLC vs *EED*-WT SCLC. **H**. Heatmaps and profile plots of shared binding peaks of EED and H3K27me3 (red) and unique peaks to EED (purple) and H3K27me3 (blue). White is low, red is high. FC=Fold Change. For correlation plots, green is high, white is low.

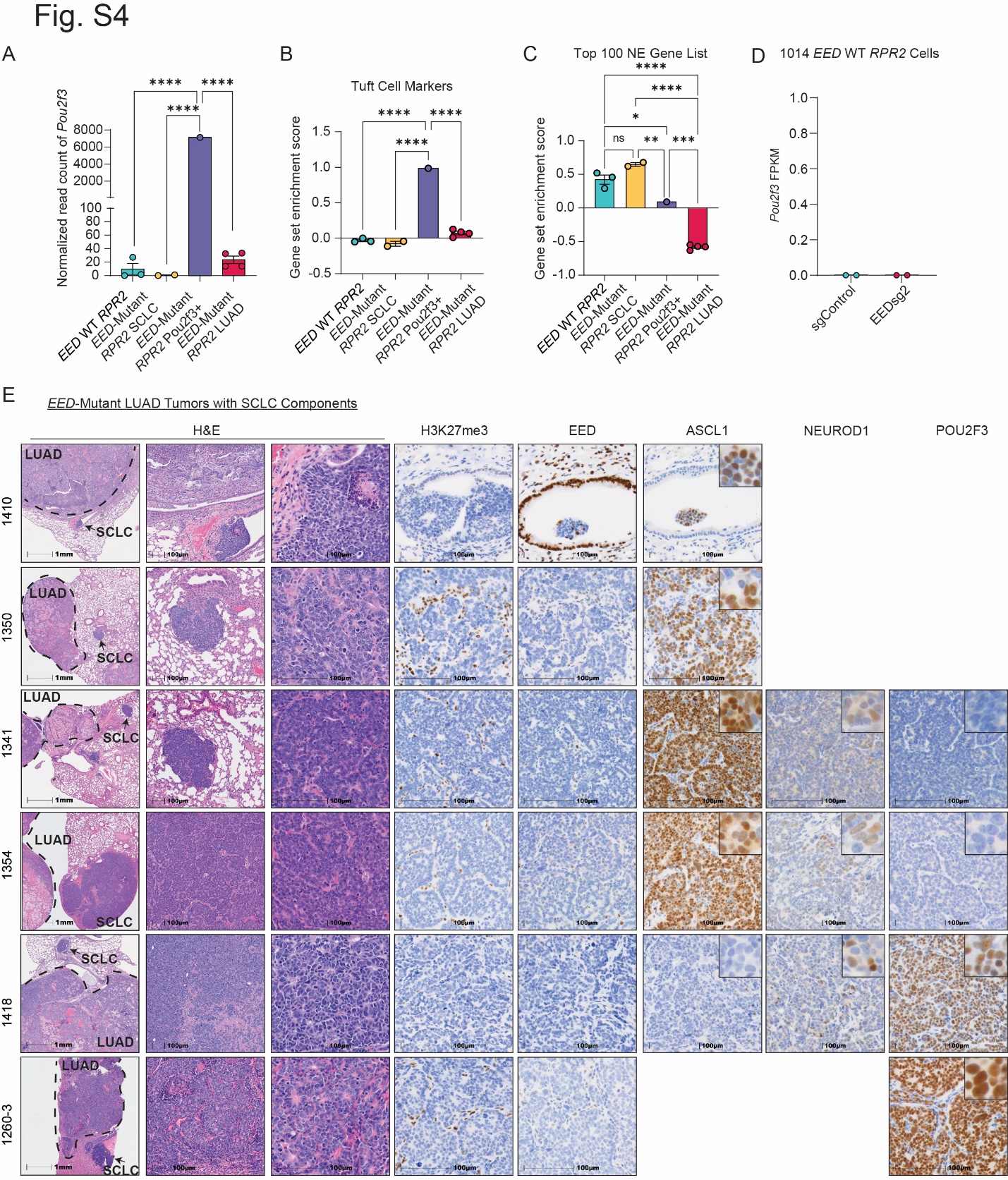

**Figure S4. Analysis of SCLC Components in *EED*-Mutant *RPR2* LUAD Reveals Loss of EED and Heterogenous Expression of SCLC Lineage Transcription Factors. Related to Figure 4.**

**A-C**. Bar plots of normalized read counts of *Pou2f3* (**A**), gene set enrichment scores of tuft cell markers (**B**), and top 100 neuroendocrine genes from RNA-seq data from Fig. 2-3 from the GEMM lung tumors indicated. Student’s t-test was used to calculated two-sided p values. *=p<0.05, **=p<0.01, ***=p<0.001, ****=p<0.0001. **D**. Dot plot of Pou2f3 FPKM values from RNA-seq data (see Fig. 3L) from 1014 *RPR2* EED-WT SCLC cells transduced with an sgRNA targeting *EED* (sg*Eed*) or a non-targeting sgRNA as a control (sgControl). **E**. Representative H&E and IHC staining for H3K27me3, EED, ASCL1, NEUROD1, and POU2F3 from *EED*-Mutant *RPR2* LUAD lung tumors that also contained a SCLC component shown in Fig. 1C. Scale bar (1^st^ column) =1 mm. Scale bar (2^nd^-8^th^ columns) =100 microns. Insets are 9X magnification of the corresponding main image. Blank slots are present because SCLC tumor nodule was lost when sectioning paraffin block. Dotted line in 1^st^ column indicates LUAD or SCLC compartment.

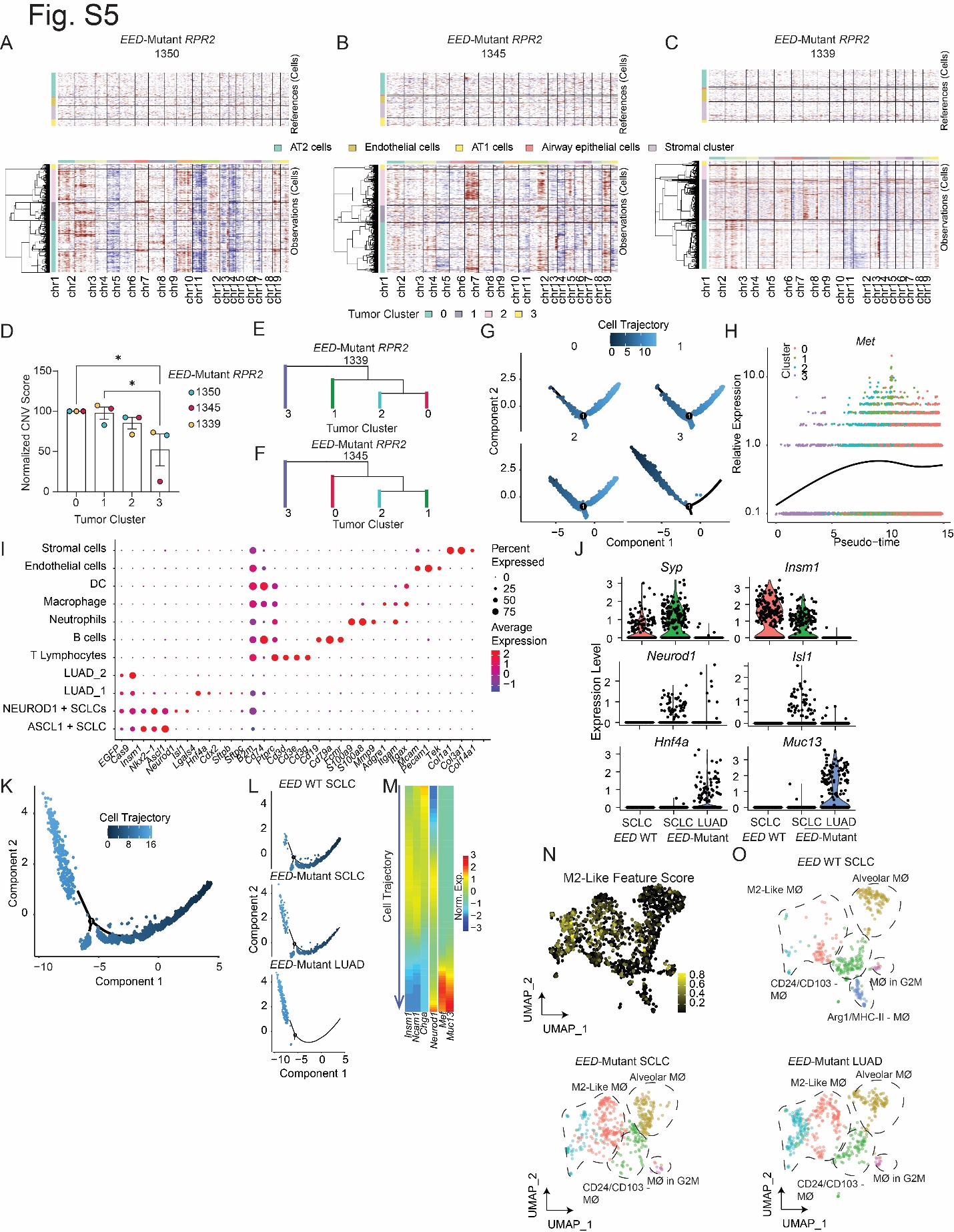

**Figure S5. EED Inactivation Promotes SCLC-to-LUAD Transformation Through a NEUROD1-Positive Intermediate Cell State and Requires Cues from the Tumor Immune Microenvironment *In Vivo.* Related to Figure 4.**

**A-C**. The landscape of copy number variations (CNVs) from snRNA-seq from each *EED*-Mutant *RPR2* LUAD tumor indicated. Cells within each tumor cluster are grouped by hierarchical clustering based on predicted CNVs. Pulmonary epithelium/stroma cells within each tumor are are used as a normal reference. For reference cells, n=1037 in *EED*-Mutant *RPR2* LUAD (1350 tumor), n=2529 in *EED*-Mutant *RPR2* LUAD (1345 tumor), n=904 in *EED*-Mutant *RPR2* LUAD (1339 tumor). For tumor cells: n=3256 in *EED*-Mutant *RPR2* LUAD (1350), n=5015 in *EED*-Mutant *RPR2* LUAD (1345), n=8319 in *EED*-Mutant *RPR2* LUAD (1339). **D**. Quantification of CNV events (normalized CNV score) from each tumor clusters in each *EED*-Mutant *RPR2* LUAD sample. Two-sided p-values were calculated using a student’s t-test. *=p<0.05, **=p<0.01, ***=p<0.001, ****=p<0.0001. **E,F**. Phylogenetic trees of tumor clusters using Maximum Parsimony method from *EED*-Mutant *RPR2* LUAD tumors 1339 (**E**) and 1345 (**F**). **G**. Pseudo-time cell trajectory projection of each separate tumor cell clusters. Black=early, blue=late. **H**. Scatter plot of *Met* relative expression in pseudo-time with each tumor cell cluster indicated. Curve indicates average expression. **I**. Expression dot plot of genes defining cell types in the scRNA-seq analysis. Expression level is visualized as dot color. Blue is low, red is high. Percent of cells expressing the indicated marker is visualized by the size of the dot. **J**. Violin plots of marker gene expression from scRNA-seq data in each sample group indicated. **K,L**. Pseudo-time tumor cell trajectory projection of 3 independent *EED*-WT SCLC, 2 independent *EED*-Mutant SCLC, and 4 independent *EED*-Mutant LUAD tumors in all samples (**K**) and each group separated (**L**). Black=early, blue=late. **M**. Heatmap of normalized expression level of marker genes along the tumor cell trajectory. Blue is low, red is high. **N,O**. UMAP of Mϕ cells from scRNA-seq with color gradients indicating expression level of M2-like feature genes (**N**) and subcluster labeling in indicated sample groups (**O**). Black is low, yellow is high. Mϕ, macrophage.

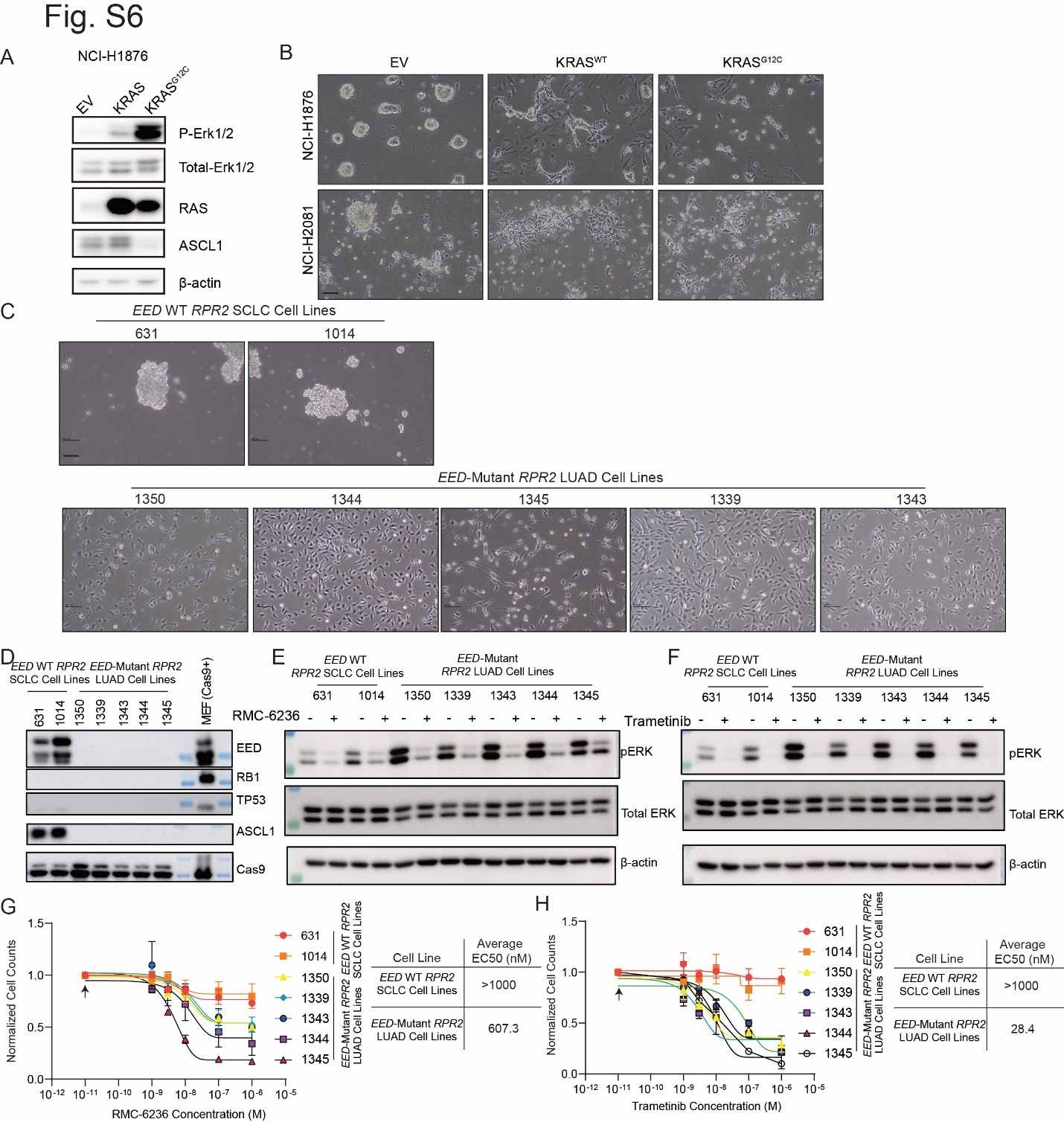

**Figure S6. *EED*-Mutant *RPR2* LUAD Cell Lines are Relatively Hypersensitivity to RAS and MAPK Inhibition and Oncogenic RAS signaling promotes a non-NE Phenotype in SCLC. Related to Figure 5.**

**A**. Immunoblot analysis of the human NCI-H1876 SCLC cell line lentivirally transduced to overexpress KRAS^WT^, constitutively activated KRAS^G12C^, or the corresponding empty vector (EV). **B**. Representative micrographs of NCI-H1876 and NCI-H2081 cells overexpressing the indicated constructs. Scale bar=50 microns. **C**. Representative micrographs of cell lines made from *EED*-WT *RPR2* lung tumors (631, 1014) and cell lines made from *EED*-Mutant *RPR2* LUAD tumors (1350, 1344, 1345, 1339, and 1343). Scale bar=100 microns. **D**. Immunoblot analysis of cell lines in C confirming loss of EED and ASCL1 in *EED*-Mutant *RPR2* LUAD tumors and loss of RB1 and TP53 in all cell lines. Cas9 expressing mouse embryonic fibroblasts (MEFs) were used as control. **E-F**. Immunoblot analysis of cell lines from C treated overnight at 1 micromolar with RAS-ON inhibitor RMC-6236 (**E**) and MEK inhibitor trametinib (**F**). **G,H.** Normalized dose response curves with EC50 indicated of each cell line from C treated with RMC-6235 (**G**) or trametinib (**H**). If 50% killing were not reached at 1000 nM, 1000 nM was used in calculating the average EC50. Arrow in G, H indicated DMSO-treated sample used for normalization.

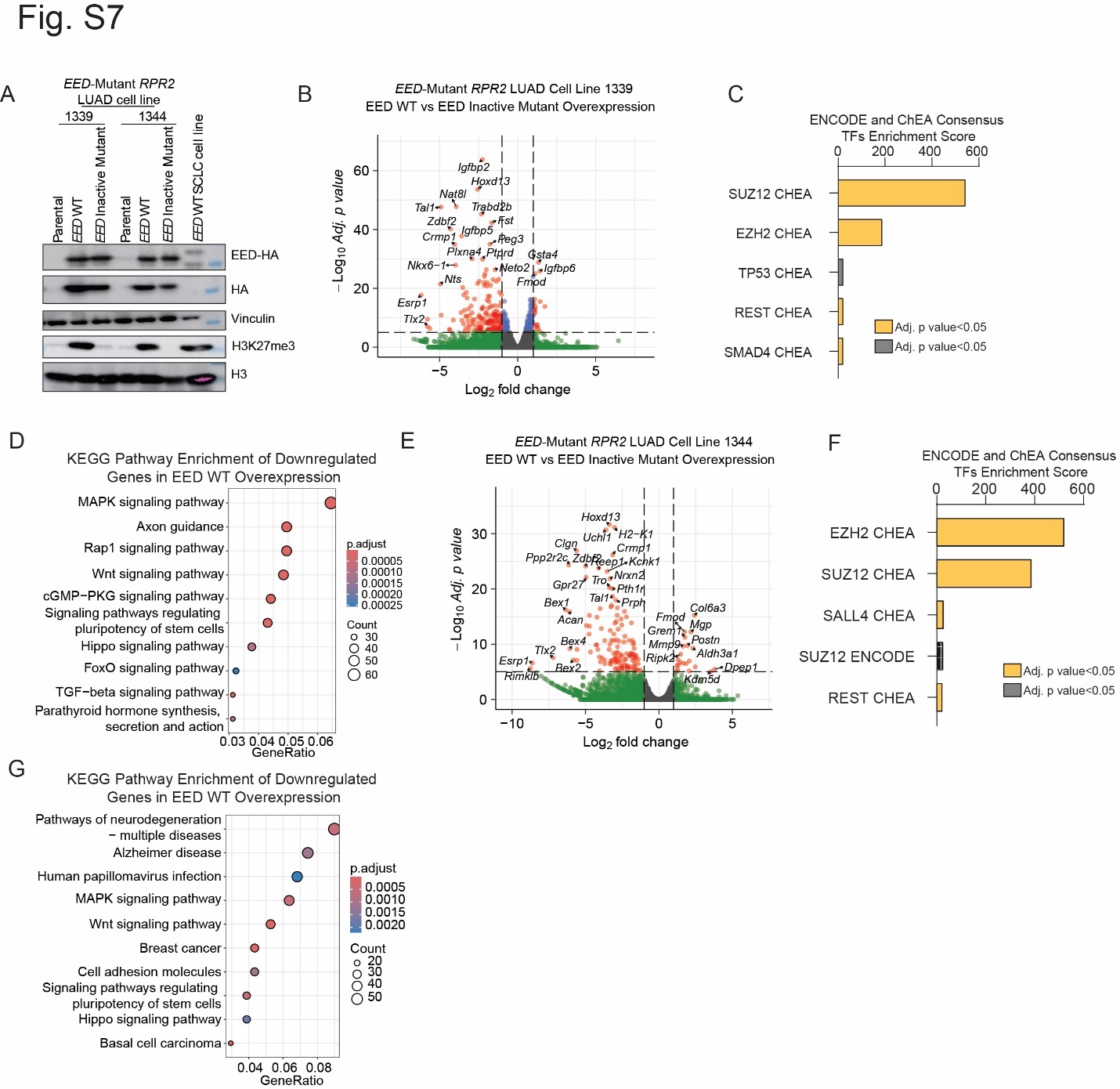

**Figure S7. EED Overexpression Represses Oncogenic RAS Signaling. Related to Figure 5.**

**A**. Immunoblots of *EED*-Mutant *RPR2* LUAD cell lines (1339 and 1344) lentivirally transduced to re-express EED WT or an EED catalytic dead mutant (EED inactive mutant) which fails to restore H3K27me3 relative to EED-WT. EED WT SCLC *RPR2* cell line (1014 cells) are included as a benchmark control. **B,E**. Volcano plots of RNA-seq data comparing EED WT vs. EED inactive mutant in the *EED*-Mutant *RPR2* LUAD cell lines 1339 (**B**) and 1344 (**E**). n=2 biological independent experiments for each condition for each cell line. **C,F**. Bar plot of ENCODE and ChEA Consensus TFs enrichment score from top enriched transcription factors whose targets were enriched in the differentially downregulated genes in RNA-seq data from B, E comparing EED WT vs EED inactive mutant in *EED*-Mutant *RPR2* LUAD cell lines 1339 (**C**) and 1344 (**F**). Yellow: significantly enriched with adjusted p-value<0.05, grey: adjusted p-value<0.05. **D,G**. Dot plots of KEGG pathway enrichment analysis of significantly downregulated genes in EED WT vs EED inactive mutant in *EED*-Mutant *RPR2* LUAD cell lines 1339 (**D**) and 1344 (**G**). BH adjusted p-values were visualized by the dot color, red=low, blue=high. Enriched gene counts were visualized by the size of the dots. Gene ratio=enriched gene number/total differentially downregulated genes.

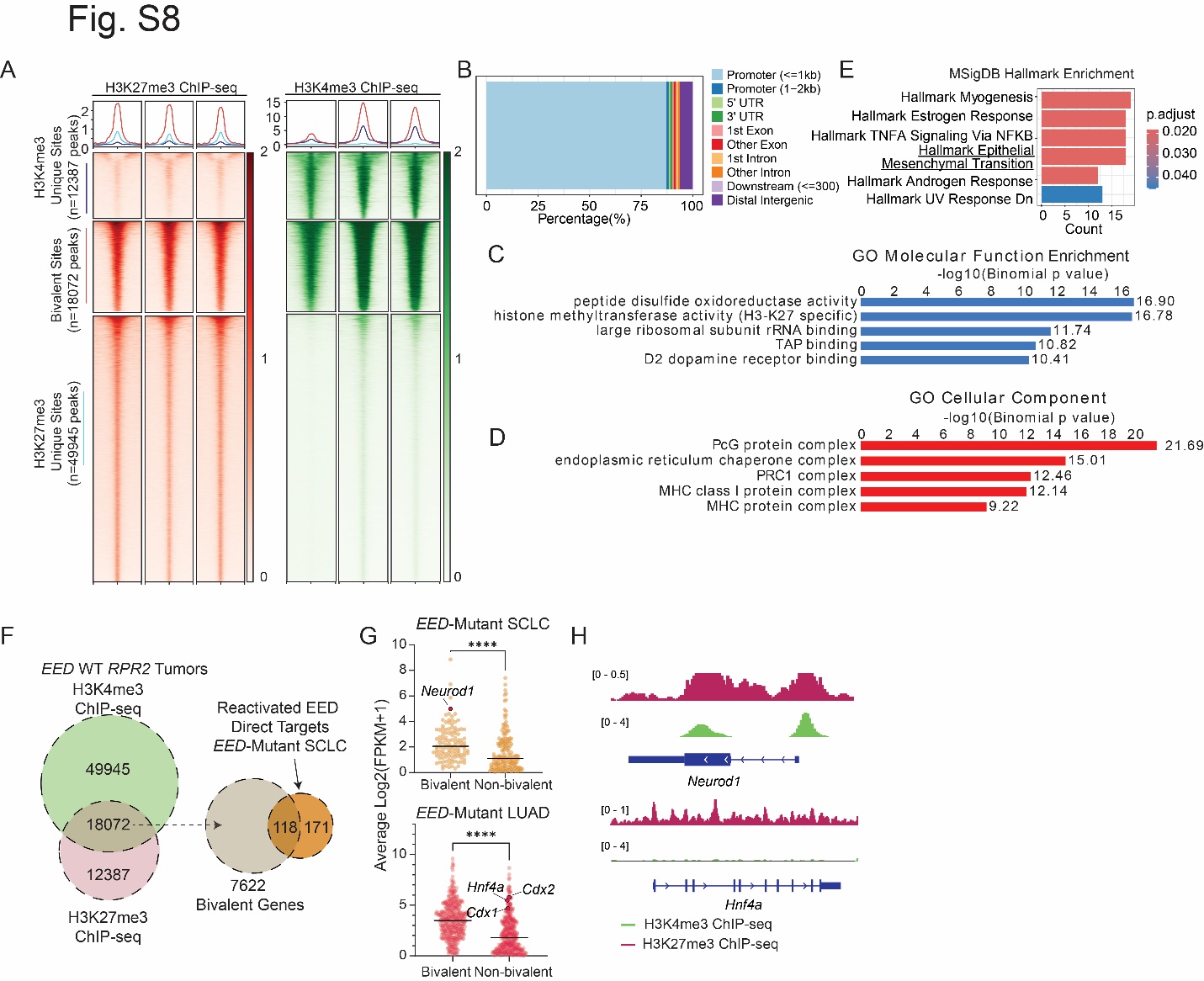

**Figure S8. Loss of the PRC2 Complex Initiates SCLC to LUAD Histological Transformation in Murine Tumors Through Epigenetic Activation of Bivalent Genes Marked by both H3K27me3 and H3K4me3. Related to Figure 5.**

**A**. Heatmaps and profile plots of shared binding peaks of H3K27me3 and H3K4me3 (red) and unique peaks to H3K4me3 (purple) and H3K27me3 (blue) in murine *EED*-WT *RPR2* SCLC lung tumors. **B**. Genomic distribution of bivalent peaks around the gene body. **C,D**. Genomic Regions Enrichment of Annotations Tool (GREAT) analysis of bivalent peaks for gene ontology molecular function enrichment (**C**) and cellular component enrichment (**D**). Binomial p-values are indicated. **E.** Bar plot of hallmark pathway enrichment of genes with bivalent peaks in its promoter region (absolute distance <2 kb from TSS) in *EED*-WT *RPR2* SCLC. BH adjusted p-values were visualized by the bar color, red=low, blue=high. Count: Enriched gene counts. **F**. Schematic showing strategy to identify bivalent genes reactivated upon EED loss by overlapping peaks of H3K4me3 and H3K27me3 ChIP-seq data from *EED*-WT *RPR2* SCLC tumors (left) and then overlap these bivalent peaks with reactivated EED direct target genes in *EED*-Mutant *RPR2* SCLC in Fig. 5C (right). **G**. Expression of reactivated EED direct targets that are bivalent or not in *EED*-Mutant SCLC. **H**. Tracks of averaged H3K27me3 (green) and H3K4me3 (red) ChIP-seq at *Neurod1* (top) and *Hnf4a* (bottom) showing *Neurod1*, but not *Hnf4a* is marked with bivalent peaks.

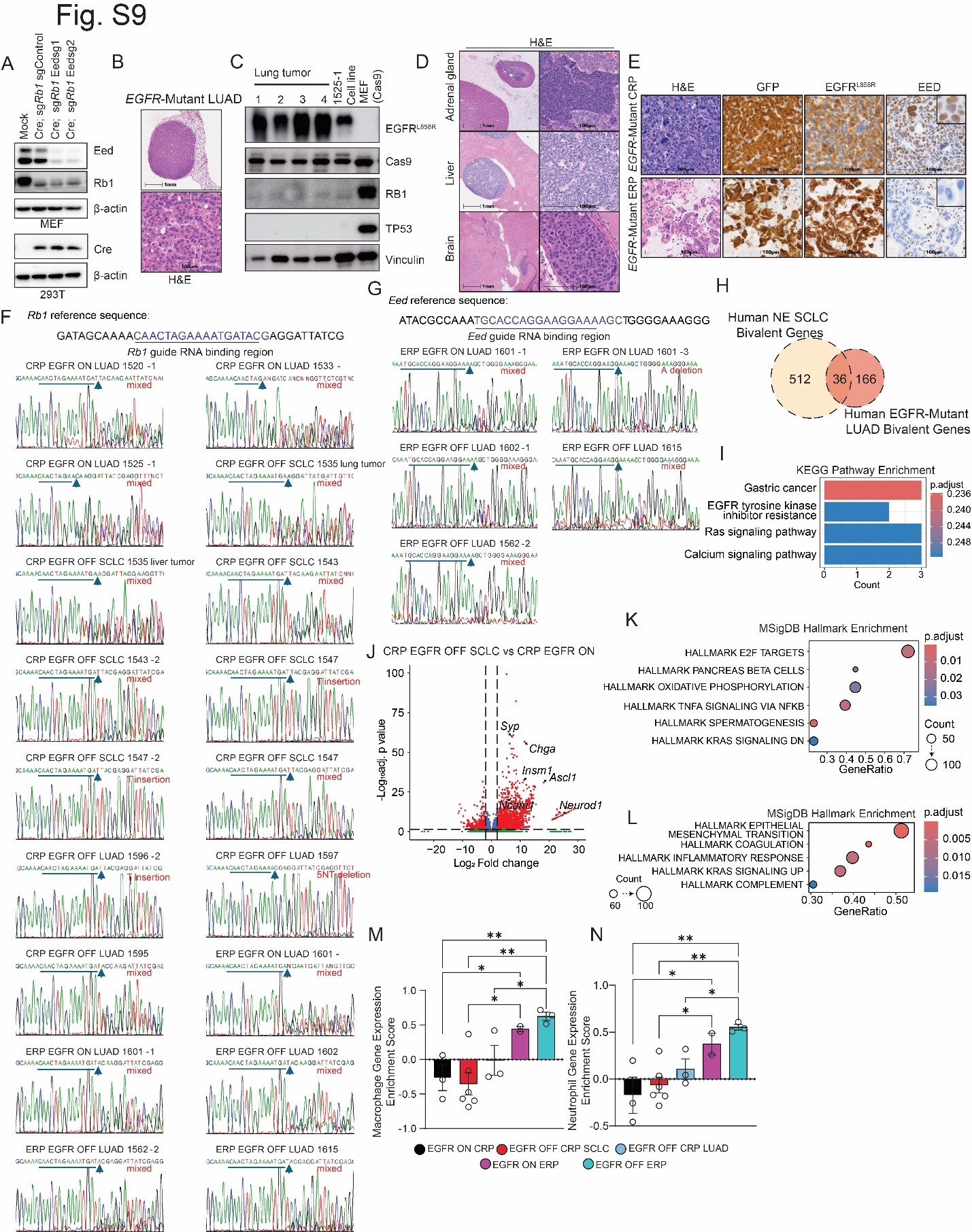

**Figure S9. EED Inactivation Blocks LUAD to SCLC Transformation as a Mechanism of Resistance to EGFR Inactivation in an *EGFR*-Mutant GEMM. Related to Figure 6.**

**A.** Immunoblot analysis of Cas9 expressing mouse embryonic fibroblast (MEF) cells and 293T cells transduced with indicated adenoviruses. **B**. Representative H&E of a representative *EGFR*-Mutant lung tumor transduced with CMV-cre;sgControl;sg*Rb1* adenovirus and treated with Dox (Doxycycline) until the mouse became symptomatic from tumor burden. Scale bar (top =1 mm, scale bar (bottom)=100 microns. **C**. Immunoblot analysis of lung tumors and a derived cell line from *EGFR*-Mutant sgControl sgRb1 Trp53-/- (CRP) mice treated with Dox until mice became clinically symptomatic from lung tumor burden in Dox ON mouse. Cas9 expressing MEFs were used as control. **D**. Representative H&E and IHC staining for GFP, EGFR^L858R^, and EED from lung tumors from *EGFR*-Mutant CRP and ERP mice treated with Dox. Scale bar=100 microns. Insets are 9X magnification. **E**. Representative H&E images of distant metastasis in adrenal gland, liver, and brain from *EGFR*-Mutant sgControl, sg*Rb1,* *Trp53*-/- (CRP) with tumor recurrence after Dox withdrawal according to the treatment strategy in **Fig. 6B**. Scale bar (left) = 1 mm, scale bar (right)=100 microns. **F,G**. Chromatograms from Sanger sequencing after PCR amplification of the DNA sequence around *Rb1* (**F**) and *Eed* (**G**) guide RNA binding regions. Blue arrows indicate sgRNA cut site. Red text indicates type of CRISPR-mediated edits. **H,I**. Schematic (**H**) and KEGG pathway enrichment of overlapping bivalent genes in human NE SCLC and *EGFR*-Mutant LUAD PDX tumors (**I**) from **Fig. 6G,K**. BH adjusted p-values were visualized by the bar color, red=low, blue=high. Count: Enriched gene counts. **J**. Volcano plot of RNA-seq data from Fig. 7G of EGFR Dox-OFF CRP tumors with SCLC histological transformation (HT) vs. EGFR Dox-ON CRP tumors. Vertical lines, fold change=2. Horizontal line, adjusted p value=0.05. **K**. Hallmark pathway enrichment analysis of differentially expressed genes in EGFR Dox-OFF CRP tumors with SCLC transformation vs. EGFR Dox-ON CRP tumors. **L**. Hallmark pathway enrichment analysis of differentially expressed genes in EGFR Dox-OFF CRP tumors that recurred as LUAD vs. EGFR Dox-ON CRP tumors. BH adjusted p-values were visualized by the bar color, red=low, blue=high. Count: Enriched gene counts. **M,N**. Bar plots of gene set enrichment score of tumor associated macrophage gene signature (**M**) and neutrophil gene signature (**N**). Student’s t-test was used to calculate two-sided p-values. *=p<0.05, **=p<0.01, ***=p<0.001, ****=p<0.0001. CRP: sgControl; sg*Rb1*; *Trp53*-/-, ERP: sg*Eed*; sg*Rb1*; *Trp53*-/-.

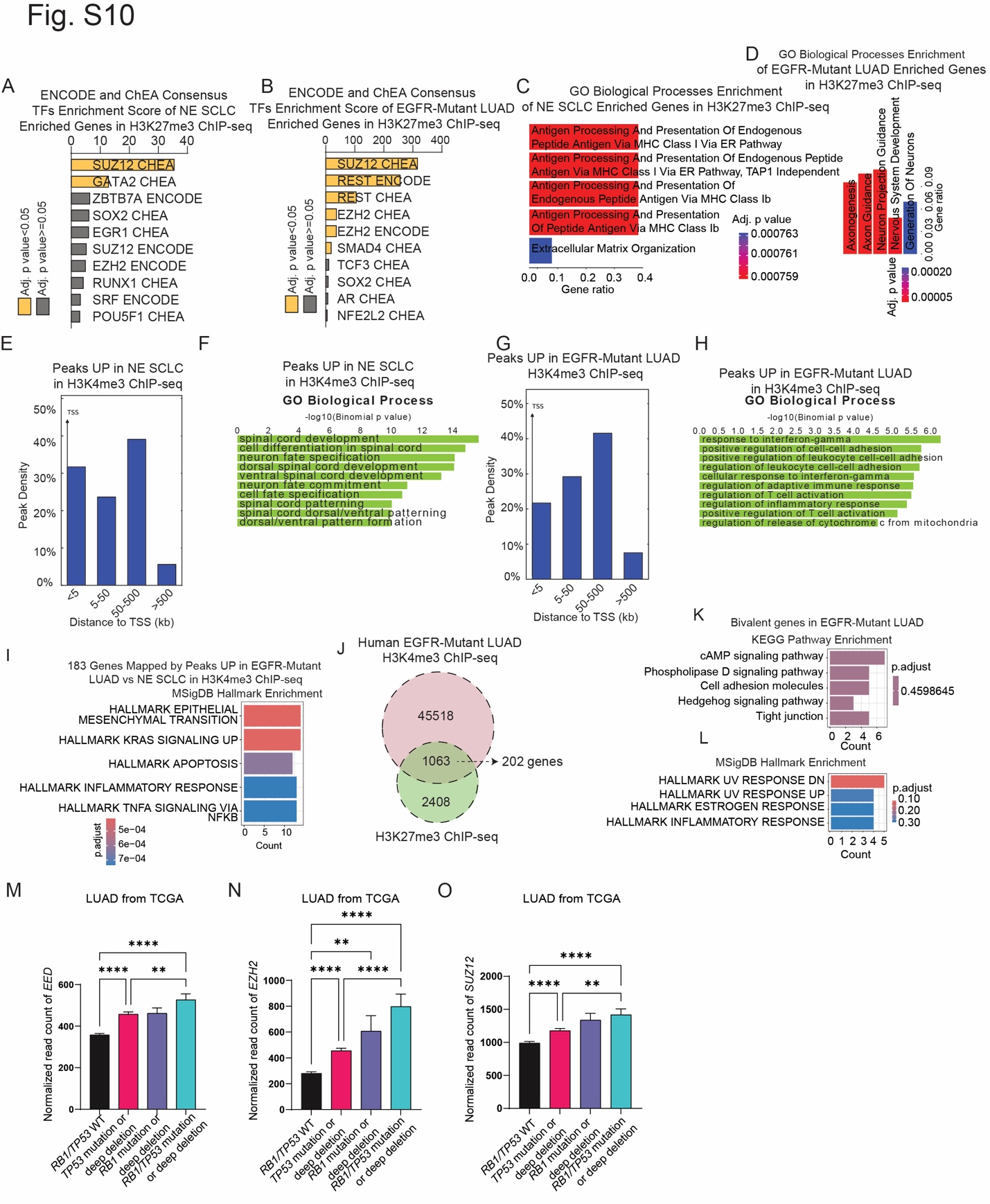

**Figure S10. Human SCLC Blocked LUAD Oncogenic RAS/MAPK/PI3K-Akt Signaling Through Epigenetically Poised Activation of Bivalent Genes Marked by both H3K27me3 and H3K4me3. Related to Figure 7.**

**A,B**. Bar plot of ENCODE and ChEA Consensus TFs enrichment score from top enriched transcription factors in human NE SCLC (**A**) and *EGFR*-Mutant LUAD (**B**) PDX tumors whose target genes have significantly gained H3K27me3 peaks. Yellow: significantly enriched with adjusted p-value<0.05, grey: adjusted p value<0.05. **C,D**. Bar plot of gene ontology biological processes enrichment of genes that have significantly gained H3K27me3 peaks in human NE SCLC (**C**) and *EGFR*-Mutant LUAD (**D**). BH adjusted p-values were visualized by the bar color, red=low, blue=high. Gene ratio=enriched gene number/total number of genes that have significantly gained H3K27me3 peaks in human NE SCLC (**C**) and *EGFR*-Mutant LUAD (**D**). **E,G**. H3K4me3 peak density bar plots of absolute distance to the TSS of significant peaks in human NE SCLC (**E**) and *EGFR*-Mutant LUAD (**G**) PDX tumors. TSS=transcription start site. **F,H**. GREAT analysis for gene ontology biological process enrichment of H3K4me3 peaks upregulated in NE SCLC (**F**) and *EGFR*-Mutant LUAD (**H**). Binomial p-values are indicated. **I**. Bar plot of hallmark pathway enrichment of genes with H3K4me3 peaks upregulated in *EGFR*-Mutant LUAD vs. NE SCLC human PDX tumors. BH-adjusted p-values are visualized by the bar color, red=low, blue=high. Count: Enriched gene counts. O. Schematic to identify bivalent genes in human *EGFR*-Mutant LUAD. **J-L**. Schematic of identifying bivalent genes in human EGFR-Mutant LUAD by overlapping H3K27me3 and H3K4me3 ChIP-seq (**J**), Bar plots of enriched KEGG pathways (**K**) and hallmark pathways (**L**) of bivalent genes in human *EGFR*-Mutant LUAD PDX tumors. BH adjusted p-values are visualized by the bar color, red=low, blue=high. Count: Enriched gene counts. **M-O.** Bar plots of RNA-seq batch normalized expression of *EED* (**M**), *EZH2* (**N**), *SUZ12* (**O**) from human LUAD tumors with *RB1* and *TP53* genetic alterations indicated in the Cancer Genome Atlas (TCGA)^68^. *RB1/TP53* WT: n=241 tumors, *TP53* mutation or deep deletion: n=152 tumors, *RB1* mutation or deep deletion: n=5 tumors, *RB1/TP53* mutation or deep deletion: n=33 tumors. One way ANOVA with Tukey test was used to calculate two-sided p values adjusted for multiple comparisons. *=p<0.05, **=p<0.01, ***=p<0.001, ****=p<0.0001.
